# Use of signals of positive and negative selection to distinguish cancer genes and passenger genes

**DOI:** 10.1101/2020.06.04.133199

**Authors:** László Bányai, Mária Trexler, Krisztina Kerekes, Orsolya Csuka, László Patthy

## Abstract

A major goal of cancer genomics is to identify all genes that play critical roles in carcinogenesis. Most approaches focused on genes that are positively selected for mutations that drive carcinogenesis and neglected the role of negative selection. Some studies have actually concluded that negative selection has no role in cancer evolution. In the present work we have re-examined the role of negative selection in tumor evolution through the analysis of the patterns of somatic mutations affecting the coding sequences of human genes. Our analyses have confirmed that tumor suppressor genes are positively selected for inactivating mutations. Oncogenes, however, were found to display signals of both negative selection for inactivating mutations and positive selection for activating mutations. Significantly, we have identified numerous human genes that show signs of strong negative selection during tumor evolution, suggesting that their functional integrity is essential for the growth and survival of tumor cells.

## Background

### Genetic, epigenetic, transcriptomic and proteomic changes driving carcinogenesis

In the last two decades the rapid advance in genomics, epigenomics, transcriptomics and proteomics permitted an insight into the molecular basis of carcinogenesis. These studies have confirmed that tumors evolve from normal tissues by acquiring a series of genetic, epigenetic, transcriptomic and proteomic changes with concomitant alterations in the control of the proliferation, survival and spread of affected cells.

The genes that play key roles in carcinogenesis, referred to as cancer genes or cancer driver genes are usually assigned to two major categories: proto-oncogenes that have the potential to promote carcinogenesis when activated or overexpressed and tumor suppressor genes that promote carcinogenesis when inactivated or repressed.

There are several alternative mechanisms that can modify the structure or expression of a cancer gene in a way that promotes carcinogenesis. These include subtle genetic changes (single nucleotide substitutions, short indels), major genetic events (deletion, amplification, translocation and fusion of genes to other genetic elements), as well as epigenetic changes affecting the expression of cancer genes. It should be pointed out that these mechanisms are not mutually exclusive: there are many examples illustrating the point that the wild type form of a cancer gene may be converted to a driver gene by multiple types of the above mechanisms.

Exomic studies of common solid tumors revealed that usually several cancer genes harbor subtle somatic mutations (point mutations, short deletions and insertions) in their translated regions but malignancy-driving subtle mutations can also occur in all genetic elements outside the coding region, namely in enhancer, silencer, insulator and promoter regions as well as in 5’- and 3’-untranslated regions. Intron or splice site mutations that alter the splicing pattern of cancer genes can also drive carcinogenesis [1]. A recent study has presented a comprehensive analysis of driver point mutations in non-coding regions across 2,658 cancer genomes [2]. A noteworthy example of how subtle mutations in regulatory regions may activate proto-oncogenes is the telomerase reverse transcriptase gene *TERT* that encodes the catalytic subunit of telomerase. Recurrent somatic mutations in melanoma and other cancers in the *TERT* promoter cause tumor-specific increase of *TERT* expression, resulting in the immortalization of the tumor cell [3].

In addition to subtle mutations, tumors also accumulate major chromosomal changes [4]. Most solid tumors display widespread changes in chromosome number, as well as chromosomal deletions and translocations [5]. Homozygous deletions of a few genes frequently drive carcinogenesis and the target gene involved in such deletions is always a tumor suppressor gene [6]. Somatic copy-number alterations, amplifications of cancer genes are also widespread in various types of cancers. In tumor tissues amplifications usually contain an oncogene whose protein product is abnormally active simply because the tumor cell contains 10 to 100 copies of the gene per cell, compared with the two copies present in normal cells [7, 8]. Chromosomal translocations may also convert wild type forms of tumor suppressor genes into forms that drive carcinogenesis if the translocation inactivates the genes by truncation or by separating them from their promoter. Similarly, translocations may activate proto-oncogenes by changing their regulatory properties [9].

The activity of cancer genes may also be altered by epigenetic mechanisms such as DNA methylation and histone modifications. It is now widely accepted that genetic and epigenetic changes go hand in hand in carcinogenesis: numerous genes involved in shaping the epigenome are mutated in common human cancers, and many genes carrying driver mutations are also affected by epigentic changes [10-14]. For example, promoter hypermethylation events may promote carcinogenesis if they lead to silencing of tumor suppressor genes; the tumor-driving role of promoter methylation is quite obvious in cases when the same tumor suppressor genes are also frequently inactivated by mutations in cancer [15]. Conversely, there is now ample evidence that promoter hypomethylation can promote carcinogenesis if they lead to increased expression of proto-oncogenes [16].

Only recently was it discovered that non-coding RNAs (ncRNAs) also play key roles in carcinogenesis [17]. An explosion of studies has shown that – based on complementary base pairing – ncRNAs may function as oncogenes (by inhibiting the activity of tumor suppressor genes), or as tumor suppressors (by inhibiting the activity of oncogenes or tumor essential genes).

Alterations in the splicing of primary transcripts of protein-coding genes have also been shown to contribute to carcinogenesis. Recent studies on cancer genomes have revealed that recurrent somatic mutations of genes encoding RNA splicing factors (e.g. *SF3B1, U2AF1, SRSF2, ZRSR2*) lead to altered splice site preferences, resulting in cancer-specific mis-splicing of genes. In the case of proto-oncogenes, changes in the splicing pattern may generate active oncoproteins, whereas abnormal splicing of tumor suppressor genes is likely to generate inactive forms of the tumor suppressor protein [18].

There is now convincing evidence that dysregulation of processes responsible for proteostasis also contributes to the development and progression of numerous cancer types [19-21]. Recent studies on tumor tissues have revealed that genetic alterations and abnormal expression of various components of the protein homeostasis pathways (e.g. *FBXW7, VHL*) contribute to progression of human cancers by excessive degradation of tumor-suppressor molecules or through impaired disposal of oncogenic proteins [22-23].

### Hallmarks of cancer and the function of genes involved in carcinogenesis

Hanahan and Weinberg have defined a set of hallmarks of cancer that allow the categorization of cancer genes with respect to their role in carcinogenesis [24]. These hallmarks describe the biological capabilities that are usually acquired during the evolution of tumor cells: these include sustained proliferative signaling, evasion of growth suppressors, evasion of cell death, acquisition of replicative immortality, acquisition of capability to induce angiogenesis and activation of invasion and metastasis. Underlying all these hallmarks are defects in genome maintenance that help the acquisition of the above capabilities. Additional emerging hallmarks of potential generality have been suggested to include tumor promoting inflammation, evasion of immune destruction and reprogramming of energy metabolism in order to most effectively support neoplastic proliferation [24].

**Figure 1** summarizes our current view of the cellular processes that play key roles in tumor evolution to emphasize their contribution to the various major hallmarks of cancer. In this representation changes in the maintenance of the genome, epigenome, transcriptome and proteome occupy a central position since they increase the chance that various constituents of other cellular pathways will experience alterations that favor the acquisition of capabilities that permit the proliferation, survival and metastasis of tumor cells.

**Figure 1.**
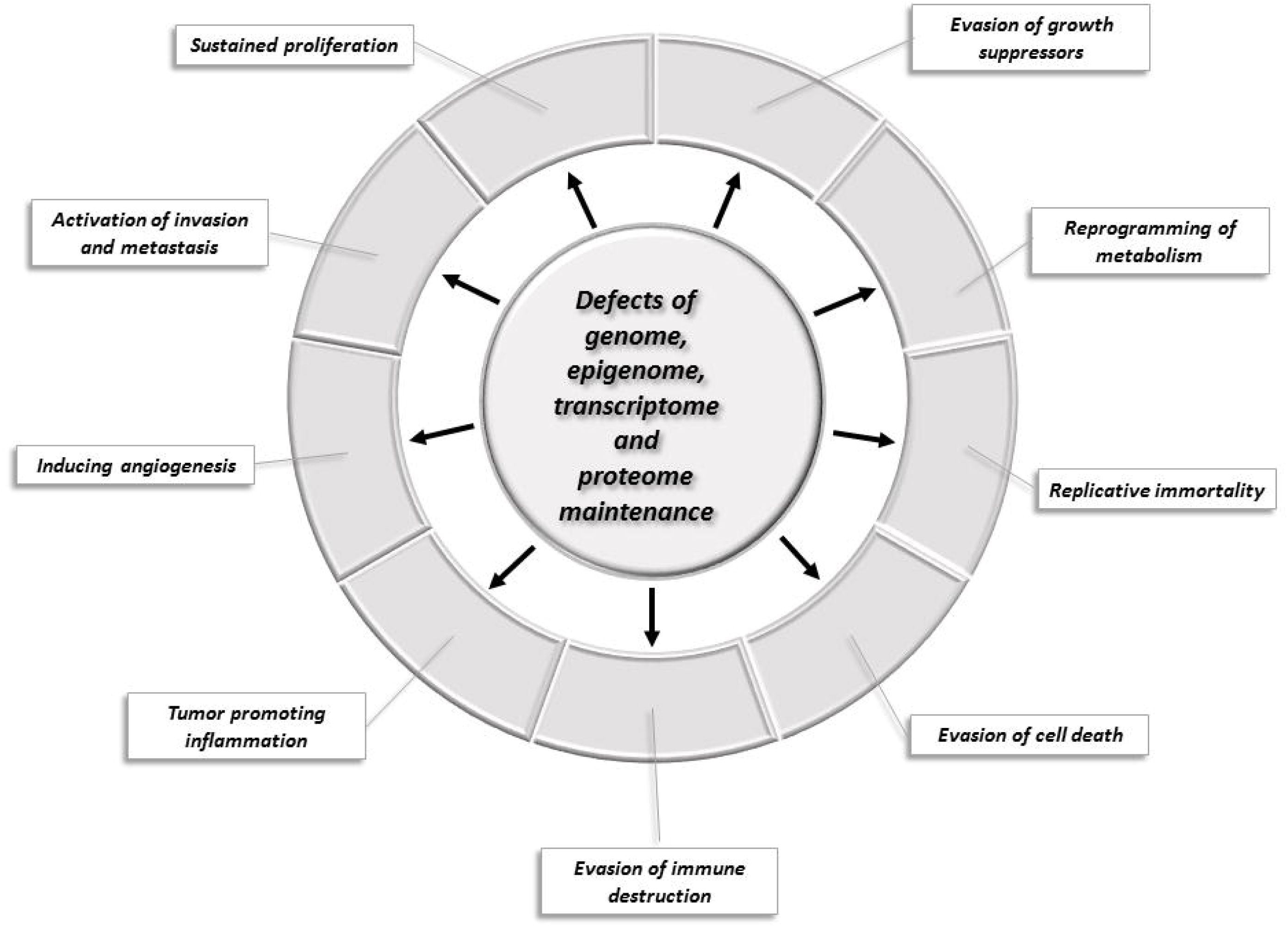
Changes of key cellular processes contributing to carcinogenesis. The central circle refers to processes involved in the maintenance of the integrity of the genome, epigenome, transcriptome and proteome: defects in these processes increase the chance that genes and proteins of other cellular pathways (represented by segments of the outer circle) will suffer alterations that favor the acquisition of capabilities that permit the proliferation, survival and metastasis of tumor cells.

### Chronology of tumor evolution: initiation and progression

In the first phase of carcinogenesis a cell may acquire a mutation that permits it to proliferate abnormally, in the next phase other mutations allow the expansion of cell number and this process of mutations (and associated epigenetic, transcriptomic and proteomic alterations) continues, thus generating a primary tumor that can eventually metastasize to distant organs. Recent studies on the chronology and genomic landscape of the events that drive carcinogenesis in multiple myeloma suggest that complex structural changes of the genome occur early, whereas point mutations occur in later disease phases [25].

Individual instances of cancer may be initiated by specific combinations of mutations affecting a small number of cancer genes. According to current estimates the number of cancer driving mutations needed for the full development of cancer ranges from two-eight depending on cancer type [26-27]. A recent integrative analysis of 2,658 whole-cancer genomes and their matching normal tissues across 38 tumor types revealed that, on average, cancer genomes contain 4–5 driver mutations [28].

Although the temporal order of the mutations affecting genes of key pathways differs among cancer types, it appears that a common feature is that mutations of genes that regulate apoptosis occur in the early phases of tumor progression, whereas mutations of genes involved in invasion pathways are observed only in the last stages of carcinogenesis [29]. It has been suggested that the reason why the loss of apoptotic control is a critical step for initiating cancer is that the larger the surviving cell population, the higher the number of cells at risk of acquiring additional mutations.

Analyses of the mutation landscapes and evolutionary trajectories of various tumor tissues have identified *BRAF, KRAS, TP53, RB* or *APC* as the key genes whose mutation is most likely to initiate carcinogenesis, permitting the cell to divide abnormally [26]. In the case of ovarian cancers *TP53* mutation is believed to be the earliest tumorigenic driver event, with presence in nearly all cases of ovarian cancer [30]. The prevalence of *TP53* mutations and *BRCA* deficiency in these tumors leads to incompetent DNA repair promoting subsequent steps of carcinogenesis. Studies on the evolution of melanoma from precursor lesions have revealed that the vast majority of melanomas harbored *TERT* promoter mutations, indicating that these immortalizing mutations are selected at an unexpectedly early stage of neoplastic progression [31].

The life history and evolution of mutational processes and driver mutation sequences of 38 types of cancer has been analyzed recently by whole-genome sequencing analysis of 2,658 cancers. This study has shown that early oncogenesis is characterized by mutations in a constrained set of driver genes and that the driver mutations that most commonly occur in a given cancer also tend to occur the earliest [32].

### Cancer genes and passenger genes

The prominent role of *KRAS* and *TP53* genes in initiating carcinogenesis is also reflected by the observation that their mutation rate in tumors far exceeds those of other genes, suggesting that their mutations are subject to positive selection during tumor evolution. Since one of the major goals of cancer research is to identify all genes that drive carconogenesis, several types of approaches have been developed based on the premise that, thanks to positive selection, the rate of mutation of ‘driver genes’ must be significantly higher in the tumor tissue than those of ‘passenger genes’ that have no role in the development of cancer but simply happen to mutate in the same tumor [33-34].

Unfortunately, methods based on mutation frequency alone cannot reliably indicate which genes are cancer drivers because the background mutation rates differ significantly as a consequence of intrinsic characteristics of DNA sequence and chromatin structure [35]. Mutation hotspots that depend on the nucleotide sequence context, the mechanism of mutagenesis and the action of the repair and replication machineries are called intrinsic mutation hotspots [36]. Genes enriched in intrinsic mutation hotspots may accumulate mutations at a significantly higher rate than other genes, creating the illusion of positive selection: based on recurrent mutations they may be mistakenly identified as cancer driver genes [37-38].

In principle this danger may be avoided if we compare the mutation pattern of the gene in the tumor tissue with that in the normal tissue the tumor has originated from. However, since the rate of mutation in such hotspots depends not only on the nucleotide sequence but also on the mechanism of mutagenesis and the integrity of DNA repair pathways [38-39] mutation hotspots that arise during carcinogenesis could still create the illusion of positive selection.

Chromatin organization is also known to have a major influence on regional mutation rates in human cancer cells [40-41]. Since large-scale chromatin features, such as replication time and accessibility influence the rate of mutations, this may hinder the distinction of cancer driver genes whose high mutation rate reflects positive selection and passenger genes whose high mutation rate is the result of the distinctive features of the chromatin region in which they reside. Moreover, since the cell-of-origin chromatin organization shapes the mutational landscape, rates of somatic mutagenesis of genes in cancer are highly cell-type-specific [42]. Actually, since regional mutation density of ‘passenger’ mutations across the human chromosomes is correlated with the cell type the tumor had originated from, this feature may be used to classify human tumors [43].

By comparing the exome sequences of 3,083 tumor-normal pairs Lawrence and coworkers [44] have discovered an extraordinary variation in mutation frequency and spectrum within cancer types across the genome, which is strongly correlated with DNA replication timing and transcriptional activity. The authors have shown that by incorporating mutational heterogeneity into their analyses, many of the apparent artefactual findings could be eliminated improving the identification of genes truly associated with cancer. In a more recent study Lawrence *et al*. [45] compared the frequency of somatic point mutations in exome sequences from 4,742 human cancers and their matched normal-tissue samples across 21 cancer types and identified 33 genes that were not previously known to be significantly mutated in cancer. They have concluded that a total of 224 genes are significantly mutated in one or more tumor types.

However, since background mutational frequency estimates are not sensitive enough, the list of driver genes identified on the basis of somatic mutation rate alone is likely to be incomplete, but may also contain false positives. To overcome these limitations of mutation rate-based approaches, several attempts have been made to use additional features that may distinguish driver genes and passenger genes. A major group of such approaches incorporates observations about the impact of mutations on the structure and function of well-characterized proteins encoded by proto-oncogenes and tumor suppressor genes. Several computational methods have been developed to identify driver missense mutations most likely to generate functional changes that causally contribute to tumorigenesis [46-48].

In a different type of approach Youn and Simon [49] identified cancer driver genes as those for which the non-silent mutation rate is significantly greater than a background mutation rate estimated from silent mutations, indicating that the non-silent mutations are subject to positive selection. The authors have identified 28 genes as driver genes, the majority of the significant matches (e.g. *EGFR, CDKN2A, KRAS, STK11, TP53, NF1, RB1 PTEN* and *NRAS*), were well characterized oncogenes or tumor suppressor genes known from earlier studies.

In a more recent study Zhou *et al*. [50] have identified 365 genes for which the ratio of the nonsynonymous to synonymous substitution rate was significantly increased, suggesting that they are subject to the positive selection of driver mutations. It should be pointed out here that an obvious limitation of such approaches is that they implicitly assume that synonymous substitutions are – *per definitionem* – silent and are thus selectively neutral since they do not affect the sequence of the protein. However, the fact should not be ignored that this is not necessarily true: some synonymous mutations may have a significant impact on splicing, RNA stability, RNA folding and translation of the transcript of the affected gene and may thus actually act as driver mutations [51-53].

Vogelstein *et al*. [54] have used a heuristic approach to identify cancer driver genes. Since the patterns of mutations in the first and best-characterized oncogenes and tumor suppressor genes were found to be highly characteristic and nonrandom, the authors assumed that the same characteristics are generally valid and may be used to identify previously uncharacterized cancer genes. For example, since many known oncogenes were found to be recurrently mutated at the same amino acid positions, to classify a gene as an oncogene, it was required that >20% of the recorded mutations in the gene are at recurrent positions and are missense. Similarly, since in the case of known tumor suppressors the driver mutations most frequently truncate the tumor suppressor proteins, to be classified as a tumor suppressor gene, it was required that >20% of the recorded mutations in the gene are truncating (nonsense or frameshift) mutations. Along these lines, Vogelstein *et al*., [54] have analyzed the patterns of the subtle mutations in the Catalogue of Somatic Mutations in Cancer (COSMIC) database to identify driver genes. As a proof of the reliability of this “20/20 rule” it was emphasized that all well-documented cancer genes passed these criteria [54]. Although this indicates that the approach detects known cancer genes, it does not guarantee that it detects all driver genes. Acknowledging that additional cancer driver genes might exist, the authors have introduced the term “Mut-driver gene” for genes that contain a sufficient number or type of driver gene mutations to unambiguously distinguish them from other genes, whereas for cancer genes that are expressed aberrantly in tumors but not frequently mutated they proposed the term “Epi-driver gene”.

Based on these analyses, it has been concluded that out of the 20,000 human protein-coding genes, only 125 genes qualify as Mut-driver genes, of these, 71 are tumor suppressor genes and 54 are oncogenes [54]. Although the authors have expressed their conviction that nearly all genes mutated at significant frequencies had already been identified and that the number of Mut-driver genes is nearing saturation, this conclusion may not be justified since the criteria used to identify oncogenes and tumor suppressors appear to be too stringent and somewhat arbitrary.

In search of additional cancer driver genes Tamborero *et al*. [55] employed five complementary methods to find genes showing signals of positive selection and identified a list of 291 “high-confidence cancer driver genes” acting on 3,205 tumors from 12 different cancer types. Bailey *et al*. [56] used multiple advanced algorithms to identify cancer driver genes and driver mutations. Based on their PanCancer and PanSoftware analysis spanning 9,423 tumor exomes, comprising all 33 of The Cancer Genome Atlas projects and using 26 computational tools they have identified 299 driver genes showing signs of positive selection. Their sequence and structure-based analyses detected >3,400 putative missense driver mutations and 60%–85% of the predicted mutations were validated experimentally as likely drivers.

Zhao *et al*., [57] have developed driverMAPS (Model-based Analysis of Positive Selection), a model-based approach for driver gene identification that captures elevated mutation rates in functionally important sites and spatial clustering of mutations. Using this approach the authors have identified 255 known driver genes as well as 170 putatively novel driver genes.

Currently COSMIC, the Catalogue Of Somatic Mutations In Cancer (https://cancer.sanger.ac.uk/cosmic) is the most detailed and comprehensive resource for exploring the effect of subtle somatic mutations of driver genes in human cancer [58] but COSMIC also covers all the genetic mechanisms by which somatic mutations promote cancer, including non-coding mutations, gene fusions, copy-number variants. In parallel with COSMIC’s variant coverage, the Cancer Gene Census (CGC, https://cancer.sanger.ac.uk/census) describes a curated catalogue of genes driving every form of human cancer [59]. CGC has recently introduced functional descriptions of how each gene drives disease, summarized into the cancer hallmarks. The 2018 CGC describes in detail the effect of a total of 719 cancer-driving genes, encompassing Tier 1 genes (574 genes) and a list of Tier 2 genes (145 genes) from more recent cancer studies that show less detailed indications of a role in cancer.

In a different type of approach, Torrente *et al*. [60] used comprehensive maps of human gene expression in normal and tumor tissues to identify cancer related genes. These analyses identified a list of genes with systematic expression change in cancer. The authors have noted that the list is significantly enriched with known cancer genes from large, public, peer-reviewed databases, whereas the remaining ones were proposed as new cancer gene candidates. A recent study has provided a comprehensive catalogue of cancer-associated transcriptomic alterations with the top-ranking genes carrying both RNA and DNA alterations. The authors have noted that this catalogue is enriched for cancer census genes [61].

Using transposon mutagenesis in mice several laboratories have conducted forward genetic screens and identified thousands of candidate genetic drivers of cancer that are highly relevant to human cancer. The Candidate Cancer Gene Database (CCGD, http://ccgd-starrlab.oit.umn.edu/) is a manually curated database containing a unified description of all identified candidate driver genes [62].

In summary, although a variety of approaches have been developed to identify ‘cancer genes’, there is significant disagreement as to the number of genes involved in carcinogenesis. Some of the studies argue that the number is in the 200-700 range, other approaches suggest that their number may be much higher. Since the ultimate goal of cancer genome projects is to discover therapeutic targets it is important to identify all true cancer genes and distinguish them from passenger genes and candidates that do not play a significant role in the process of carcinogenesis.

It should be pointed out, however, that the majority of genomics-based methods were biased as they defined the aim of cancer genomics as the identification of mutated driver genes (equating them with ‘cancer genes’) that are causally implicated in oncogenesis [63]. In all these studies, the underlying rationale for interpreting a mutated gene as causal in cancer development is that the mutations are likely to have been positively selected because they confer a growth advantage on the cell population from which the cancer has developed. An inevitable consequence of this focus on positive selection was that most studies neglected the possibility that negative selection may also play a significant role in tumor evolution.

### Carcinogenesis as an evolutionary process

In principle, with respect to its effect on carcinogenesis, a somatic mutation may promote or may hinder carcinogenesis or may have no effect on carcinogenesis. In cancer genomics the mutations that promote carcinogenesis (and are subject to positive selection during tumor evolution) are called ‘driver mutations’ to distinguish them from ‘passenger mutations’ that do not play a role in carcinogenesis (and are not subject to positive or negative selection during tumor evolution). Mutations that impair the growth, survival and invasion of tumor cells have received much less attention although they are also expected to play a significant role in shaping the mutation pattern of genes during carcinogenesis. Hereafter we will refer to this category of mutations as ‘cancer blocking mutations’ since they are deleterious from the perspective of tumor growth.

In cancer research genes are usually assigned to just two categories with respect to their role in carcinogenesis: 1) ‘passenger genes’ (or bystander genes) that play no significant role in carcinogenesis and their mutations are passenger mutations; 2) ‘driver genes’ that drive carcinogesis when they acquire driver mutations.

The problem with this usual binary driver gene-passenger gene categorization is that some genes with functions essential for the growth and survival of tumor cells (hereafter referred to as ‘tumor essential genes’) may not easily fit into the usual ‘driver gene’ category. It is to be expected that during tumor evolution the coding sequences of driver genes (tumor suppressor genes, proto-oncogenes), passenger genes and tumor essential genes will experience markedly different patterns of selection. The mutation patterns of selectively neutral, *bona fide* passenger genes are likely to reflect the lack of positive and negative selection, whereas in the case of tumor essential genes purifying selection is expected to dominate. In the case of tumor suppressor genes, the mutation pattern would reflect positive selection for truncating driver mutations.

Proto-oncogenes, however are expected to show signs of both positive selection for activating mutations and negative selection for inactivating, ‘cancer blocking’ mutations as their activity is essential for their oncogenic role. It must be emphasized that in the coding regions of proo-oncogenes positive selection for driver mutations is expected to favor nonsynonymous substitutions over synonymous substitutions only at sites that are critical for the novel, oncogenic function. For these sites (and these sites only) the ratio of nonsynonymous to synonymous rates is expected to be significantly greater than one reflecting positive selection. If there are many such sites in a protein, or selection is extremely strong the overall nonsynonymous to synonymous ratio for the entire protein may also be significantly higher than one, otherwise the effect of positive selection on the synonymous to nonsynonymous ratio may be overridden by purifying selection at other sites [64].

In harmony with some of these expectations, using just the ratio of the nonsynonymous to synonymous substitution rate as a measure of positive or negative selection, Zhou *et al*. [50] have shown that in cancer genomes, the majority of genes had nonsynonymous to synonymous substitution rate values close to one, suggesting that they belong to the passenger gene category. The authors have identified a total of 365 potential cancer driver genes that had nonsynonymous to synonymous substitution rate values significantly greater than one (reflecting the dominance of positive selection), whereas 923 genes had nonsynonymous to synonymous substitution rate values significantly less than one, leading the authors to suggest that these negatively selected genes may be important for the growth and survival of cancer cells.

Realizing that genes whose wild-type coding sequences are needed for tumor growth are also of key interest for cancer research Weghorn and Sunyaev [65] have also focused on the role of negative selection in human cancers. As the authors have pointed out, identification and analysis of true negatively selected, ‘undermutated’ genes is particularly difficult since the sparsity of mutation data results in lower statistical power, making conclusions less reliable. Although the signal of negative selection was exceedingly weak, the authors have noted that the group of negatively selected candidate genes is enriched in cell-essential genes identified in a CRISPR screen [66], consistent with the notion that one of the potential causes of negative selection is the maintenance of genes that are responsible for basal cellular functions. Based on pergene estimates of negative selection inferred from the pan-cancer analysis the authors have identified 147 genes with strong negative selection. The authors have noted that among the 13 genes showing the strongest signs of negative selection there are several genes (*ATAT1, BCL2, CLIP1, GALNT6, CKAP5* and *REV1*) that are known to promote carcinogenesis.

In a similar work Martincorena *et al.* [67] have used the normalized ratio of non-synonymous to synonymous mutations, to quantify selection in coding sequences of cancer genomes. Using a nonsynonymous to synonymous substitution rate value >1 as a marker of cancer genes under positive selection, they have identified 179 cancer genes, with about 50% of the coding driver mutations being found to occur in novel cancer genes. The authors, however, have concluded that purifying selection is practically absent in tumors since nearly all (> 99%) coding mutations are tolerated and escape negative selection. The authors have suggested that this remarkable absence of negative selection on coding point mutations in cancer indicates that the vast majority of genes are dispensable for any given somatic lineage, presumably reflecting the buffering effect of diploidy and the inherent resilience and redundancy built into most cellular pathways.

The key message of Martincorena *et al.* [67] that negative selection has no role in cancer evolution had a major impact on cancer genomics research as reflected by several commentaries in major journals of the field that have propagated this conclusion [68-70].

In view of the contradicting conclusions of Martincorena *et al.* [67], Weghorn and Sunyaev [65] and Zhou *et al*., [50] it is important to reexamine the significance of negative selection of protein-coding genes in tumor evolution. As pointed out above, detection of negative selection may have been impeded by the fact that putative tumor essential genes – unlike classical driver genes – are likely to be undermutated and the tools used for the analyses may have not been sensitive enough to identify weaker signals of selection. In the present work we have tried to overcome these problems by limiting our work to transcripts of human genes that have at least 100 verified somatic mutations. Furthermore, to increase the sensitivity of our approach we have used analyses combining different signals of selection manifested in synomymous, nonsynonymous, nonsense substitutions as well as subtle inframe and frameshift indels.

In the present work we have identified a large group of human genes that show clear signs of negative selection during tumor evolution, suggesting that their functional integrity is essential for the growth and survival of tumor cells. Significantly, the group of negatively selected genes includes genes that play critical roles in the Warburg effect of cancer cells, others mediate invasion and metastasis of tumor cells, indicating that negatively selected tumor essential genes may prove a rich source for novel targets for tumor therapy.

Improved detection of signals of selection has also permitted the identification of numerous novel cancer gene candidates that are likely to play important roles in carcinogenesis as tumor suppressor genes or as oncogenes.

## Results

Cancer somatic mutation data were extracted from COSMIC v88, the Catalogue Of Somatic Mutations In Cancer, which includes single nucleotide substitutions and small insertions/deletions affecting the coding sequence of human genes. The downloaded file (CosmicMutantExport.tsv, release v88) contained data for 29415 transcripts (**Supplementary file 1**). For all subsequent analyses we have retained only transcripts containing mutations that were annotated under ‘Mutation description’ as substitution or subtle insertion/deletion. This dataset contained data for 29405 transcripts containing 6449721 mutations (substitution and short indels, SSI) and 29399 transcripts containing 6141650 substitutions only (SO). **Supplementary file 2** contains the metadata for these SO and SSI datasets.

To increase the statistical power of our analyses we have limited our work to transcripts that have at least 100 somatic mutations. Hereafter, unless otherwise indicated, our analyses refer to datasets containing transcripts with at least 100 somatic mutations. This limitation eliminated ∼38% of the transcripts that contain very few mutations but reduced the number of total mutations only by 9% (**Supplementary file 1**). It should be noted that this limitation increases the statistical power of our analyses but disfavors the identification of some negatively selected genes.

Since we were interested in the selection forces that operate during tumor, only confirmed somatic mutations were included in our analyses. In COSMIC such mutations are annotated under ‘Mutation somatic status’ as Confirmed Somatic, i.e. confirmed to be somatic in the experiment by sequencing both the tumor and a matched normal tissue from the same patient. As to ‘Sample Type, Tumor origin’: we have excluded mutation data from cell-lines, organoid-cultures, xenografts since they do not properly represent human tumor evolution at the organism level. We have found that by excluding cell lines we have eliminated many artifacts of spurious recurrent mutations caused by repeated deposition of samples taken from the same cell-line at different time-points. To eliminate the influence of polymorphisms on the conclusions we retained only somatic mutations flagged ‘n’ for SNPs. (**Supplementary file 1**). **Supplementary file 3** contains the metadata for transcripts containing at least 100 confirmed somatic, non polymorphic mutations identified in tumor tissues.

As the gold standard of ‘known’ cancer genes we have used the lists of oncogenes (OG) and tumor suppressor genes (TSG) identified by Vogelstein *et al*. [54]. As another list of ‘known’ cancer genes we have also used the genes of the Cancer Gene Census [59].

In our datasets the numerical variables for sets of human genes were expressed as mean and standard deviation for each group of data. For each variable, the means for the various groups were compared using the t-test for independent samples. Statistical significance was set as a P value of <0.05.

We have used several approaches to estimate the contribution of silent, amino acid changing and truncating mutations to somatic mutations of human protein-coding genes during tumor evolution. We have used two major types of calculations: one in which we have restricted our analyses to single nucleotide substitutions (hereafter referred to as SO for ‘substitution only’) and a version in which we have also taken into account subtle indels (hereafter referred to as SSI for ‘substitutions and subtle indels’).

### Analyses of subtle mutations in tumor tissues

In the simplest case we have calculated for each transcript the fraction of somatic substitutions that could be assigned to the synonymous (fS), nonsynonymous (fM) and nonsense mutation (fN) category (**Supplementary files 2** and **3**). In the version that also included data for indels we have calculated the fraction of mutations corresponding to synonymous substitutions (indel_fS), but have merged nonsynonymous substitutions and short inframe indels in the category of mutations that lead to changes in the amino acid sequence (indel_fM). Nonsense substitutions and short frame-shift indels were included in the third category of mutations (indel_fN) as both types of mutation lead eventually to stop codons that truncate the protein (**Supplementary files 2** and **3**).

Analyses of datasets (**Supplementary file 3**) containing substitutions only have shown that in 3D scatter plots transcripts are present in a cluster (**Figure 2A**) characterized by values of 0.2436±0.0619, 0.7090±0.0556 and 0.0475±0.0322 for synonymous, nonsynonymous and nonsensense substitutions, respectively. The mean values for synonymous, nonsynonymous and nonsensense substitutions in this cluster are very close to those expected if we assume that the structure of the genetic code has the most important role in determining the probability of somatic substitutions during tumor evolution of human genes (**Supplementary file 4**). Since each codon can undergo nine types of single-base substitutions, point mutations in the 61 sense codons can lead to 549 types of single-base substitutions. Of these, 392 result in the replacement of one amino acid by another (nonsynonymous substitutions), whereas 134 result in silent mutations (synonymous substitutions) and 23 generate stop codons [64]. Based on the structure of the genetic code, in the absence of selection one would thus expect that a fraction of 0.24408 would be synonymous, 0.71403 of the single-base substitutions would be nonsynonymous and 0.04189 would be nonsense mutations.

**Figure 2.**
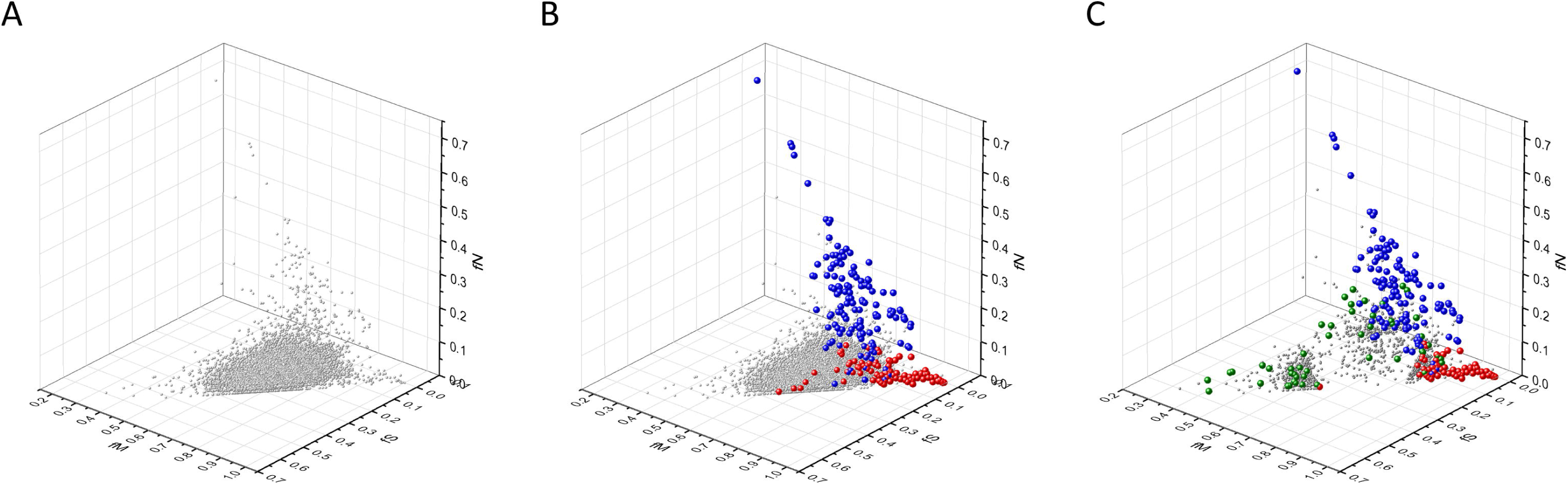
Analyses of fS, fM and fN parameters of human protein-coding genes of tumor tissues. The figure shows the results of the analysis of 13803 transcripts containing at least 100 subtle, confirmed somatic mutations from tumor tissues, including only mutations identified as not SNPs. Axes *x, y* and *z* represent the fractions of somatic single nucleotide substitutions that are assigned to the synonymous (fS), nonsynonymous (fM) and nonsense (fN) categories, respectively. In **Panel A** each gray ball represents a human transcript; note that the majority of human genes are present in a dense cluster. **Panel B** highlights the positions of transcripts of the genes identified by Vogelstein *et al*., (2013) as oncogenes (OGs, large red balls) or tumor suppressor genes (TSGs, large blue balls). It is noteworthy that these driver genes separate significantly from the central cluster and from each other: OGs have a significantly larger fraction of nonsynonymous, whereas TSGs have significantly larger fraction of nonsense substitutions. **Panel C** shows data only for candidate cancer genes present in the CG_SO^2SD^_SSI^2SD^ list (see **Supplementary file 6**). The positions of transcripts of the genes identified by Vogelstein *et al*., (2013) as oncogenes (OGs, large red balls) or tumor suppressor genes (TSGs, large blue balls) are highlighted. The positions of novel cancer gene transcripts validated in the present work are highlighted as large green balls.

It is noteworthy, however, that the fS, fM and fN values of the best known cancer genes Vogelstein *et al*., [54] deviate from those characteristic of the majority of human genes (**Figure 2B**). In harmony with earlier observations, the values for OGs show a marked shift of fM to higher values, reflecting positive selection for missense mutations, whereas the fN values of TSGs are significantly higher, reflecting positive selection for truncating nonsense mutations (**Supplementary file 4).**

The genes (6198 transcripts) with values that deviate from mean values of fS, fM and fN by more than 1SD have also included the majority of OGs and TSGs; only 4 OG transcripts are present in the central cluster deviating from mean fM, fS and fN values by ≤1SD. It is noteworthy that the 6198 transcripts also contained the majority (440 out of 741) of the transcripts of CGC genes, suggesting that the mutation pattern of most CGC genes also deviates significantly from those of passenger genes present in the central cluster (**Supplementary file 4**). The genes in the central cluster are characterized by fraction values of 0.24548±0.03079, 0.71084±0.0274 and 0.04368±0.01572 for synonymous, nonsynonymous and nonsensense substitutions, respectively. Note that these values are very close to those expected from the structure of the genetic code in the absence of selection (**Supplementary file 4**). This central cluster of genes (**Supplementary file 3**) is hereafter referred to as PG_SO^f_1SD^ (for Passenger Gene_Substitution Only deviating from mean fM, fS and fN values by ≤1SD).

The genes (1060 transcripts) with values that deviate from mean values of fS, fM and fN by more than 2SD included 62 OG and 119 TSG driver gene transcripts, but 42 driver gene transcripts were present in the cluster that deviates from the mean by ≤2SD. Using this more stringent cut-off value the number of additional CGC genes represented in the 1060 transcripts was reduced to 142 out of 741 (**Supplementary file 4**). This candidate cancer gene set defined by 2SD cut-off value is hereafter referred to as CG_SO^f_2SD^ for Cancer Gene_Substitution Only deviating from mean fM, fS and fN values by more than 2SD (**Supplementary file 4**).

Out of the 1060 transcripts present in CG_SO^f_2SD^, 737 transcripts are derived from genes that are not included in the OG, TSG and CGC cancer gene lists (**Supplementary files 3** and **4**). Since the majority of these 737 transcripts (derived from 617 genes) have parameters that assign them to the OG or TSG clusters, we assume that they also qualify as candidate oncogenes or tumor suppressor genes. There is, however, a third group of genes that deviate from both the central passenger gene cluster and the clusters of OGs and TSGs (**Figure 2B**): their high fS and low fM and fN values suggest that they experience purifying selection during tumor evolution, raising the possibility that they may correspond to tumor essential genes (TEGs) important for the growth and survival of tumors. The 617 putative cancer genes listed in CG_SO^f_2SD^ of **Supplementary file 3**, were subjected to further analyses to decide whether they qualify as candidate oncogenes, tumor suppressor genes, tumor essential genes or the deviation of their mutation pattern from those of passenger genes is not the result of selection (see section on **Analyses of candidate cancer gene sets**).

Known cancer genes (OGs and TSGs) also separate from the majority of human genes in 3D scatter plots of parameters rSM, rNM, rNS defined as the ratio of fS/fM, fN/fM, fN/fS, respectively (**Figure 3**). In these plots OGs separate from the central cluster in having lower rSM and rNM values, whereas TSGs have higher rNS and rNM values than those of the central cluster ((**Figure 3, A1, A2, Supplementary file 4**). The set of genes (4744 transcripts) with values that deviate from the mean by more than 1SD contained 80 OG transcripts, 132 TSG transcripts and an additional 371 CGC gene transcripts. The central cluster of genes (that deviate from mean rSM, rNM and rNS values by ≤1SD is hereafter referred to as PG_SO^r2_1SD^ (for Passenger Gene_Substitution Only deviating from mean rSM, rNM and rNS values by ≤ 1SD).

**Figure 3.**
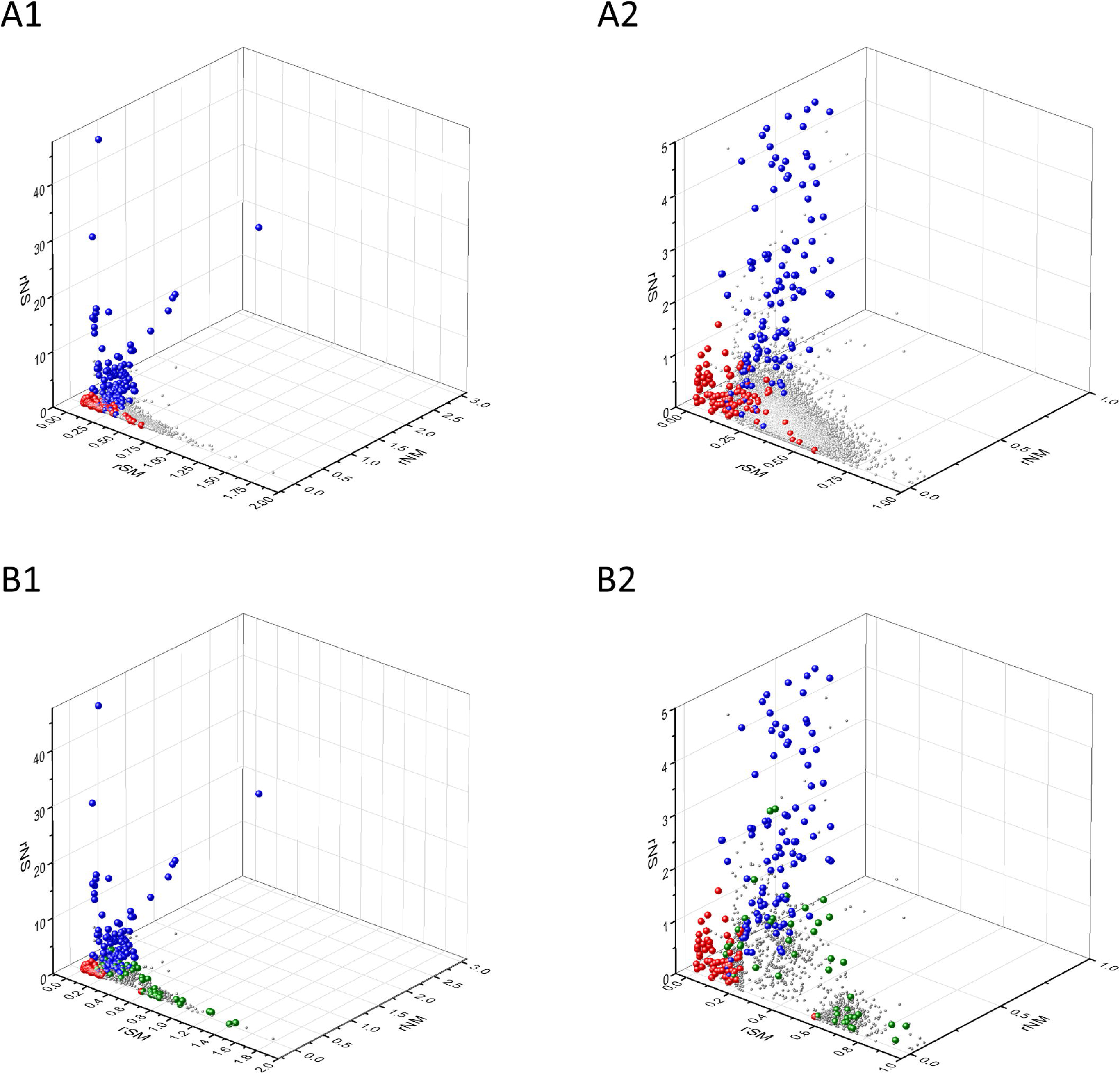
Analyses of rSM, rNM, rNS parameters of human protein-coding genes of tumor tissues. The figure shows the results of the analysis of 13803 transcripts containing at least 100 subtle, confirmed somatic mutations from tumor tissues, including only mutations identified as not SNPs. Axes *x, y* and *z* represent the rSM, rNM, rNS values defined as the ratio of fS/fM, fN/fM, fN/fS, respectively. Each ball represents a human transcript; the positions of transcripts of the genes identified by Vogelstein *et al*., (2013) as oncogenes (OGs, large red balls) or tumor suppressor genes (TSGs, large blue balls) are highlighted. **Panels A1, A2** show the distribution of the 13803 transcripts at different magnification. Note that the majority of human genes are present in a dense cluster but known OGs and TSGs separate significantly from the central cluster and from each other. The rNS and rNM values of TSGs are higher, whereas the rSM and rNM values of OGs are lower than those of passenger genes. **Panels B1, B2** show data only for candidate cancer genes present in the CG_SO^2SD^_SSI^2SD^ list (see **Supplementary file 6**). The positions of transcripts of the genes identified by Vogelstein *et al*., (2013) as oncogenes (OGs, large red balls) or tumor suppressor genes (TSGs, large blue balls) are highlighted. The positions of novel cancer gene transcripts validated in the present work are highlighted as large green balls.

The candidate cancer gene set defined by 2SD cut-off value (**Supplementary file 3**) is hereafter referred to as CG_SO^r2_2SD^ for Cancer Gene_Substitution Only deviating from mean rSM, rNM, rNS values by more than 2SD (**Supplementary file 4**). This gene set has a total of 780 transcripts, containing 40 transcripts of OGs, 103 transcripts of TSGs genes, an additional 79 transcripts of CGC genes and 558 transcripts derived from 468 genes that are not found in the OG, TSG and CGC cancer gene lists (**Supplementary file 4**).

The mean parameters of TSGs differ markedly from those of passenger genes in that rNS and rNM values are higher (**Figure 3A1, A2. Supplementary file 4**), reflecting the dominance of positive selection for inactivating mutations. The parameters for OGs on the other hand, differ from those of passenger genes in that rSM and rNM values are significantly lower (**Figure 3A1, A2** and **Supplementary file 4**), reflecting positive selection for missense mutations and negative selection of nonsense mutations. Interestingly, in these plots some oncogenes (e.g. *BCL2*) have unusually high values of rSM and low values of rNM (e.g. **Figures 3A1, A2** and **Supplementary file 3**) suggesting that in the case of these oncogenes purifying selection may dominate over positive selection for amino acid changing mutations.

As mentioned above, the candidate cancer gene set defined by a cut-off value of 2SD contains 558 transcripts derived from 468 genes that are not found in the OG, TSG or CGC lists. Since the majority of these genes have parameters that assign them to the OG or TSG clusters, they can be regarded as candidate oncogenes or tumor suppressor genes. It is noteworthy, however, that there is a group of genes that deviate from the clusters of passenger genes, OGs and TSGs in that they have unusually high rSM values and low rNM and rNS values. Since these values may be indicative of purifying selection we assumed that they may correspond to tumor essential genes important for the growth and survival of tumors. The 468 putative cancer genes listed in CG_SO^r2_2SD^ of **Supplementary file 3** were subjected to further analyses to decide whether they qualify as candidate oncogenes, tumor suppressor genes or tumor essential genes (see section on **Analyses of candidate cancer gene sets**).

The separation of known cancer genes from the majority of human genes is even more obvious in 3D scatter plots of parameters rSMN, rMSN and rNSM defined as the ratio of fS/(fM+fN), fM/(fS+fN) and fN/(fS+fM), respectively (**Figure 4 A1, A2**). In these plots the gene transcripts are present in a three-pronged cluster, with OGs and TSG being present on separate spikes of this cluster (**Figure 4**).

**Figure 4.**
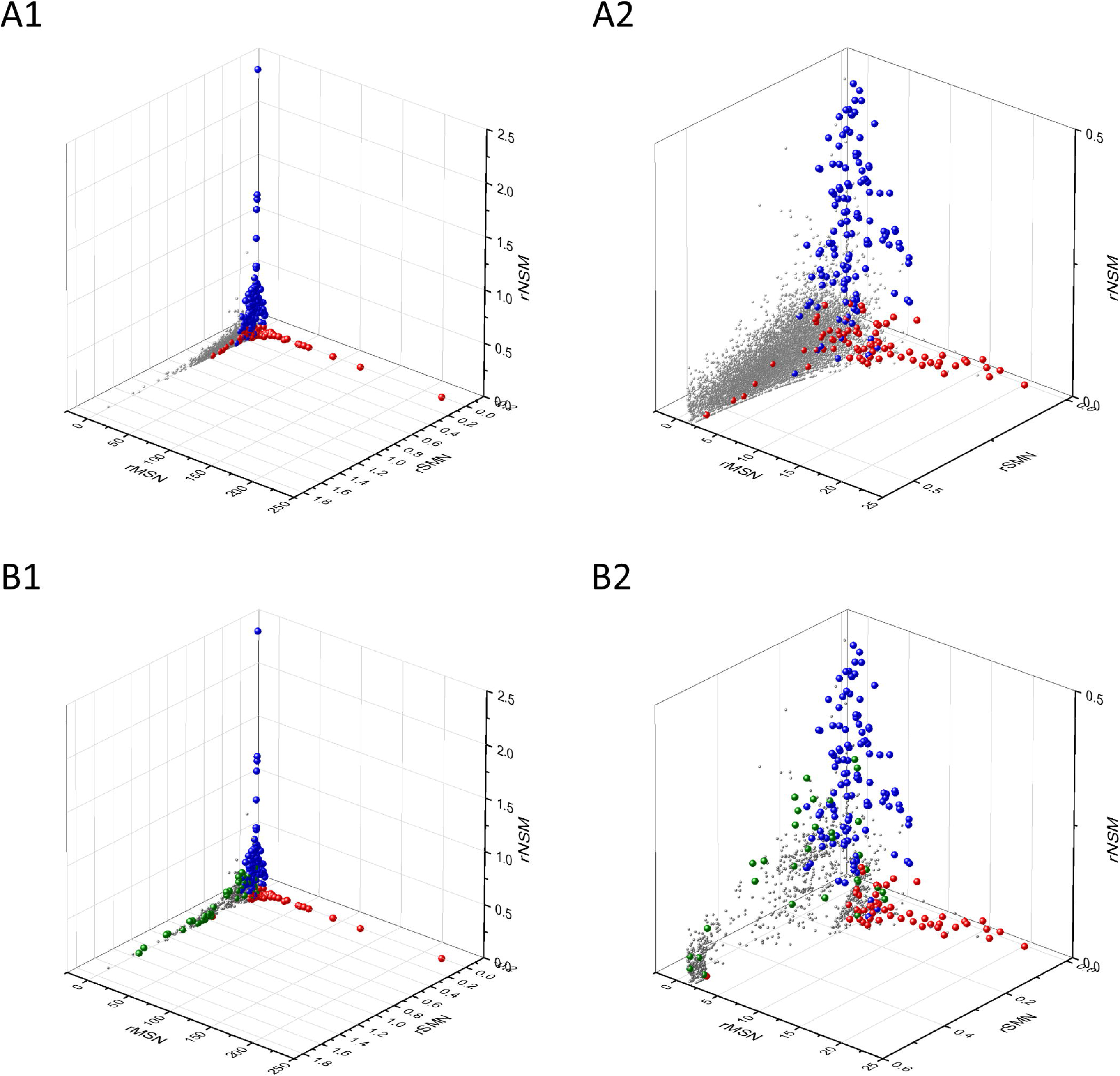
Analyses of rSMN, rMSN and rNSM parameters of human protein-coding genes of tumor tissues. The figure shows the results of the analysis of transcripts containing at least 100 subtle, confirmed somatic mutations from tumor tissues, including only mutations identified as not SNPs. Axes *x, y* and *z* represent the rSMN, rMSN and rNSM defined as the ratio of fS/(fM+fN), fM/(fS+fN) and fN/(fS+fM). Each ball represents a human transcript; the positions of transcripts of the genes identified by Vogelstein *et al*., (2013) as oncogenes (OGs, large red balls) or tumor suppressor genes (TSGs, large blue balls) are highlighted. **Panels A1, A2** show the distribution of the 13803 transcripts at different magnification. Note that the majority of human genes are present in a dense cluster but known OGs and TSGs separate significantly from the central cluster and from each other. The rNSM values of TSGs are higher, their rMSN and rSMN are lower than those of passenger genes. OGs also separate from passenger genes in that their rMSN values are higher and their rSMN and rNSM values are lower than those of passenger genes. **Panels B1, B2** show data only for candidate cancer genes present in the CG_SO^2SD^_SSI^2SD^ list (see **Supplementary file 6**). The positions of transcripts of the genes identified by Vogelstein *et al*., (2013) as oncogenes (OGs, large red balls) or tumor suppressor genes (TSGs, large blue balls) are highlighted. The positions of novel cancer gene transcripts validated in the present work are highlighted as large green balls.

The set of genes (4400 transcripts) with values that deviate from the mean by more than 1SD contained 77 OG transcripts, 132 TSG transcripts and an additional 347 CGC gene transcripts. The central cluster of genes, deviating from mean rSMN, rMSN and rNSM values by ≤1SD is hereafter referred to as PG_SO^r3_1SD^ (for Passenger Gene_Substitution Only deviating from mean rSMN, rMSN and rNSM values by ≤ 1SD).

The candidate cancer gene set defined by 2SD cut-off value (**Supplementary file 3**) is hereafter referred to as CG_SO^r3_2SD^ for Cancer Gene_Substitution Only deviating from mean rSMN, rMSN and rNSM values by more than 2SD (**Supplementary file 4**). This gene set has a total of 751 transcripts, containing transcripts of 35 OGs, 103 TSGs, an additional 80 CGC genes and 533 transcripts (derived from 448 genes) not found in the OG, TSG and CGC cancer gene lists (**Supplementary files 3** and **4**).

The mean parameters of TSGs differ markedly from those of passenger genes in as much as rNSM values of TSGs are higher but rSMN and rMSN values are lower (**Supplementary file 4**), reflecting the dominance of positive selection for inactivating mutations. In the case of the majority of OGs the rMSN values are higher and rNSM and rSMN values are lower than those of passenger genes (**Supplementary file 4**), reflecting positive selection for missense mutations and purifying selection avoiding nonsense mutations. Interestingly, some oncogenes have unusually high scores of rSMN (**Figures 4 A1, A2, Supplementary file 3**) suggesting that in these cases (e.g. *BCL2*) purifying selection dominates over positive selection for amino acid changing mutations.

As mentioned above, the candidate cancer gene set defined by a cut-off values of 2SD contains 533 transcripts (derived from 448 genes) not found in the OG, TSG or CGC lists. Since the majority of these genes have parameters that assign them to the clusters containing OGs or TSGs, they can be regarded as candidate oncogenes or tumor suppressor genes.

In these 3D scatter plots the existence of a group of genes that deviates from the clusters of passenger genes, OGs and TSGs is even more obvious (**Figure 4**): their high rSMN and low rMSN and rNSM values suggest that they experience purifying selection during tumor evolution, suggesting that they may be essential for the survival of tumors as oncogenes or tumor essential genes. The putative cancer genes listed in CG_SO^r3_2SD^ of **Supplementary file 3**, were subjected to further analyses to decide whether they qualify as candidate oncogenes, tumor suppressor genes or tumor essential genes (see section on **Analyses of candidate cancer gene sets**).

The three types of analyses for Substitutions Only, illustrated in **Figures 2-4** were also carried out for datasets in which both substitutions and subtle indels (Substitutions and Subtle Indels, SSI) were used, by merging nonsynonymous substitutions and short inframe indels in the category of mutations that introduce subtle changes in the amino acid sequence (indel_fM) and by including nonsense substitutions and short frame-shift indels in the category of mutations (indel_fN) that generate stop codons. (For details of these analyses see **Appendix 1).**

Comparison of the data obtained by SO and SSI analyses (**Supplementary file 3**) revealed that inclusion of indels has only minor influence on the separation of the clusters of PGs and CGs. For example, comparison of the lists of PGs identified with 1SD cut-off values for the three types of SO analyes (PG_SO^f_1SD^, PG_SO^r2_1SD^, PG_SO^r3_1SD^) with the corresponding lists identified for SSI analyses (PG_SSI^f_1SD^, PG_SSI^r2_1SD^, PG_SSI^r3_1SD^) revealed that the lists in the three types of SO/SSI pairs show more than 90% identity (**Supplementary file 5**). Similarly, the lists of CGs identified with 2SD cut-off values for the three types of SO analyses (CG_SO^f_2SD^, CG_SO^r2_2SD^, CG_SO^r3_2SD^) with the corresponding lists identified for SSI analyses (CG_SSI^f_2SD^, CG_SSI^r2_2SD^, CG_SSI^r3_2SD^) revealed that the three pairs of lists show 78%, 87% and 92% identity, respectively (**Supplementary file 5**).

## Discussion

### Analyses of candidate cancer gene sets

The parameters of the 1158 transcripts present in at least one of the various CG_SO^2SD^ lists and the 1333 transcripts present in at least one of the various CG_SSI^2SD^ lists (**Supplementary file 6**) differ from those of passenger genes in a way that assigns them to the clusters of genes positively selected for inactivating mutations or the clusters of genes positively selected for missense mutations or the clusters of negatively selected genes (see **Figure 2C, Figure 3 B1, B2** and **Figure 4 B1, B2**). To check the validity and predictive value of the assumption that the genes assigned to these clusters play significant roles in carcinogenesis we have selected a number of genes for further analyses from the 1457 transcripts present in the combined list (CG_SO^2SD^_SSI^2SD^) of candidate cancer genes (**Supplementary file 6**).

The selection of genes was based on three criteria: 1) the candidate gene is among the genes showing the strongest signals of selection characteristic of the given group; 2) the candidate gene is novel in the sense that it is not listed among the 145 ‘gold standard’ OG and TSG cancer genes of Vogelstein *et al*., [54] or among the 719 genes of CGC [59]; 3) there is substantial experimental information in the scientific literature on the given gene to permit the assessment of the validity of the assumption that it plays a role in carcinogenesis.

The genes discussed below include genes positively selected for truncating mutations, genes positively selected for missense mutations and negatively selected genes. In the main text we summarize only the major conclusions of our analyses; annotation of the individual genes is found in **Appendix 2**. We discuss examples of negatively selected genes in somewhat greater detail in the main text since they were inevitably missed by earlier studies that focussed on positive selection of driver mutations. We also discuss some examples of ‘false’ hits, i.e. cases where the mutation parameters deviate significantly from those of passenger genes, but this deviation is not due to selection.

### Novel cancer genes positively selected for truncating mutations

We have selected genes *B3GALT1, BMPR2, BRD7, ING1, MGA, PRRT2, RASA1, RNF128, SLC16A1, SPRED1, TGIF1, TNRC6B, TTK, ZNF276, ZC3H13, ZFP36L2, ZNF750* from the combined list of 1457 candidate transcripts (**Supplementary file 6)**, whose parameters deviate most significantly (by >2SD) from those of passenger genes, with the additional restriction that only genes with indel_rNSM > 0.125 (624 genes) were included (**Supplementary file 7)**, thereby removing the majority of passenger genes, oncogenes and tumor essential genes.

Annotation of the majority of these genes (*BMPR2, BRD7, ING1, MGA, PRRT2, RASA1, RNF128, SLC16A1, SPRED1, TGIF1, TNRC6B, ZC3H13, ZFP36L2* and *ZNF750*) has provided convincing evidence for their role in carcinogenesis as tumor suppressors. Interestingly, experimental evidence indicates that *TTK*, encoding dual specificity protein kinase TTK, is a proto-oncogene that may be converted to an oncogene by truncating mutations affecting its very C-terminal end, downstream of its kinase domain (for further details see **Appendix 2**). Our annotations suggest that *B3GALT1, ZNF276* are false positives whose apparent mutation pattern deviates significantly from those of passenger genes, but this deviation is not due to selection.

Based on functional annotation of the novel cancer genes identified and validated in the present work (see **Appendix 2**) we have assigned them to various cellular processes of cancer hallmarks in which they are involved (**Table 1**.).

**Table 1.**
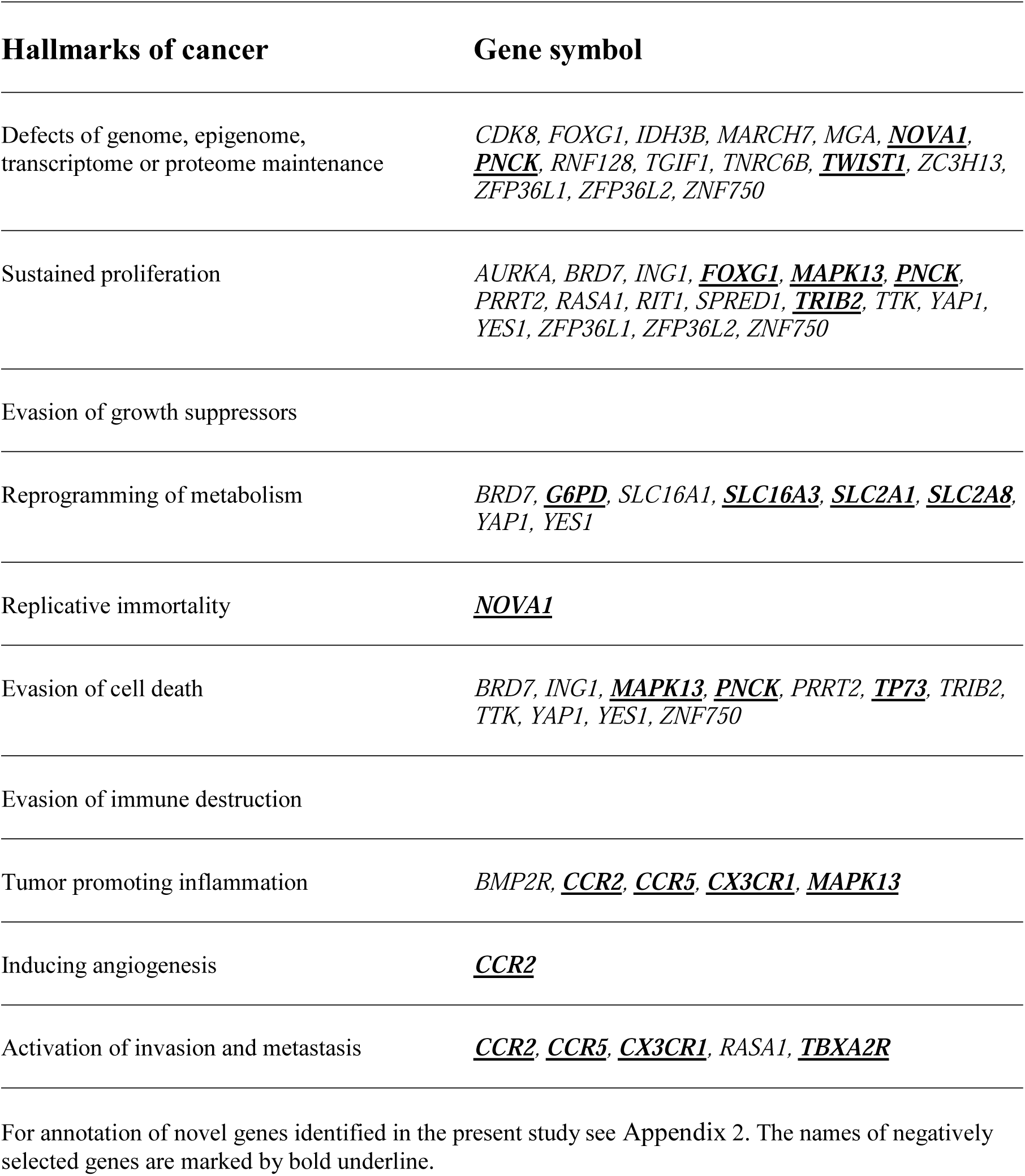
Assignment of novel positively or negatively selected cancer genes to key cellular processes of carcinogenesis

| Hallmarks of cancer | Gene symbol |
| --- | --- |
| Defects of genome, epigenome, transcriptome or proteome maintenance | *CDK8, FOXG1, IDH3B, MARCH7, MGA, <u>NOVA1</u>, <u>PNCK</u>, RNF128, TGIF1, TNRC6B, <u>TWIST1</u>, ZC3H13, ZFP36L1, ZFP36L2, ZNF750* |
| Sustained proliferation | *AURKA, BRD7, ING1, <u>FOXG1</u>, <u>MAPK13</u>, <u>PNCK</u>, PRRT2, RASA1, RIT1, SPRED1, <u>TRIB2</u>, TTK, YAP1, YES1, ZFP36L1, ZFP36L2, ZNF750* |
| Evasion of growth suppressors |  |
| Reprogramming of metabolism | *BRD7, <u>G6PD</u>, SLC16A1, <u>SLC16A3</u>, <u>SLC2A1</u>, <u>SLC2A8</u>, YAP1, YES1* |
| Replicative immortality | *<u>NOVA1</u>* |
| Evasion of cell death | *BRD7, ING1, <u>MAPK13</u>, <u>PNCK</u>, PRRT2, <u>TP73</u>, TRIB2, TTK, YAP1, YES1, ZNF750* |
| Evasion of immune destruction |  |
| Tumor promoting inflammation | *BMP2R, <u>CCR2</u>, <u>CCR5</u>, <u>CX3CR1</u>, <u>MAPK13</u>* |
| Inducing angiogenesis | *<u>CCR2</u>* |
| Activation of invasion and metastasis | *<u>CCR2</u>, <u>CCR5</u>, <u>CX3CR1</u>, RASA1, <u>TBXA2R</u>* |

For annotation of novel genes identified in the present study see Appendix 2. The names of negatively selected genes are marked by bold underline.

Comparison of the list of 624 genes present in this dataset (CG_SSI^2SD^ rNSM>0.125) with lists identified by others (**Supplementary file 7**) revealed that ∼60-100 of our candidate TSG-like genes are also found in several gene lists identified by analyses of somatic mutations of tumor tissues. Many of the genes selected for annotation are present in at least one of the candidate gene lists identified by others; the genes of *MGA, RASA1, TGIF1, ZFP36L2* and *ZNF750* are present in multiple cancer gene lists (**Supplementary file 7**). It is noteworthy, however, that *RNF128, SLC16A1, SPRED1, TNRC6B* and *TTK* are novel in that they are found only among the candidate cancer genes identified by forward genetic screens in mice [62] or among the genes whose expression changes in cancer [60].

### Novel cancer genes positively selected for missense mutations

We have selected genes *AURKA, CDK8, IDH3B, MARCH7, RIT1, YAP1, YES1* from the combined list of 1457 candidate transcripts, whose parameters deviate most significantly (by >2SD) from those of passenger genes (**Supplementary file 7)**, but only genes with rMSN>3.00 (440) were used, thereby removing the majority of passenger genes, tumor suppressor genes and tumor essential genes.

Annotation of these genes has confirmed that they play important roles in carcinogenesis as oncogenes. Three of these genes encode kinases (Aurora kinase A, also known as breast tumor-amplified kinase, cyclin-dependent kinase 8, tyrosine-protein kinase Yes, also known as proto-oncogene c-Yes) but unlike many other oncogenic kinases, these oncogenes do not show significant clustering of missense mutations. In fact, only in the case of *IDH3B* and *RIT1* did we observe clustering of missense mutations, indicating that recurrent mutation is not an obligatory property of proto-oncogenes.

Based on functional annotation of the novel oncogenes identified and validated in the present work (see **Appendix 2**) we have assigned them to various cellular processes of cancer hallmarks in which they are involved (**Table 1**.).

Comparison of this list of 440 genes (CG_SO^2SD^ rMSN>3.00) with the lists of cancer genes identified by others (**Supplementary file 7**) revealed that ∼60-100 of our candidate oncogene-like genes are present in cancer gene lists identified by analyses of somatic mutations of tumor tissues.

Out of the genes that we have selected for annotation only the *RIT1* gene has been identified by others as an oncogene, based on the analysis of somatic mutations (**Supplementary file 7**). *AURKA* and *IDH3B* are not found in any of the lists of cancer genes, whereas *CDK8, MARCH7, YAP1* and *YES1* are listed among the more than 9000 candidate cancer genes identified by forward genetic screens in mice [62]. Interestingly, TTK, identified as a gene positively selected for truncating mutations (see list CG_SSI^2SD^ rNSM > 0.125), but annotated as an oncogene, is also present in the list of genes positively selected for missense mutations (CG_SO^2SD^ rMSN>3.00).

### Negatively selected tumor essential genes

We have selected genes *CX3CR1, FOXG1, FOXP2, G6PD, MAPK13, MLLT3, NOVA1, PNCK, RUNX2, SLC16A3, SLC2A1, SLC2A8, TBP, TBXA2R, TP73, TRIB2* from the lists of cancer genes whose parameters deviate most significantly (by >2SD) from those of passenger genes, but only genes with rSMN > 0.5 (505 genes) were used to eliminate the majority of passenger genes, oncogenes and tumor suppressor genes.

Although our analyses have confirmed that in the majority of cases (*CX3CR1, FOXG1, G6PD, MAPK13, NOVA1, PNCK, SLC16A3, SLC2A1, SLC2A8, TBXA2R*,, *TP73, TRIB2)* the high synonymous to nonsynonymous and synonymous to nonsense mutation rates could be interpreted as evidence for purifying selection during tumor evolution, there were several examples (e.g. *DSPP, FOXP2, MLLT3, RUNX2, TBP*) where high synonymous to nonsynonymous and synonymous to nonsense mutation rates were found to reflect increased rates of synonymous substitution (due to the presence of mutation hotspots), rather than decreased rates of nonsynonymous and nonsense substitutions that could be due to purifying selection (for details see **Appendix 2**).

Annotation of the genes *CX3CR1, FOXG1, G6PD, MAPK13, NOVA1, PNCK, SLC16A3, SLC2A1, SLC2A8, TBXA2R, TP73, TRIB2* have confirmed that all of them play important roles in carcinogenesis (see **Appendix 2**) permitting their assignment to various cellular processes of cancer hallmarks (**Table 1**.). In harmony with the notion that negative selection reflects their essential role in tumor evolution, there is evidence that they fulfill pro-oncogenic functions by promoting cell proliferation (*FOXG1, MAPK13, PNCK, TRIB2*), evasion of cell death (*MAPK13, PNCK, TP73*), promoting replicative immortality (*NOVA1*), reprogramming of energy metabolism of cancer cells (*G6PD, SLC16A3, SLC2A1, SLC2A8*), inducing tumor promoting inflammation (*CCR2, CCR5, CX3CR1, MAPK13*) and invasion and metastasis (*CCR2, CCR5, CX3CR1, TBXA2R*).

Not surprisingly, none of these genes are present in the lists of positively selected driver genes (CG_SSI^2SD^ rNSM > 0.125 and CG_SO^2SD^ rMSN > 3.00, **Supplementary file 7**). It is noteworthy, however, that *G6PD, MAPK13, PNCK, SLC16A3* and *SLC2A1* are listed among the candidate cancer genes identified by forward genetic screens in mice [62].

Comparison of our list of 505 negatively selected genes (CG_SO^2SD^_rSMN>0.5) with that of Weghorn and Sunyaev [65] has revealed very little similarity (**Supplementary file 8).** Only 1 of the 147 genes identified by Weghorn and Sunyaev [65] is also present in the list of top-ranking negatively selected genes identified in the present study. A greater similarity was observed when we compared our list of negatively selected genes with that of Zhou *et al*. [50]: 32 of the 112 genes identified by Zhou *et al*., [50] are also present among the 505 negatively selected genes identified in the present work (**Supplementary file 8).** It is noteworthy that top-ranking genes present in both lists include the *TBP* gene, and the *MLLT3* gene. As discussed in **Appendix 2**, the apparent signals of negative selection (high synonymous to nonsynonymous rates) of genes like *DSPP, FOXP2, MLLT3, RUNX2* and *TBP* reflect the presence of mutation hotspots and not purifying selection. Zhou *et al.* [50] have noted that „some cancer genes also show negative selection in cancer genomes, such as the oncogene *MLLT3*”. Although they point out that „interestingly, *MLLT3* has recurrent synonymous mutations at amino acid positions 166 to 168” they do not seem to realize that this observation of recurrent silent substitutions (in a poly-Ser region of the protein) questions the validity of the claim that the unusually low nonsynonymous to synonymous rate is due to negative selection (for more detail see **Appendix 2**).

Otherwise, the lack of more extensive overlap of top-ranking negatively selected genes identified in the present study with those identified by others based on synonymous to nonsynonymous rates [50, 65] is probably due to the fact that in the present work we have combined multiple aspects of purifying selection and have increased the statistical power of our analyses by limiting our work to transcripts that have at least 100 somatic mutations.

It must also be pointed out that the conclusions drawn from earlier studies searching for signs of negative selection are highly controversial [50, 65, 67]. Zhou et al., [50] have succeeded in identifying a large set of negatively selected genes that were suggested to be important for the growth and survival of cancer cells. Although Weghorn and Sunyaev [65] have acknowledged that in their analyses the signals of purifying selection were exceedingly weak, they have identified a group of negatively selected genes that was enriched in cell-essential genes [66], leading them to propose that the major cause of negative selection during tumor evolution is the maintenance of genes that are responsible for basal cellular functions.

The third, much-publicized study, however, propagated the conclusion that negative selection has no role in tumor evolution [67-70]. Martincorena *et al.* [67] have argued that the practical absence of purifying selection during tumor evolution is due to the buffering effect of diploidy and functional redundancy of most cellular pathways.

The influence of functional redundancy on the essentiality of genes has been examined in a recent study [71]. The authors have used CRISPR score profiles of 558 genetically heterogeneous tumor cell lines and converted continuous values of gene CRISPR scores to binary essential and nonessential calls. These analyses have shown that 1014 genes belong to a category of ‘broadly essential genes’, i.e. these genes were found to be essential in at least 90% of the 558 cell lines. De Kegel and Ryan [71] have shown that, compared to singleton genes, paralogs are less frequently essential and that this is more evident when considering genes with multiple paralogs or with highly sequence-similar paralogs.

In order to assess the contribution of cell-essentiality to purifying selection during tumor evolution we have plotted various measures of negative selection of human genes as a function of their cell-essentiality scores determined by De Kegel and Ryan [71]. These analyses have shown that the cell-essentiality scores of negatively selected genes (CG_SO^2SD^ rSMN>0.5) are not significantly different from those of passenger genes (**Figure 5)**.

**Figure 5.**
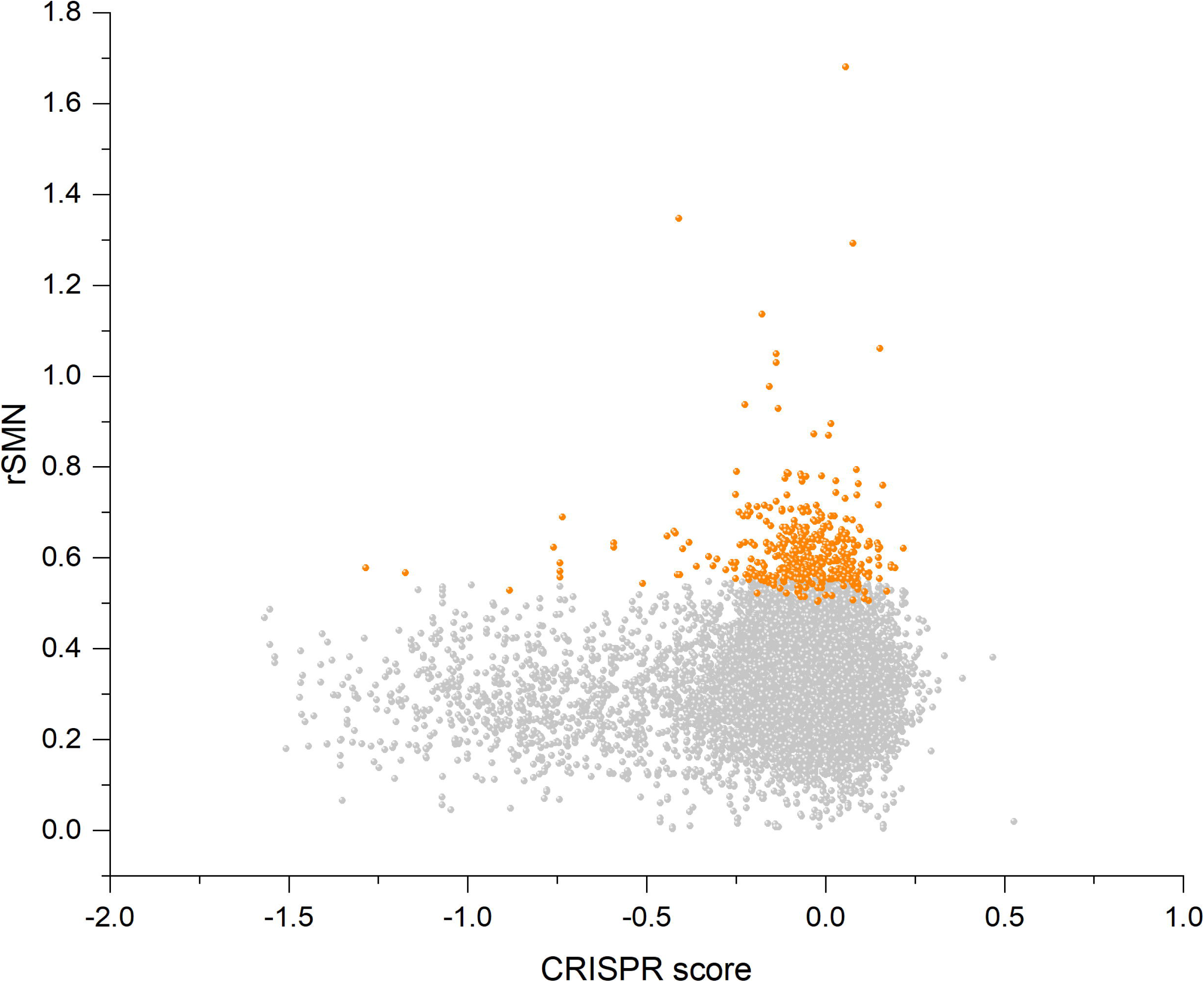
Lack of correlation between cell-essentiality scores of human genes and negative selection during tumor evolution. The figure shows the results of the analysis of transcripts containing at least 100 subtle, confirmed somatic, non polymorphic mutations from tumor tissues. The abscissa indicates the cell-essentiality score of the genes, the ordinate shows the rSMN parameters of the transcripts. Each ball represents a human transcript. Transcripts showing strongest signals of negatively selection (CG_SO^2SD^ rSMN>0.5) are represented by dark orange balls.

Comparison of CRISPR scores (−0.07665±0.17269) of the cluster of negatively selected genes of CG_SO^2SD^ rSMN>0.5) listed in **Supplementary file 8** with CRISPR scores (−0.09506±0.24168) of cluster of passenger genes (PG_SO^r3_1SD^) revealed that they are not significantly different (p>0.05), indicating that cell-essentiality *per se* does not explain purifying selection.

Comparison of the lists of negatively selected genes identified in the present work with the 1014 ‘broadly essential genes’ defined by De Kegel and Ryan [71] has revealed that there is practically no overlap between the two groups. Only 6 of the 1014 broadly essential genes are included in our list of negatively selected genes (**Supplementary file 8**). This observation also suggests that cell-essentiality defined by CRISPR scores determined experimentally on cell lines is not relevant for negative selection during tumor evolution *in vivo*.

Our analyses of cases of strong purifying selection suggest that it has more to do with a function specifically required by the tumor cell for its growth, survival and metastasis than with general basic cellular functions (**Table 1**). It is noteworthy in this respect, that the genes showing the strongest signals of negative selection include several plasma membrane receptor proteins (e.g. *ACKR3, CCR2, CCR5, CX3CR1, TBXA2R*) that cancer cells utilize to promote migration, invasion and metastasis (**Appendix 2**). Significantly, these proteins exert their biological functions (in cell migration, inflammation, angiogenesis etc.) primarily at the organism level, therefore their cell-essentiality scores may have little to do with their overall essentiality for tumor growth and metastasis. Inspection of the data of De Kegel and Ryan [71] shows that *ACKR3, CX3CR1, TBXA2R* were not assigned to the essential category in any of the 558 tumor cell lines tested.

Although negatively selected genes essential for carcinogenesis include proteins involved in cell-level processes, in that they promote cell proliferation (*FOXG1, MAPK13, PNCK, TRIB2*), evasion of cell death (*MAPK13, PNCK, TP73*), replicative immortality (e.g. *NOVA1*), or that they are crucial for the reprogramming of energy metabolism in cancer cells (e.g. *GAPD, SLC16A3, SLC2A1, SLC2A8*) their negative selection is unlikely to be a mere reflection of their basic cellular functions. Rather, it reflects the exceptional role of the corresponding cancer hallmarks (evasion of cell death, replicative immortality, reprogramming of metabolism) in carcinogenesis (**Figure 1**). In harmony with this conclusion *NOVA1, SLC16A3, SLC2A8, TP73* were assigned to the essential category by De Kegel and Ryan [71] in less than 10% of the 558 tumor cell lines tested. *SLC2A1* (glucose transporter 1) is an exception to some extent in as much as it was found to be cell-essential in 41% of the cell lines.

Significantly, several nutrient transporter protein genes (*SLC16A3, SLC2A1* and *SLC2A8)* were found among the genes showing strongest signs of purifying selection. The most likely explanation for their essentiality is that tumor cells have an increased demand for nutrients and this demand is met by enhanced cellular entry of nutrients through upregulation of specific transporters [72]. The uncontrolled cell proliferation of tumor cells involves major adjustments of energy metabolism in order to support cell growth and division in the hypoxic microenvironments in which they reside. Otto Warburg was the first to observe an anomalous characteristic of cancer cell energy metabolism: even in the presence of oxygen, cancer cells limit their energy metabolism largely to glycolysis, leading to a state that has been termed “aerobic glycolysis” [73, 74]. Cancer cells are known to compensate for the lower efficiency of ATP production through glycolysis than oxidative phosphorylation by upregulating glucose transporters, such as facilitated glucose transporter member 1, GLUT1 (encoded by the *SLC2A1* gene), thus increasing glucose import into the cytoplasm [75-77].

The markedly increased uptake of glucose has been documented in many human tumor types, by visualizing glucose uptake through positron emission tomography. The reliance of tumor cells on glycolysis is also supported by the hypoxia response system: under hypoxic conditions not only glucose transporters but also multiple enzymes of the glycolytic pathway are upregulated [75, 76, 78-80].

In our view, the central role of GLUT1 in cancer metabolism is reflected by the fact that the *SLC2A1* gene encoding this glucose transporter is among the genes that show the strongest signals of purifying selection. The key importance of GLUT1 in cancer may be illustrated by the fact that high levels of GLUT1 expression correlates with a poor overall survival and is associated with increased malignant potential, invasiveness and poor prognosis [81-83]. The strict requirement for GLUT1 in the early stages of mammary tumorigenesis highlights the potential for glucose restriction as a breast cancer preventive strategy [84]. The tumor essentiality of GLUT1 may also be illustrated by the fact that knockdown of GLUT1 inhibits cell glycolysis and proliferation and inhibits the growth of tumors [85]. In view of its essentiality for tumor growth, GLUT1 is a promising target for cancer therapy [86-88].

Recent studies suggest that the *YAP1-TEAD1-GLUT1* axis plays a major role in reprogramming of cancer energy metabolism by modulating glycolysis [89]. These authors have shown that *YAP1* and *TEAD1* are involved in transcriptional control of the glucose transporter *GLUT1*: whereas knockdown of *YAP1* inhibited glucose consumption, and lactate production of breast cancer cells, overexpression of GLUT1 restored glucose consumption and lactate production.

Besides GLUT1 another glucose transporter, GLUT8 (encoded by the *SLC2A8 gene*) also shows strong signals of negative selection, arguing for its importance in tumor survival. In harmony with this interpretation there is evidence that GLUT8 is overexpressed in and is required for proliferation and viability of tumors [90-91].

Due to abnormal conversion of pyruvic acid to lactic acid by tumor cells even under normoxia, the altered metabolism of glucose consuming tumors must rapidly efflux lactic acid to the microenvironment to maintain a robust glycolytic flux and to prevent poisoning themselves [92].

Survival and maintenance of the glycolytic phenotype of tumor cells is ensured by monocarboxylate transporter 4 (MCT4, encoded by the *SLC16A3* gene) that efficiently transports L-lactate out of the cell [72]. Significantly, MCT4, encoded by the *SLC16A3* gene also shows strong signals of negative selection, in harmony with its importance in tumor survival.

As high metabolic and proliferative rates in cancer cells lead to production of large amounts of lactate, extruding transporters are essential for the survival of cancer cells as illustrated by the fact that knockdown of MCT4 increased tumor-free survival and decreased *in vitro* proliferation rate of tumor cells [93]. Using a functional screen Baenke *et al*., [94] have also demonstrated that monocarboxylate transporter 4 is an important regulator of breast cancer cell survival: MCT4 depletion reduced the ability of breast cancer cells to grow, suggesting that it might be a valuable therapeutic target. In harmony with the essentiality of MCT4 for tumor growth, several studies indicate that expression of the hypoxia-inducible monocarboxylate transporter MCT4 is increased in tumors and its expression correlates with clinical outcome, thus it may serve as a valuable prognostic factor [95-97]. Consistent with the key importance of MCT4 for the survival of tumor cells, its selective inhibition to block lactic acid efflux appears to be a promising therapeutic strategy against highly glycolytic malignant tumors [98-101].

Interestingly, the thromboxane A2 receptor gene (*TBXA2R*) as well as several chemokine receptor protein genes (*CCR2, CCR5, CX3CR1)* were also found among the genes showing strong signs of purifying selection. The most likely explanation for their essentiality for tumor growth is that tumor cells rely on these receptors in various steps of invasion and metastasis (see **Appendix 2**). It is noteworthy in this respect that another member of the family of chemokine receptors, the atypical chemokine receptor 3, *ACKR3* is also among the genes showing very high values of rSMN, suggesting negative selection of missense and nonsense mutations (**Supplementary file 7)**. Significantly, *ACKR3* is a well-known oncogene, present in Tier 1 of the Cancer Gene Census. Several studies support the key role of *ACKR3* in tumor invasion and metastasis [102-107]. Since knock-down or pharmacological inhibition of *ACKR3* has been shown to reduce tumor invasion and metastasis, ACKR3 is a promising therapeutic target for the control of tumor dissemination (for further details see **Appendix 2**).

## Conclusions

One of the major goals of cancer research is to identify all ‘cancer genes’, i.e. genes that play a role in carcinogenesis. In the last two decades several types of approaches have been developed to achieve this goal, but the majority of the work focused on subtle mutations affecting the coding regions of genes. The implicit assumption of most of these studies was that a distinguishing feature of cancer genes is that they are positively selected for mutations that drive carcinogenesis. As a result of combined efforts the PCAWG driver list identifies a total of 722 protein-coding genes as cancer driver genes and 22 non-coding driver mutations [2, 28].

In a recent editorial, commenting on a suite of papers on the genetic causes of cancer, Nature has expressed the view that the core of the mission of cancer-genome sequencing projects — to provide a catalogue of driver mutations that could give rise to cancer — has been achieved [108]. It is noteworthy, however, that, although on average, cancer genomes were shown to contain 4–5 driver mutations, in around 5% of cases no drivers were identified in tumors [28]. As pointed out by the authors, this observation suggests that cancer driver discovery is not yet complete, possibly due to failure of the available bioinformatic algorithms. The authors have also suggested that tumors lacking driver mutations may be driven by mutations affecting cancer-associated genes that are not yet described for that tumor type, however, using driver discovery algorithms on tumors with no known drivers, no individual genes reached significance for point mutations [28].

In our view, these observations actually suggest that a rather large fraction of cancer genes remains to be identified. Assuming that tumors, on average, must have driver mutations affecting at least 4 or 5 cancer genes and that known and unknown cancer genes play similar roles in carcinogenesis, the observation that a 0.05 fraction of tumors has no known drivers (i.e. they are driven by 4-5 unknown cancer drivers) indicates that about half of the drivers is still unknown. If we assume that ∼50% of cancer genes is still unknown 3-6% (0.5^5^-0.5^4^, i.e. 0.03125-0.0625 fraction) of tumors is expected to lack any of the known driver genes, and to be driven by 4 or 5 unknown driver mutations. Since the list of known drivers used in the study of the ICGC/TCGA Pan-Cancer Analysis of Whole Genomes Consortium [28] comprises 722 driver genes, these observations suggest that hundreds of cancer driver genes remain to be identified.

In the present work we have used analyses that combined multiple types of signals of selection, permitting improved detection of positive and negative selection. Our analyses have identified a large number of novel positively selected cancer gene candidates, many of which could be shown to play significant roles in carcinogenesis as tumor suppressors and oncogenes. Significantly, our analyses have identified a major group of human genes that show signs of strong negative selection during tumor evolution, suggesting that the integrity of their function is essential for the growth and survival of tumor cells. Our analyses of representative members of negatively selected genes have confirmed that they play crucial pro-oncogenic roles in various cancer hallmarks (**Table 1**.). It is important to emphasize that a survey of the group of oncogenes and pro-oncogenic tumor essential genes reveals that they form a continuum in as much as there are numerous known oncogenes where negative selection also dominates (e.g. *ACKR3, BCL2*).

Although some groups have investigated the role of negative selection in tumor evolution earlier [50, 65, 67] the study that received the greatest attention has reached the conclusion that negative selection has no role in tumor evolution [67-70]. The data presented here contradict this conclusion.

We believe that the approach reported here will promote the identification of numerous novel tumor suppressor genes, oncogenes and pro-oncogenic genes that may serve as therapeutic targets.

## Methods

Cancer somatic mutation data were extracted from COSMIC v88, the Catalogue Of Somatic Mutations In Cancer, which includes single nucleotide substitutions and small insertions/deletions affecting the coding sequence of human genes. The downloaded file (CosmicMutantExport.tsv, release v88) contained data for 29415 transcripts (**Supplementary file 1**). For all subsequent analyses we have retained only transcripts containing mutations that were annotated under ‘Mutation description’ as substitution or subtle insertion/deletion. This dataset contained data for 29405 transcripts containing 6449721 mutations (substitution and short indels, SSI) and 29399 transcripts containing 6141650 substitutions only (SO).

To increase the statistical power of our analyses we have limited our work to transcripts that have at least 100 somatic mutations. Hereafter, unless otherwise indicated, our analyses refer to datasets containing transcripts with at least 100 somatic mutations. This limitation eliminated ∼38% of the transcripts that contain very few mutations but reduced the number of total mutations only by 9% (**Supplementary file 1**). It should be noted that this limitation increases the statistical power of our analyses but disfavors the identification of some negatively selected genes.

Since we were interested in the selection forces that operate during tumor, only confirmed somatic mutations were included in our analyses. In COSMIC such mutations are annotated under ‘Mutation somatic status’ as Confirmed Somatic, i.e. confirmed to be somatic in the experiment by sequencing both the tumor and a matched normal tissue from the same patient. As to ‘Sample Type, Tumor origin’: we have excluded mutation data from cell-lines, organoid-cultures, xenografts since they do not properly represent human tumor evolution at the organism level. We have found that by excluding cell lines we have eliminated many artifacts of spurious recurrent mutations caused by repeated deposition of samples taken from the same cell-line at different time-points. To eliminate the influence of polymorphisms on the conclusions we retained only somatic mutations flagged ‘n’ for SNPs. (**Supplementary file 1**).

In our datasets the numerical variables for sets of human genes were expressed as mean and standard deviation for each group of data. For each variable, the means for the various groups were compared using the t-test for independent samples. Statistical significance was set as a P value of <0.05.

We have used several approaches to estimate the contribution of silent, amino acid changing and truncating mutations to somatic mutations of human protein-coding genes during tumor evolution. We have used two major types of calculations: one in which we have restricted our analyses to single nucleotide substitutions (referred to as SO for ‘substitution only’) and a version in which we have also taken into account subtle indels (referred to as SSI for ‘substitutions and subtle indels’).

## Ethics declarations

### Ethics approval and consent to participate

Not applicable

### Consent for publication

Not applicable

## Availability of data and materials

The datasets supporting the conclusions of this article are included within the article and its additional files.

## Competing interests

The authors declare that they have no competing interests.

## Funding

LB, KK, MT and LP are supported by the GINOP-2.3.2-15-2016-00001 grant of the Hungarian National Research, Development and Innovation Office (NKFIH), OC is supported by the NVKP_16-1-2016-0005 grant of the Hungarian National Research, Development and Innovation Office (NKFIH).

## Contributions

LP designed the project and coordinated the research; LB, and LP performed the analyses of the somatic mutation datasets of tumor tissues; MT, KK, LB, OC and LP carried out the annotation of candidate cancer genes; LP wrote the original draft. LB, MT, KK and OC reviewed and edited the manuscript. All authors read, commented, and approved the final manuscript.

## Additional files

### Appendix 1

The file describes SSI analyses (Substitutions and Subtle Indels analyses) of silent, amino acid changing and truncating somatic mutations of human protein-coding genes of tumor tissues.

In SSI analyses subtle mutations affecting the coding sequences of protein coding genes were assigned to three categories: S, silent synonymous substitutions, M, merging nonsynonymous substitutions and short inframe indels that change but do not disrupt coding sequences, and N, merging nonsense substitutions and short frame-shift indels as both types of mutations lead eventually to stop codons that truncate the proteins.

### Appendix 2

The file contains description of selected genes identified in the present study displaying strong signatures of positive and/or negative selection and which are novel in the sense that they are not included in the most widely used cancer gene lists (Vogelstein *et al*. 2013; Sondka *et al*., 2018).

### Transparent reporting form

**Supplementary file 1.** Statistics of transcripts and subtle somatic mutations of human protein coding genes of the different datasets analyzed.

**Supplementary file 2.** SO (Substitution Only) and SSI (Substitutions and Subtle Indel) analyses of somatic mutations of transcripts of human protein coding genes. Transcripts of OGs (oncogenes) and TSGs (tumor suppressor genes) of the cancer gene list of Vogelstein *et al*. (2013) are highlighted by brick red and blue backgrounds, respectively. Transcripts of CGC (Cancer Gene Census) genes (Sondka *et al*., 2018) that do not correspond to OGs or TSGs of the cancer gene list of Vogelstein *et al*. (2013) are highlighted by yellow background.

**Supplementary file 3.** SO (Substitution Only) and SSI (Substitutions and Subtle Indel) analyses of somatic mutations of transcripts of human protein coding genes that have at least 100 confirmed somatic, non polymorphic mutations identified in tumor tosses. The table also contains lists of genes (PG_SO^f_1SD^, PG_SO^r2_1SD^, PG_SO^r3_1SD^, PG_SSI^f_1SD^, PG_SSI^r2_1SD^, PG_SSI^r3_1SD^) whose parameters deviate from the mean values by ≤1SD as well as lists of genes (CG_SO^f_1SD^, CG_SO^r2_1SD^, CG_SO^r3_1SD^, CG_SSI^f_1SD^, CG_SSI^r2_1SD^, CG_SSI^r3_1SD^) whose parameters deviate from the mean values by >1SD. Table also contains lists of genes (CG_SO^f_2SD^, CG_SO^r2_2SD^, CG_SO^r3_2SD^, CG_SSI^f_2SD^, CG_SSI^r2_2SD^, CG_SSI^r3_2SD^) whose parameters deviate from the mean values by >2SD as well as lists of genes (PG_SO^f_2SD^, PG_SO^r2_2SD^, PG_SO^r3_2SD^, PG_SSI^f_2SD^, PG_SSI^r2_2SD^, PG_SSI^r3_2SD^) whose parameters deviate from the mean values by <2SD. Transcripts of OGs (oncogenes) and TSGs (tumor suppressor genes) of the cancer gene list of Vogelstein *et al*. (2013) are highlighted by brick red and blue backgrounds, respectively. Transcripts of CGC (Cancer Gene Census) genes (Sondka *et al*., 2018) that do not correspond to OGs or TSGs of the cancer gene list of Vogelstein *et al*. (2013) are highlighted by yellow background.

**Supplementary file 4**. Statistics of the results of SO (Substitution Only) and SSI (Substitutions and Subtle Indel) analyses of the data presented in **Supplementary file 3.** The column marked ‘Expected’ indicates the parameters (highlighted by orange background) expected if we assume that the structure of the genetic code determines the probability of somatic substitutions.

**Supplementary file 5**. Comparison of the results of SO (Substitution Only) and SSI (Substitutions and Subtle Indel) analyses.

**Supplementary file 6.** Lists of genes (CG_SO^f_2SD^, CG_SO^r2_2SD^, CG_SO^r3_2SD^, CG_SSI^f_2SD^, CG_SSI^r2_2SD^, CG_SSI^r3_2SD^) whose parameters deviate from the mean values by >2SD. Transcripts of OGs (oncogenes) and TSGs (tumor suppressor genes) of the cancer gene list of Vogelstein *et al*. (2013) are highlighted by brick red and blue backgrounds, respectively. Transcripts of CGC (Cancer Gene Census) genes (Sondka *et al*., 2018) that do not correspond to OGs or TSGs of the cancer gene list of Vogelstein *et al*. (2013) are highlighted by yellow background.

**Supplementary file 7**. Comparison of the lists of genes in datasets CG_SSI^2SD^_rNSM> 0.125 and CG_SO^2SD^_rMSN>3.00 with the lists of cancer genes identified by others (VOG, Vogelstein *et al*., 2013; TAM, Tamborero *et al*. 2013; LAW, Lawrence *et al*. 2014; ABB, Abbott *et al*., 2015; TOR, Torrente *et al*. 2016; ZHO, Zhou *et al*. 2017; MAR, Martincorena *et al.* 2017; BAI, Bailey *et al*. 2018; SON, Sondka *et al*., 2018; ZHA, Zhao *et al*., 2019). Transcripts of OGs (oncogenes) and TSGs (tumor suppressor genes) of the cancer gene list of Vogelstein *et al*. (2013) are highlighted by brick red and blue backgrounds, respectively. Transcripts of CGC genes (SON, Sondka *et al*., 2018) that do not correspond to OGs or TSGs of the cancer gene list of Vogelstein *et al*. (2013) are highlighted by yellow background. Novel positively or negatively selected cancer genes validated in the present work are highlighted in dark green background.

**Supplementary file 8**. Comparison of the list of negatively selected genes, CG^2SD^_rSMN>0.5 with the lists of negatively selected genes (WEG and ZHOU), defined by Zhou et al., (2017), and Weghorn and Sunyaev (2017), respectively as well as the list of genes (De Kegel) identified by De Kegel and Ryan (2019) as broadly essential genes. Negatively selected genes discussed in detail in the present work are highlighted in dark green background.

## Notes

### Competing Interest Statement

The authors have declared no competing interest.

## References

1. Diederichs S, Bartsch L, Berkmann JC, Fröse K, Heitmann J, Hoppe C, Iggena D, Jazmati D, Karschnia P, Linsenmeier M, Maulhardt T, Möhrmann L, Morstein J et al. The dark matter of the cancer genome: aberrations in regulatory elements, untranslated regions, splice sites, non-coding RNA and synonymous mutations. EMBO Mol Med. 2016; 8:442–457.

2. Rheinbay E, Nielsen MM, Abascal F, Wala JA, Shapira O, Tiao G, Hornshøj H, Hess JM, Juul RI, Lin Z, Feuerbach L, Sabarinathan R, Madsen T. et al. PCAWG Consortium. Analyses of non-coding somatic drivers in 2,658 cancer whole genomes. Nature. 2020;578:102–111.

3. Heidenreich B, Rachakonda PS, Hemminki K, Kumar R. TERT promoter mutations in cancer development. Curr Opin Genet Dev. 2014; 24:30–37.

4. Li Y, Roberts ND, Wala JA, Shapira O, Schumacher SE, Kumar K, Khurana E, Waszak S, Korbel JO, Haber JE, Imielinski M; PCAWG Structural Variation Working Group, Weischenfeldt J, Beroukhim R, Campbell PJ; PCAWG Consortium. Patterns of somatic structural variation in human cancer genomes. Nature. 2020; 578:112–121.

5. Lengauer C, Kinzler KW, Vogelstein B. Genetic instabilities in human cancers. Nature. 1998; 396:643–649.

6. Cheng J, Demeulemeester J, Wedge DC, Vollan HKM, Pitt JJ, Russnes HG, Pandey BP, Nilsen G, Nord S, Bignell GR, White KP, Børresen-Dale AL, Campbell PJ et al. Pan-cancer analysis of homozygous deletions in primary tumours uncovers rare tumour suppressors. Nat Commun. 2017; 8:1221.

7. Beroukhim R, Mermel CH, Porter D, Wei G, Raychaudhuri S, Donovan J, Barretina J, Boehm JS, Dobson J, Urashima M, Mc Henry KT, Pinchback RM, Ligon AH et al. The landscape of somatic copy-number alteration across human cancers. Nature. 2010; 463:899–905.

8. Verhaak RGW, Bafna V, Mischel PS. Extrachromosomal oncogene amplification in tumour pathogenesis and evolution. Nat Rev Cancer. 2019; 19:283–288.

9. Haller F, Bieg M, Will R, Körner C, Weichenhan D, Bott A, Ishaque N, Lutsik P, Moskalev EA, Mueller SK, Bähr M, Woerner A, Kaiser B et al. Enhancer hijacking activates oncogenic transcription factor NR4A3 in acinic cell carcinomas of the salivary glands. Nat Commun. 2019; 10:368.

10. Yang YA, Yu J. EZH2, an epigenetic driver of prostate cancer. Protein Cell. 2013; 4:331–341.

11. Chen YC, Gotea V, Margolin G, Elnitski L. Significant associations between driver gene mutations and DNA methylation alterations across many cancer types. PLoS Comput Biol. 2017; 13:e1005840.

12. Di Domenico A, Wiedmer T, Marinoni I, Perren A. Genetic and epigenetic drivers of neuroendocrine tumours (NET). Endocr Relat Cancer. 2017; 24:R315–R334.

13. Roussel MF, Stripay JL. Epigenetic Drivers in Pediatric Medulloblastoma. Cerebellum. 2018; 17:28–36.

14. Chatterjee A, Rodger EJ, Eccles MR. Epigenetic drivers of tumourigenesis and cancer metastasis. Semin Cancer Biol. 2018; 51:149–159.

15. Pfeifer GP. Defining Driver DNA Methylation Changes in Human Cancer. Int J Mol Sci. 2018; 19. pii: E1166.

16. Van Tongelen A, Loriot A, De Smet C. Oncogenic roles of DNA hypomethylation through the activation of cancer-germline genes. Cancer Lett. 2017; 396:130–137.

17. Slack FJ, Chinnaiyan AM. The Role of Non-coding RNAs in Oncology. Cell. 2019; 179:1033–1055.

18. Dvinge H, Kim E, Abdel-Wahab O, Bradley RK. RNA splicing factors as oncoproteins and tumour suppressors. Nat Rev Cancer. 2016; 16:413–430.

19. Mofers A, Pellegrini P, Linder S, D’Arcy P. Proteasome-associated deubiquitinases and cancer. Cancer Metastasis Rev. 2017; 36:635–653.

20. Chen Y, Zhang Y, Guo X. Proteasome dysregulation in human cancer: implications for clinical therapies. Cancer Metastasis Rev. 2017; 36:703–716.

21. Voutsadakis IA. Proteasome expression and activity in cancer and cancer stem cells. Tumour Biol. 2017; 39:1010428317692248.

22. Ge Z, Leighton JS, Wang Y, Peng X, Chen Z, Chen H, Sun Y, Yao F, Li J, Zhang H, Liu J, Shriver CD, Hu H et al. Integrated Genomic Analysis of the Ubiquitin Pathway across Cancer Types. Cell Rep. 2018; 23:213-226.e3.

23. Bernassola F, Chillemi G, Melino G. HECT-Type E3 Ubiquitin Ligases in Cancer. Trends Biochem Sci. 2019;. pii: S0968-0004(19)30180-X.

24. Hanahan D, Weinberg RA. Hallmarks of cancer: the next generation. Cell. 2011; 144:646–674.

25. Maura F, Bolli N, Angelopoulos N, Dawson KJ, Leongamornlert D, Martincorena I, Mitchell TJ, Fullam A, Gonzalez S, Szalat R, Abascal F, Rodriguez-Martin B, Samur M, et al. Genomic landscape and chronological reconstruction of driver events in multiple myeloma. Nat Commun. 2019; 10:3835.

26. Vogelstein B, Kinzler KW. The Path to Cancer - Three Strikes and You’re Out. N Engl J Med. 2015; 373:1895–1898.

27. Anandakrishnan R, Varghese RT, Kinney NA, Garner HR. Estimating the number of genetic mutations (hits) required for carcinogenesis based on the distribution of somatic mutations. PLoS Comput Biol. 2019; 15:e1006881.

28. Campbell, P.J., Getz, G., Korbel, J.O. et al. Pan-Cancer Analysis of Whole Genomes Consortium. Pan-cancer analysis of whole genomes. Nature. 2020;578:82–93.

29. Gerstung M, Eriksson N, Lin J, Vogelstein B, Beerenwinkel N. The temporal order of genetic and pathway alterations in tumorigenesis. PLoS One. 2011; 6:e27136.

30. Bashashati A, Ha G, Tone A, Ding J, Prentice LM, Roth A, Rosner J, Shumansky K, Kalloger S, Senz J, Yang W, McConechy M et al. Distinct evolutionary trajectories of primary high-grade serous ovarian cancers revealed through spatial mutational profiling. J Pathol. 2013; 231:21–34.

31. Shain AH, Yeh I, Kovalyshyn I, Sriharan A, Talevich E, Gagnon A, Dummer R, North J, Pincus L, Ruben B, Rickaby W, D’Arrigo C, Robson A. et al. The Genetic Evolution of Melanoma from Precursor Lesions. N Engl J Med. 2015; 373:1926–1936.

32. Gerstung M, Jolly C, Leshchiner I, Dentro SC, Gonzalez S, Rosebrock D, Mitchell TJ, Rubanova Y, Anur P, Yu K, Tarabichi M, Deshwar A, Wintersinger J. et al.; PCAWG Evolution & Heterogeneity Working Group, Spellman PT, Wedge DC, Van Loo P; PCAWG Consortium. The evolutionary history of 2,658 cancers. Nature. 2020;578:122–128.

33. Parmigiani G, Boca S, Lin J, Kinzler KW, Velculescu V, Vogelstein B. Design and analysis issues in genome-wide somatic mutation studies of cancer. Genomics. 2009; 93:17–21.

34. Meyerson M, Gabriel S, Getz G. Advances in understanding cancer genomes through second-generation sequencing. Nat Rev Genet. 2010; 11:685–696.

35. Michaelson JJ, Shi Y, Gujral M, Zheng H, Malhotra D, Jin X, Jian M, Liu G, Greer D, Bhandari A, Wu W, Corominas R, Peoples A, et al. Whole-genome sequencing in autism identifies hot spots for de novo germline mutation. Cell. 2012; 151:1431–1442.

36. Rogozin IB, Pavlov YI. Theoretical analysis of mutation hotspots and their DNA sequence context specificity. Mutat Res. 2003; 544:65–85.

37. Carter H. Mutation hotspots may not be drug targets. Science. 2019; 364:1228–1229.

38. Buisson R, Langenbucher A, Bowen D, Kwan EE, Benes CH, Zou L, Lawrence MS. Passenger hotspot mutations in cancer driven by APOBEC3A and mesoscale genomic features. Science. 2019; 364 pii: eaaw2872.

39. Poulos RC, Wong YT, Ryan R, Pang H, Wong JWH. Analysis of 7,815 cancer exomes reveals associations between mutational processes and somatic driver mutations. PLoS Genet. 2018; 14:e1007779.

40. Schuster-Böckler B, Lehner B. Chromatin organization is a major influence on regional mutation rates in human cancer cells. Nature. 2012; 488:504–507.

41. Gonzalez-Perez A, Sabarinathan R, Lopez-Bigas N. Local Determinants of the Mutational Landscape of the Human Genome. Cell. 2019; 177:101–114.

42. Polak P, Karlić R, Koren A, Thurman R, Sandstrom R, Lawrence M, Reynolds A, Rynes E, Vlahovicek K, Stamatoyannopoulos JA, Sunyaev SR. Cell-of-origin chromatin organization shapes the mutational landscape of cancer. Nature. 2015; 518:360–364.

43. Salvadores M, Mas-Ponte D, Supek F. Passenger mutations accurately classify human tumors. PLoS Comput Biol. 2019; 15:e1006953.

44. Lawrence MS, Stojanov P, Polak P, Kryukov GV, Cibulskis K, Sivachenko A, Carter SL, Stewart C, Mermel CH, Roberts SA, Kiezun A, Hammerman PS, McKenna A et al. Mutational heterogeneity in cancer and the search for new cancer-associated genes. Nature. 2013; 499:214–218.

45. Lawrence MS, Stojanov P, Mermel CH, Robinson JT, Garraway LA, Golub TR, Meyerson M, Gabriel SB, Lander ES, Getz G. Discovery and saturation analysis of cancer genes across 21 tumour types. Nature. 2014; 505:495–501.

46. Kaminker JS, Zhang Y, Watanabe C, Zhang Z. CanPredict: a computational tool for predicting cancer-associated missense mutations. Nucleic Acids Res. 2007; 35(Web Server issue):W595–598.

47. Carter H, Chen S, Isik L, Tyekucheva S, Velculescu VE, Kinzler KW, Vogelstein B, Karchin R. Cancer-specific high-throughput annotation of somatic mutations: computational prediction of driver missense mutations. Cancer Res. 2009; 69:6660–6667.

48. Nussinov R, Jang H, Tsai CJ, Cheng F. Precision medicine and driver mutations: Computational methods, functional assays and conformational principles for interpreting cancer drivers. PLoS Comput Biol. 2019; 15:e1006658.

49. Youn A, Simon R. Identifying cancer driver genes in tumor genome sequencing studies. Bioinformatics. 2011; 27:175–181.

50. Zhou Z, Zou Y, Liu G, Zhou J, Wu J, Zhao S, Su Z, Gu X. Mutation-profile-based methods for understanding selection forces in cancer somatic mutations: a comparative analysis. Oncotarget. 2017; 8:58835–58846.

51. Supek F, Miñana B, Valcárcel J, Gabaldón T, Lehner B. Synonymous mutations frequently act as driver mutations in human cancers. Cell. 2014; 156:1324–1335.

52. Hurst LD, Batada NN. Depletion of somatic mutations in splicing-associated sequences in cancer genomes. Genome Biol. 2017; 18:213.

53. Sharma Y, Miladi M, Dukare S, Boulay K, Caudron-Herger M, Groß M, Backofen R, Diederichs S. A pan-cancer analysis of synonymous mutations. Nat Commun. 2019; 10:2569.

54. Vogelstein B, Papadopoulos N, Velculescu VE, Zhou S, Diaz LA Jr, Kinzler KW. Cancer genome landscapes. Science. 2013; 339:1546–1558.

55. Tamborero D, Gonzalez-Perez A, Perez-Llamas C, Deu-Pons J, Kandoth C, Reimand J, Lawrence MS, Getz G, Bader GD, Ding L, Lopez-Bigas N. Comprehensive identification of mutational cancer driver genes across 12 tumor types. Sci Rep. 2013; 3:2650.

56. Bailey MH, Tokheim C, Porta-Pardo E, Sengupta S, Bertrand D, Weerasinghe A, Colaprico A, Wendl MC, Kim J, Reardon B, Kwok-Shing Ng P, Jeong KJ, Cao S et al. Comprehensive Characterization of Cancer Driver Genes and Mutations. Cell. 2018; 174:1034–1035.

57. Zhao S, Liu J, Nanga P, Liu Y, Cicek AE, Knoblauch N, He C, Stephens M, He X. Detailed modeling of positive selection improves detection of cancer driver genes. Nat Commun. 2019; 10:3399.

58. Tate JG, Bamford S, Jubb HC, Sondka Z, Beare DM, Bindal N, Boutselakis H, Cole CG, Creatore C, Dawson E, Fish P, Harsha B, Hathaway C, et al. COSMIC: the Catalogue Of Somatic Mutations In Cancer. Nucleic Acids Res. 2019; 47:D941–D947.

59. Sondka Z, Bamford S, Cole CG, Ward SA, Dunham I, Forbes SA. The COSMIC Cancer Gene Census: describing genetic dysfunction across all human cancers. Nat Rev Cancer. 2018; 18:696–705.

60. Torrente A, Lukk M, Xue V, Parkinson H, Rung J, Brazma A. Identification of Cancer Related Genes Using a Comprehensive Map of Human Gene Expression. PLoS One. 2016; 11:e0157484.

61. Calabrese C, Davidson NR, Demircioğlu D, Fonseca NA, He Y, Kahles A, Lehmann KV, Liu F, Shiraishi Y, Soulette CM, Urban L, Greger L, Li S. et al. PCAWG Consortium. Genomic basis for RNA alterations in cancer. Nature. 2020;578:129–136.

62. Abbott KL, Nyre ET, Abrahante J, Ho YY, Isaksson Vogel R, Starr TK. The Candidate Cancer Gene Database: a database of cancer driver genes from forward genetic screens in mice. Nucleic Acids Res. 2015; 43:D844–848.

63. Futreal PA, Coin L, Marshall M, Down T, Hubbard T, Wooster R, Rahman N, Stratton MR. A census of human cancer genes. Nat Rev Cancer. 2004; 4:177–183.

64. Patthy, L. (1999) Protein Evolution, 2nd ed.; Blackwell Publishing Ltd.

65. Weghorn D, Sunyaev S. Bayesian inference of negative and positive selection in human cancers. Nat Genet. 2017; 49:1785–1788.

66. Wang T, Birsoy K, Hughes NW, Krupczak KM, Post Y, Wei JJ, Lander ES, Sabatini DM. Identification and characterization of essential genes in the human genome. Science. 2015; 350:1096–1101.

67. Martincorena I, Raine KM, Gerstung M, Dawson KJ, Haase K, Van Loo P, Davies H, Stratton MR, Campbell PJ. Universal Patterns of Selection in Cancer and Somatic Tissues. Cell. 2017; 171:1029-1041.e21.

68. Bakhoum SF, Landau DA. Cancer Evolution: No Room for Negative Selection. Cell. 2017; 171:987–989.

69. Koch L. Cancer genomics: The driving force of cancer evolution. Nat Rev Genet. 2017; 18:703.

70. Vitale I, Galluzzi L. Everybody In! No Bouncers at Tumor Gates. Trends Genet. 2018; 34:85–87.

71. De Kegel B, Ryan CJ. Paralog buffering contributes to the variable essentiality of genes in cancer cell lines. PLoS Genet. 2019;15:e1008466.

72. Ganapathy V, Thangaraju M, Prasad PD. Nutrient transporters in cancer: relevance to Warburg hypothesis and beyond. Pharmacol Ther. 2009; 121:29–40.

73. Warburg O. On respiratory impairment in cancer cells. Science. 1956; 124:269–270.

74. Warburg O. On the origin of cancer cells. Science. 1956; 123:309–314.

75. Jones RG. Thompson C.B. Tumor suppressors and cell metabolism: a recipe for cancer growth. Genes Dev. 2009; 23:537–548.

76. DeBerardinis RJ. Lum JJ. Hatzivassiliou G. Thompson CB. The biology of cancer: Metabolic reprogramming fuels cell growth and proliferation. Cell Metab. 2008; 7:11–20.

77. Hsu P.P. Sabatini D.M. Cancer cell metabolism: Warburg and beyond. Cell. 2008; 134:703–707.

78. Semenza GL. HIF-1: upstream and downstream of cancer metabolism. Curr. Opin. Genet. Dev. 2010; 20:51–56.

79. Semenza GL. Defining the role of hypoxia-inducible factor 1 in cancer biology and therapeutics. Oncogene. 2010; 29:625–634.

80. Kroemer G. Pouyssegur J. Tumor cell metabolism: Cancer’s Achilles’ heel. Cancer Cell. 2008; 13:472–482.

81. Wang J, Ye C, Chen C, Xiong H, Xie B, Zhou J, Chen Y, Zheng S, Wang L. Glucose transporter GLUT1 expression and clinical outcome in solid tumors: a systematic review and meta-analysis. Oncotarget. 2017; 8:16875–16886.

82. Deng Y, Zou J, Deng T, Liu J. Clinicopathological and prognostic significance of GLUT1 in breast cancer: A meta-analysis. Medicine (Baltimore). 2018; 97:e12961.

83. de Castro TB, Mota AL, Bordin-Junior NA, Neto DS, Zuccari DAPC. Immunohistochemical Expression of Melatonin Receptor MT1 and Glucose Transporter GLUT1 in Human Breast Cancer. Anticancer Agents Med Chem. 2018; 18:2110–2116.

84. Wellberg EA, Johnson S, Finlay-Schultz J, Lewis AS, Terrell KL, Sartorius CA, Abel ED, Muller WJ, Anderson SM. The glucose transporter GLUT1 is required for ErbB2-induced mammary tumorigenesis. Breast Cancer Res. 2016; 18:131.

85. Xiao H, Wang J, Yan W, Cui Y, Chen Z, Gao X, Wen X, Chen J. GLUT1 regulates cell glycolysis and proliferation in prostate cancer. Prostate. 2018; 78:86–94.

86. Shibuya K, Okada M, Suzuki S, Seino M, Seino S, Takeda H, Kitanaka C. Targeting the facilitative glucose transporter GLUT1 inhibits the self-renewal and tumor-initiating capacity of cancer stem cells. Oncotarget. 2015; 6:651–661.

87. Noguchi C, Kamitori K, Hossain A, Hoshikawa H, Katagi A, Dong Y, Sui L, Tokuda M, Yamaguchi F. D-Allose Inhibits Cancer Cell Growth by Reducing GLUT1 Expression. Tohoku J Exp Med. 2016; 238:131–141.

88. Chen Q, Meng YQ, Xu XF, Gu J. Blockade of GLUT1 by WZB117 resensitizes breast cancer cells to adriamycin. Anticancer Drugs. 2017; 28:880–887.

89. Lin C, Xu X. YAP1-TEAD1-Glut1 axis dictates the oncogenic phenotypes of breast cancer cells by modulating glycolysis. Biomed Pharmacother. 2017; 95:789–794.

90. Goldman NA, Katz EB, Glenn AS, Weldon RH, Jones JG, Lynch U, Fezzari MJ, Runowicz CD, Goldberg GL, Charron MJ. GLUT1 and GLUT8 in endometrium and endometrial adenocarcinoma. Mod Pathol. 2006; 19:1429–1436.

91. McBrayer SK, Cheng JC, Singhal S, Krett NL, Rosen ST, Shanmugam M. Multiple myeloma exhibits novel dependence on GLUT4, GLUT8, and GLUT11: implications for glucose transporter-directed therapy. Blood. 2012; 119:4686–4697.

92. Mathupala SP, Colen CB, Parajuli P, Sloan AE. Lactate and malignant tumors: a therapeutic target at the end stage of glycolysis (Review). J Bioenerg Biomembr. 2007; 39:73–77.

93. Andersen AP, Samsøe-Petersen J, Oernbo EK, Boedtkjer E, Moreira JMA, Kveiborg M, Pedersen SF. The net acid extruders NHE1, NBCn1 and MCT4 promote mammary tumor growth through distinct but overlapping mechanisms. Int J Cancer. 2018; 142:2529–2542.

94. Baenke F, Dubuis S, Brault C, Weigelt B, Dankworth B, Griffiths B, Jiang M, Mackay A, Saunders B, Spencer-Dene B, Ros S, Stamp G, Reis-Filho JS et al. Functional screening identifies MCT4 as a key regulator of breast cancer cell metabolism and survival. J Pathol. 2015; 237:152–165.

95. Witkiewicz AK, Whitaker-Menezes D, Dasgupta A, Philp NJ, Lin Z, Gandara R, Sneddon S, Martinez-Outschoorn UE, Sotgia F, Lisanti MP. Using the “reverse Warburg effect” to identify high-risk breast cancer patients: stromal MCT4 predicts poor clinical outcome in triple-negative breast cancers. Cell Cycle. 2012; 11:1108–1117.

96. Doyen J, Trastour C, Ettore F, Peyrottes I, Toussant N, Gal J, Ilc K, Roux D, Parks SK, Ferrero JM, Pouysségur J. Expression of the hypoxia-inducible monocarboxylate transporter MCT4 is increased in triple negative breast cancer and correlates independently with clinical outcome. Biochem Biophys Res Commun. 2014; 451:54–61.

97. Baek G, Tse YF, Hu Z, Cox D, Buboltz N, McCue P, Yeo CJ, White MA, DeBerardinis RJ, Knudsen ES, Witkiewicz AK. MCT4 defines a glycolytic subtype of pancreatic cancer with poor prognosis and unique metabolic dependencies. Cell Rep. 2014; 9:2233–2249.

98. Choi SY, Xue H, Wu R, Fazli L, Lin D, Collins CC, Gleave ME, Gout PW, Wang Y. The MCT4 Gene: A Novel, Potential Target for Therapy of Advanced Prostate Cancer. Clin Cancer Res. 2016; 22:2721–2733.

99. Todenhöfer T, Seiler R, Stewart C, Moskalev I, Gao J, Ladhar S, Kamjabi A, Al Nakouzi N, Hayashi T, Choi S, Wang Y, Frees S, Daugaard M. et al. Selective Inhibition of the Lactate Transporter MCT4 Reduces Growth of Invasive Bladder Cancer. Mol Cancer Ther. 2018; 17:2746–2755.

100. Choi SYC, Ettinger SL, Lin D, Xue H, Ci X, Nabavi N, Bell RH, Mo F, Gout PW, Fleshner NE, Gleave ME, Collins CC, Wang Y. Targeting MCT4 to reduce lactic acid secretion and glycolysis for treatment of neuroendocrine prostate cancer. Cancer Med. 2018 7:3385–3392.

101. Zhao Y, Li W, Li M, Hu Y, Zhang H, Song G, Yang L, Cai K, Luo Z. Targeted inhibition of MCT4 disrupts intracellular pH homeostasis and confers self-regulated apoptosis on hepatocellular carcinoma. Exp Cell Res. 2019:111591.

102. Li XX, Zheng HT1, Huang LY, Shi DB, Peng JJ, Liang L, Cai SJ. Silencing of CXCR7 gene represses growth and invasion and induces apoptosis in colorectal cancer through ERK and β-arrestin pathways. Int J Oncol. 2014; 45:1649–5167.

103. Stacer AC, Fenner J, Cavnar SP, Xiao A, Zhao S, Chang SL, Salomonnson A, Luker KE, Luker GD. Endothelial CXCR7 regulates breast cancer metastasis. Oncogene. 2016; 35:1716–1724.

104. Zhao ZW, Fan XX, Song JJ, Xu M, Chen MJ, Tu JF, Wu FZ, Zhang DK, Liu L, Chen L, Ying XH, Ji JS. ShRNA knock-down of CXCR7 inhibits tumour invasion and metastasis in hepatocellular carcinoma after transcatheter arterial chemoembolization. J Cell Mol Med. 2017; 21:1989–1999.

105. Puddinu V, Casella S, Radice E, Thelen S, Dirnhofer S, Bertoni F, Thelen M. ACKR3 expression on diffuse large B cell lymphoma is required for tumor spreading and tissue infiltration. Oncotarget. 2017; 8:85068–85084.

106. Melo RCC, Ferro KPV, Duarte ADSS, Olalla Saad ST. CXCR7 participates in CXCL12-mediated migration and homing of leukemic and normal hematopoietic cells. Stem Cell Res Ther. 2018; 9:34.

107. Qian T, Liu Y, Dong Y, Zhang L, Dong Y, Sun Y, Sun D. CXCR7 regulates breast tumor metastasis and angiogenesis in vivo and in vitro. Mol Med Rep. 2018; 17:3633–3639.

108. EDITORIAL. The era of massive cancer sequencing projects has reached a turning point. The future of cancer genomics lies in the clinic. Nature. 2000; 578, 7–8.

