## Appendix 1 for "Use of signals of positive and negative selection to distinguish cancer genes and passenger genes"

### Analyses of somatic substitutions and subtle indel mutations of human protein-coding genes of tumor tissues

We have used two major types of analyses of silent, amino acid changing and truncating somatic mutations of human protein-coding genes of tumor tissues: one in which we have restricted our analyses to single nucleotide substitutions (SO or 'substitution only' analyes, for details, see main text).

Here we describe the analyes that also take into account subtle indels (SSI or 'substitutions and subtle indels' analyses). In these analyses subtle mutations affecting the coding sequences of protein coding genes were assigned to three categories: SIL, silent synonymous substitutions, MIS, merging nonsynonymous substitutions and short inframe indels that alter but do not disrupt coding sequence, and NON, merging nonsense substitutions and short frame-shift indels as both types of mutations lead eventually to stop codons that truncate the protein. Unless otherwise idicated, we have used datasets containing transcripts with at least 100 confirmed somatic, non polymorphic mutations identified in tumor tisses.

We have used several approaches to analyze the contribution of silent, amino acid changing and truncating mutations to somatic mutations of human protein-coding genes during tumor evolution.

In the simplest case we have calculated for each transcript the fraction of somatic mutations that could be assigned to the synonymous (indel_fS), nonsynonymous (indel_fM) and nonsense mutation (indel_fN) category.

Our analyses have shown that in the 3D representation of SSI mutations (see ***Appendix 1 – figure 1, Panel A***) genes are present in a cluster characterized by fraction values of 0.24082±0.06203, 0.70086±0.05701 and 0.05832±0.04151 for indel_fS, indel_fM and indel_fN category, respectively. The mean values for indel_fS, indel_fM and indel_fN in this cluster are very similar to those observed for fS, fM and fN in SO analyses (**Supplementary file 4**), consistent with the observation that in the dataset containing transcripts with at least 100 confirmed somatic, non polymorphic mutations identified in tumor tissues subtle indels are much rarer than single nucleotide substitutions (**Supplementary file 1**).

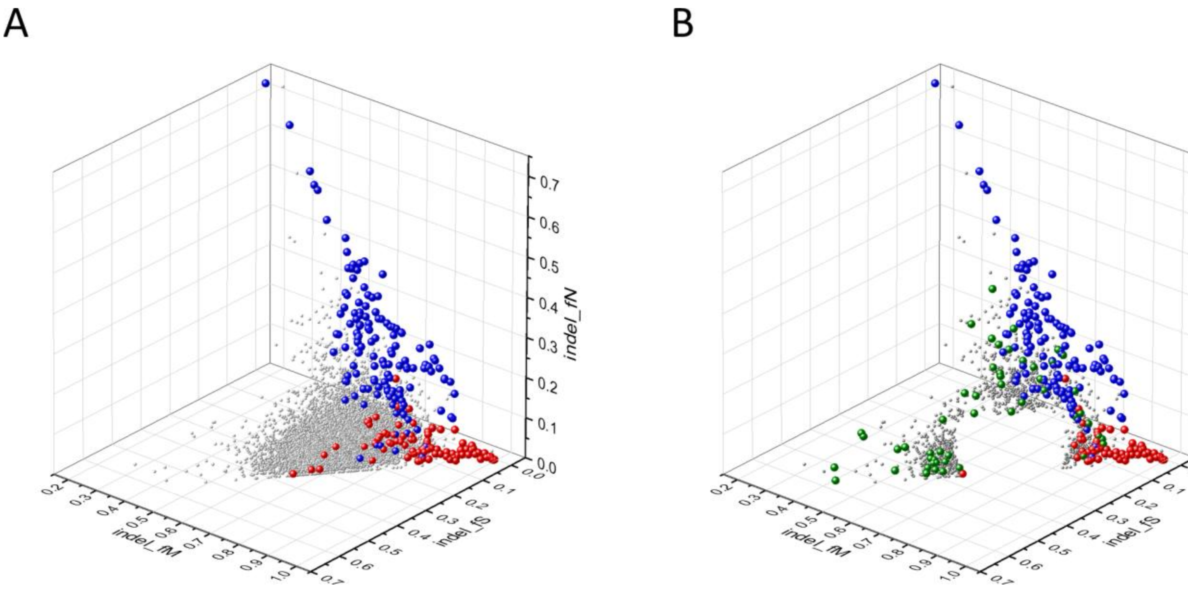


**Appendix 1 – figure 1. Analyses of indel_fS, indel_fM and indel_fN parameters of human protein-coding genes of tumor tissues.** The figure shows the results of the analysis of 13930 transcripts containing at least 100 subtle, confirmed somatic non-polymorphic mutations from tumor tissues. Axes *x*, *y* and *z* represent the fractions of somatic mutations that are assigned to the indel_fS, indel_fM and indel_fN categories. In **Panel A** each ball represents a human transcript; note that the majority of human genes are present in a dense cluster. The positions of transcripts of the genes defined by Vogelstein *et al*., (2013) as oncogenes (OGs, large red balls) or tumor suppressor genes (TSGs, large blue balls) are highlighted. It is noteworthy that these driver genes separate significantly from the central cluster and from each other: OGs have an increased fraction of indel_fM, whereas TSGs have markedly increased fraction of indel_fN. **Panel B** shows data only for candidate cancer genes present in the CG_SO$^{2SD}$_SSI$^{2SD}$ list (see **Supplementary file 6**). The positions of transcripts of the genes identified by Vogelstein *et al*., (2013) as oncogenes (OGs, large red balls) or tumor suppressor genes (TSGs, large blue balls) are highlighted. The positions of novel cancer gene transcripts validated in the present work are highlighted as large green balls.

It is noteworthy, however, that the pattern of indel_fS, indel_fM and indel_fN of the best known cancer genes (Vogelstein *et al*., 2013) deviates significantly from that characteristic of the majority of human genes (see ***Appendix 1 – figure 1, Panel A***). The values for OGs show a marked incease in indel_fM, reflecting positive selection for missense mutations, whereas the values for TSGs show significant increase in indel_fN, reflecting primarily positive selection for truncating nonsense mutations (**Supplementary file 4**).

The set of genes (6139 transcripts) with values that deviate from mean values of indel_fS, indel_fM and indel_fN by more than 1SD have also included the majority of OGs and TSGs (only 5 OG and 1 TSG transcripts remained in the central cluster). It is noteworthy that the 6139 transcripts also contained the vast majority (443 out of 748) of the transcripts of CGC genes, suggesting that the mutation pattern of most CGC genes also deviates significantly from that of passenger genes (**Supplementary file 4**). The genes in the central cluster (**Supplementary file 4**) is hereafter referred to as PG_SSI$^{f_1SD}$ (for Passenger Gene_Substitution and Subtle Indels deviating from mean indel_fS, indel_fM and indel_fN values by ≤1SD).

The set of genes (1211 transcripts) with values that deviate from mean values of indel_fS, indel_fM and indel_fN by more than 2SD included 62 OG and 123 TSG driver gene transcripts (see ***Appendix 1 – figure 1, Panel B***). Using this more stringent cut-off value the number of CGC genes identified in the 1211 transcripts was reduced to 153 out of 748 (**Supplementary file 4**). The non-passenger gene set defined by 2SD cut-off value is hereafter referred to as $CG\_SSI^{f\_2SD}$ for Cancer Gene_ Substitution and Subtle Indels deviating from mean indel_fS, indel_fM and indel_fN values by more than 2SD (**Supplementary file 4**).

The 1211 transcripts in the gene set of $CG\_SSI^{f\_2SD}$ has 873 transcripts not found in the OG, TSG and CGC cancer gene lists (**Supplementary file 3**). Since the majority of these 873 transcripts (derived from 743 genes) have parameters that assign them to the OG or TSG clusters, we assume that they also qualify as candidate oncogenes or tumor suppressor genes. There is, however, a third group of genes that deviate from both the central passenger gene cluster and the clusters of OGs and TSGs: their high indel_fS and low indel_fM and indel_fN values suggest that they experience purifying selection during tumor evolution, suggesting that they may correspond to tumor essential genes important for the growth and survival of tumors. The 743 putative cancer genes listed in $CG\_SSI^{f\_2SD}$ of **Supplementary file 3**, were subjected to further analyses to decide whether they qualify as candidate oncogenes, tumor suppressor genes, tumor essential genes or the deviation of their mutation pattern from those of passenger genes is not the result of natural selection. For some typical examples of these analyses see **Appendix 2**.

Known cancer genes (OGs and TSGs) also separate from the majority of human genes in 3D representations of parameters indel_rSM, indel_rNM, indel_rNS defined as the ratio of indel_fS/indel_fM, indel_fN/indel_fM, indel_fN/indel_fS, respectively (see ***Appendix 1 – figure 2***). In these representations (see ***Appendix 1 – figure 2, Panels A1, A2***) OGs separate from the central cluster in having significantly lower indel_rSM and indel_rNM values, whereas TSGs had significantly higher indel_rNS and indel_rNM values than those of the central cluster.

The set of genes (4518 transcripts) with values that deviate from the mean by more than 1SD contained 78 OG transcripts, 132 TSG transcripts and 368 CGC gene transcripts (**Supplementary file 4**). The central cluster of genes (that deviate from mean rSM, rNM and rNS values by ≤ 1SD) is hereafter referred to as $PG\_SSI^{r2\_1SD}$ (for Passenger Gene_Substitution and Subtle Indels deviating from mean indel_rSM, indel_rNM, indel_rNS values by ≤ 1SD).

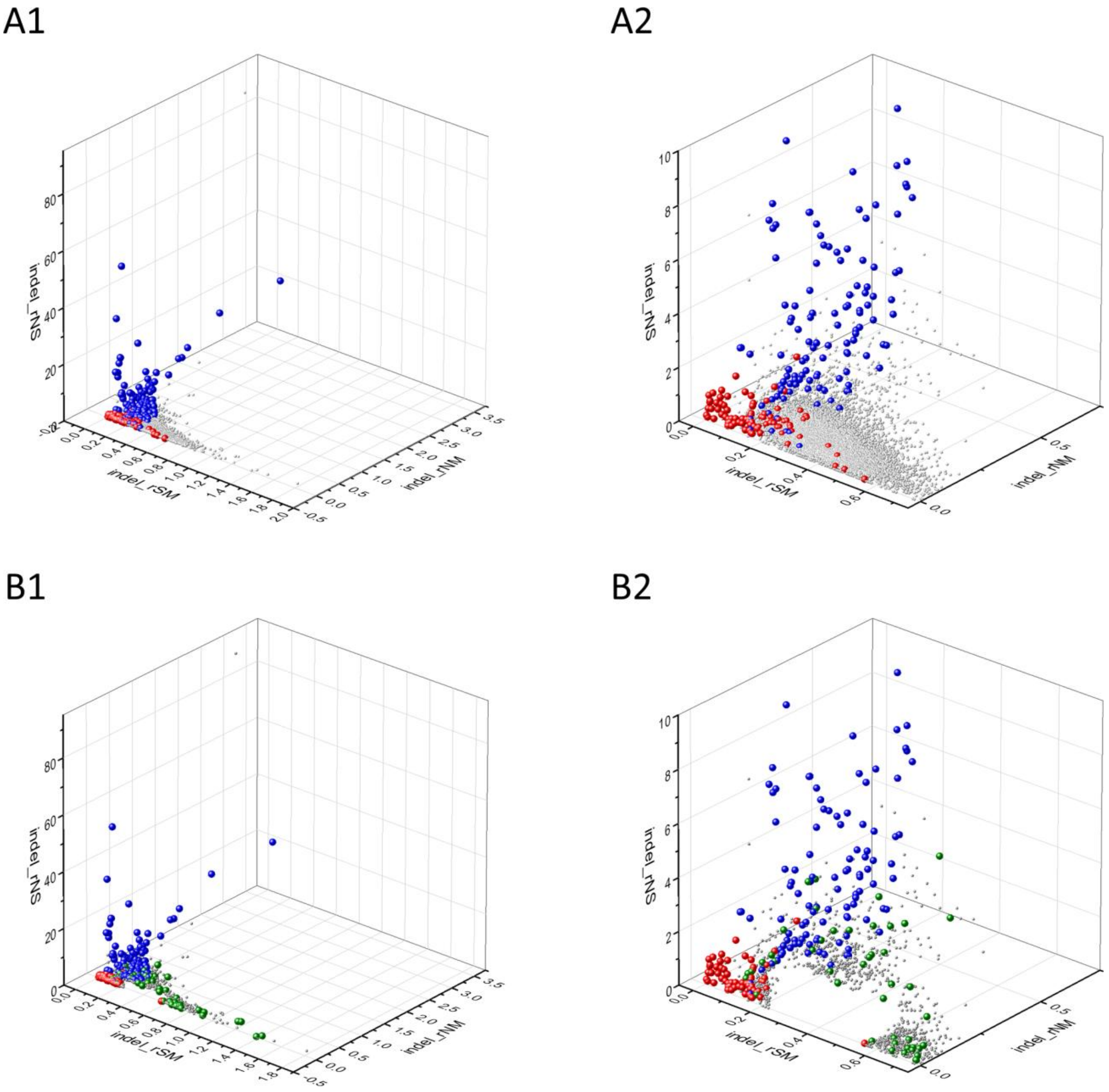


**Appendix 1 – figure 2. Analyses of indel_rSM, indel_rNM, indel_rNS parameters of human protein-coding genes of tumor tissues.** The figure shows the results of the analysis of 13930 transcripts containing at least 100 subtle, confirmed somatic mutations from tumor tissues, including only mutations identified as not SNPs. Axes *x*, *y* and *z* represent the indel_rSM, indel_rNM, indel_rNS values defined as the ratio of indel_fS/ indel_fM, indel_fN/ indel_fM, indel_fN/ indel_fS, respectively. Each ball represents a human transcript; the positions of transcripts of the genes identified by Vogelstein *et al*., (2013) as oncogenes (OGs, large red balls) or tumor suppressor genes (TSGs, large blue balls) are highlighted. **Panels A1, A2** show the distribution of the 13930 transcripts at different magnification. Note that the majority of human genes are present in a dense cluster but known OGs and TSGs separate significantly from the central cluster and from each other. The rNS and rNM values of TSGs are higher, whereas the rSM and rNM values of OGs are lower than those of passenger genes. **Panels B1,B2** show data only for candidate cancer genes present in the CG_SO$^{2SD}$_SSI$^{2SD}$ list (see **Supplementary file 6**). The positions of transcripts of the genes identified by Vogelstein *et al*., (2013) as oncogenes (OGs, large red balls) or tumor suppressor genes (TSGs, large blue balls) are highlighted. The positions of novel cancer gene transcripts validated in the present work are highlighted as large green balls.

The non-passenger gene set defined by 2SD cut-off value (see ***Appendix 1 – figure 2***. ***B1, B2***, **Supplementary file 3**) is hereafter referred to as CG_SSI$^{r2_2SD}$ for Cancer Gene_ Substitution and Subtle Indels deviating from mean indel_rSM, indel_rNM, indel_rNS values by more than 2SD (**Supplementary file 4**). This gene set has a total of 861 transcripts, containing 40 transcripts of OGs, 98 transcripts of TSGs genes, 86 transcripts of CGC genes and 637 transcripts (derived from 546 genes) not found in the OG, TSG and CGC cancer gene lists (**Supplementary file 3**).

The mean parameters of TSGs differ markedly from those of passenger genes in that rNS and rNM values are higher, reflecting the dominance of positive selection for inactivating mutations. The parameters for OGs on the other hand, differ from those of passenger genes in that indel_rSM values of OGs are significantly lower, reflecting positive selection for missense mutations (see ***Appendix 1 – figure 2, Panels A1, A2***). Interestingly, in this representation some oncogenes (e.g. *BCL2*) have unusally high scores of indel_rSM suggesting that in the case of these oncogenes purifying selection may override positive selection for amino acid changing mutations.

As mentioned above, the non-passenger gene set defined by a cut-off values of 2SD contains 637 transcripts (derived from 546 genes) not found in the OG, TSG or CGC lists. Since the majority of these genes have paramerters that assign them to the OG or TSG clusters, they can be regarded as candidate oncogenes or tumor suppressor genes. There is a group of genes that deviate from the clusters of passenger genes, OGs and TSGs (see ***Appendix 1 – figure 2, Panels B1, B2***) in that they have unusually high indel_rSM values. Since high indel_rSM values may be indicative of purifying selection we assume that they may correspond to tumor essential genes important for the growth and survival of tumors. The 546 putative cancer genes listed in CG_SO$^{indel_r2_2SD}$ of **Supplementary file 3**, were subjected to further analyses to decide whether they qualify as candidate OGs, TSGs, TEGs or the deviation of their mutation pattern from those of passenger genes is not the result of natural selection. For examples of these analyses see **Appendix 2**.

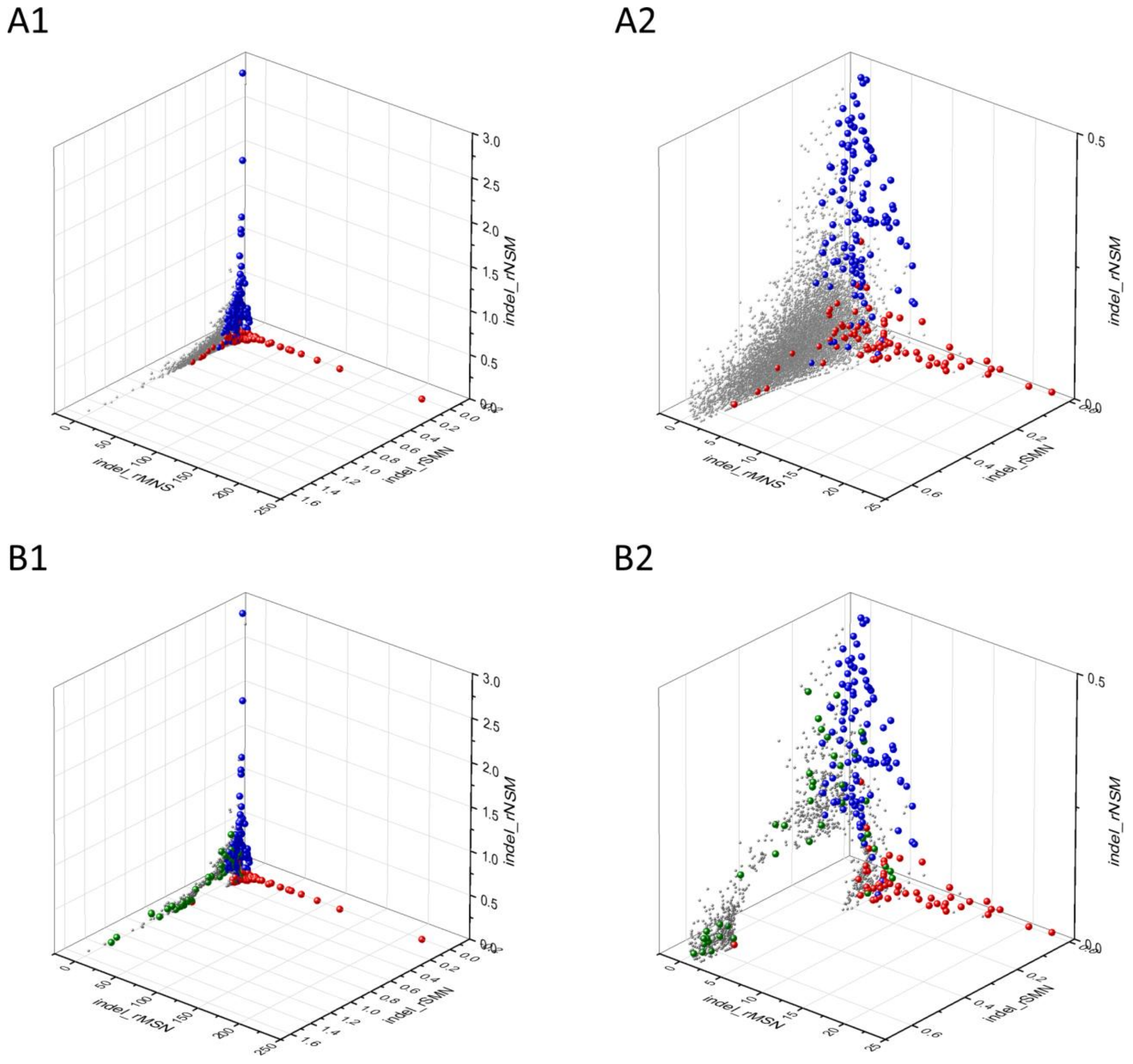


**Appendix 1 – figure 3. Analyses of indel_rSMN, indel_rMSN and indel_rNSM parameters of human protein-coding genes of tumor tissues.** The figure shows the results of the analysis of 13930 transcripts containing at least 100 subtle, confirmed somatic mutations from tumor tissues.. Axes *x*, *y* and *z* represent paramerters indel_rSMN, indel_rMSN and indel_rNSM defined as the ratio of indel_fS/(indel_fM+indel_fN), indel_fM/(indel_fS+indel_fN) and indel_fN/(indel_fS+indel_fM), respectively. Each ball represents a human transcript; the positions of transcripts of the genes defined by Vogelstein *et al*., (2013) as oncogenes (OGs, red balls) or tumor suppressor genes (TSGs, blue balls) are highlighted. **Panels A1** and **A2** show the distribution of the 13930 transcripts at different magnification. Note that the majority of human genes are present in a dense cluster but known OGs and TSGs separate significantly from the central cluster and from each other. The indel_rNSM values of TSGs are higher, their indel_rMSN and indel_rSMN are lower than those of passenger genes. OGs also separate from passenger genes in that their indel_rMSN values are higher and their indel_rSMN values are lower than those of passenger genes. **Panels B1,B2** show data at different magnification only for candidate cancer genes present in the CG_SO$^{2SD}$_SSI$^{2SD}$ list (see **Supplementary file 6**). The positions of transcripts of the genes identified by Vogelstein *et al*., (2013) as oncogenes (OGs, large red balls) or tumor suppressor genes (TSGs, large blue balls) are highlighted. The positions of novel cancer gene transcripts validated in the present work are highlighted as large green balls.

The separation of known cancer genes from the majority of human genes is also observed in 3D representations of parameters indel_rSMN, indel_rMSN and indel_rNSM defined as the ratio of indel_fS/(indel_fM+indel_fN), indel_fM/(indel_fS+indel_fN) and indel_fN/(indel_fS+indel_fM), respectively (see ***Appendix 1 – figure 3, Panels A1, A2***). In this representation the genes are present in a three pronged cluster.

The set of genes (4369 transcripts) with values that deviate from the mean by more than 1SD contained 78 OG transcripts, 132 TSG transcripts and 354 CGC gene transcripts (**Supplementary file 4)**. The central cluster of genes, deviating from mean rSMN, rMSN and rNSM values by ≤1SD is hereafter referred to as $PG\_SO^{indel\_r3\_1SD}$ (for Passenger Gene_ Substitution and Subtle Indels deviating from mean indel_rSMN, indel_rMSN and indel_rNSM values by ≤1SD).

The non-passenger gene set defined by 2SD cut-off value (see ***Appendix 1 – figure 3, Panels B1, B2,*** **Supplementary file 3**) is hereafter referred to as $CG\_SSI^{r3\_2SD}$ for Cancer Gene_ Substitution and Subtle Indels deviating from mean indel_rSMN, indel_rMSN and indel_rNSM values by more than 2SD (**Supplementary file 4**). This gene set has a total of 823 transcripts, containing transcripts of 37 OGs, 100 TSGs, 86 CGC genes and 600 transcripts (derived from 510 genes) not found in the OG, TSG and CGC cancer gene lists (**Supplementary file 3**).

The mean parameters of TSGs differ markedly from those of passenger genes in as much as indel_rNSM values of TSGs are higher and indel_rSMN values are lower, reflecting the dominance of positive selection for inactivating mutations. In the case of OGs on the other hand, indel_rMSN values are higher and indel_rNSM values are lower than those of passenger genes, reflecting positive selection for missense mutations and purifying selection avoiding nonsense mutations. Interestingly, some oncogenes have unusally high scores of indel_rSMN suggesting that in these cases (e.g. *BCL2*) purifying selection may override positive selection for amino acid changing mutations.

As mentioned above, the non-passenger gene set defined by a cut-off value of 2SD contains 600 transcripts (derived from 510 genes) not found in the OG, TSG or CGC lists. Since the majority of these genes have paramerters that assign them to the OG or TSG clusters, they can be regarded as candidate oncogenes or tumor suppressor genes.

In this representation we also note the existence of a group of genes that deviates from the clusters of passenger genes, OGS and TSGs (see ***Appendix 1 – figure 3***): their high indel_rSMN and low indel_rMSN and indel_rNSM values suggest that they experience purifying selection during tumor evolution, suggesting that they may be essential for the survival of tumors as oncogenes or tumor essential genes. The 510 putative cancer genes listed in $CG\_SSI^{r3\_2SD}$ of **Supplementary file 3**, were subjected to further analyses to decide whether they qualify as candidate oncogenes, tumor suppressor genes and tumor essential genes or the deviation of their mutation pattern from those of passenger genes is not the result of natural selection. For some typical examples of these analyses see **Appendix 2**.
