## Appendix 2 for "Use of signals of positive and negative selection to distinguish cancer genes and passenger genes"

### Examples of genes with strong sign atures of positive and/or negative selection

The assignments of the genes to key cellular processes of carcinogenesis are summarized in **Table 1** of the main text.

#### Novel cancer genes positively selected for truncating mutations

##### Beta-1,3-galactosyltransferase 1, encoded by the *B3GALT1* gene

B3GALT1 belongs to the glycosyltransferase 31 family. It transfers galactose from UDP-alpha-D-galactose to substrates with a terminal beta-N-acetylglucosamine residue. B3GALT1 is involved in the biosynthesis of the carbohydrate moieties of glycolipids and glycoproteins.

It has been suggested that loss of the activity of B3GALT1 may play an important role in aberrant protein glycosylation and tumor progression in colorectal cancers (Venkitachalam *et al*., 2016). Although such a role would be consistent with positive selection for inactivating mutations, analysis of the distribution of nonsense mutations along the protein sequence suggests that the high rNSM value is an artefact, rather than a signature of positive selection for iactivating mutations. The high rate of nonsense substitutions vs. sense substitutions is due to the fact that the majority of sequences contain nonsense substitution at a single site (p.R199*). Since there is no reason why selection would favor nonsense mutation at a single site it seems more likely that it reflects some sort of data deposition error. It is noteworthy in this respect that all the samples containing the p.R199* mutations originate from different regions of pancreatic tumor tissue samples from a single study (Yachida *et al*; 2016).

Venkitachalam S, Revoredo L, Varadan V, Fecteau RE, Ravi L, Lutterbaugh J, Markowitz SD, Willis JE, Gerken TA, Guda K. Biochemical and functional characterization of glycosylation-associated mutational landscapes in colon cancer. Sci Rep. 2016; 6:23642.

Yachida S, Wood LD, Suzuki M, Takai E, Totoki Y, Kato M, Luchini C, Arai Y, Nakamura H, Hama N, Elzawahry A, Hosoda F, Shirota T et al. Genomic Sequencing Identifies ELF3 as a Driver of Ampullary Carcinoma. Cancer Cell. 2016; 29:229-2240.

##### Bone morphogenetic protein receptor type-2, encoded by the *BMPR2* gene

Bone morphogenetic protein receptor type-2, a member of the TGF beta family of growth factor receptors. Upon ligand binding it forms a receptor complex consisting of two type II and two type I transmembrane serine/threonine kinases and activates SMAD transcriptional regulators.

There is convincing evidence in the literature that BMPR2 is a tumor suppressor. The *BMPR2* gene has been shown to contain several somatic frameshift mutations and to be inactivated in gastric and colorectal cancers with microsatellite instability (Kodach *et al*., 2008, Park *et al*., 2010). Loss of BMPR2 function has been found to result in increased tumorigenicity in human prostate cancer cells (Kim *et al*., 2004). More recent studies have shown that disruption of BMPR2 expression promotes mammary carcinoma metastases (Owens *et al*., 2012, Pickup *et al*., 2015). It was shown that loss of BMPR2 results in increased chemokine expression, which facilitates inflammation by a sustained increase in myeloid cells. The chemokines increased in *BMPR2* deleted cells correlated with poor outcome in human breast cancer patients, suggesting that BMPR2 has tumor suppressive functions in the stroma by regulating inflammation (Pickup *et al*., 2015).

Kim IY, Lee DH, Lee DK, Ahn HJ, Kim MM, Kim SJ, Morton RA. Loss of expression of bone morphogenetic protein receptor type II in human prostate cancer cells. Oncogene. 2004; 23:7651-7659.

Kodach LL, Wiercinska E, de Miranda NF, Bleuming SA, Musler AR, Peppelenbosch MP, Dekker E, van den Brink GR, van Noesel CJ, Morreau H, Hommes DW, Ten Dijke P, Offerhaus GJ, et al. The bone morphogenetic protein pathway is inactivated in the majority of sporadic colorectal cancers. Gastroenterology. 2008; 134:1332-1341

Owens P, Pickup MW, Novitskiy SV, Chytil A, Gorska AE, Aakre ME, West J, Moses HL. Disruption of bone morphogenetic protein receptor 2 (BMPR2) in mammary tumors promotes metastases through cell autonomous and paracrine mediators. Proc Natl Acad Sci U S A. 2012; 109:2814-2819

Park SW, Hur SY, Yoo NJ, Lee SH. Somatic frameshift mutations of bone morphogenic protein receptor 2 gene in gastric and colorectal cancers with microsatellite instability.
APMIS. 2010; 118:824-829.

Pickup MW, Hover LD, Polikowsky ER, Chytil A, Gorska AE, Novitskiy SV, Moses HL, Owens P. *BMPR2* loss in fibroblasts promotes mammary carcinoma metastasis via increased inflammation. Mol Oncol. 2015; 9:179-191

#### Bromodomain-containing protein 7, encoded by the *BRD*7 gene

BRD7 is a crucial component of both functional p53 and BRCA1 pathways and recent studies have fully established BRD7 as a tumor suppressor. The expression of BRD7 was shown to be downregulated in various cancers, including breast cancer, nasopharyngeal carcinoma, gastric cancer, colorectal carcinoma, ovarian cancer, lung adenocarcinoma, non-small cell lung cancer, hepatocellular carcinoma and prostate cancer. Moreover, BRD7 inhibited cancer cell growth and metastasis and promoted apoptosis *in vitro* and *in vivo* (Yu, Li and Shen, 2016, Gao, Wang and Gao, 2016, Chen *et al*., 2016, Li *et al*., 2015).

Recent studies suggest that BRD7 exerts it tumor suppressive role through multiple pathways, by suppressing cell proliferation, initiating cell apoptosis and reducing aerobic glycolysis (Niu *et al*., 2018). These studies suggest that BRD7 inhibits the Warburg effect through inactivation of the HIF1α/LDHA axis.

Chen CL, Wang Y, Pan QZ, Tang Y, Wang QJ, Pan K, Huang LX, He J, Zhao JJ, Jiang SS, Zhang XF, Zhang HX, Zhou ZQ et al.. Bromodomain-containing protein 7 (BRD7) as a potential tumor suppressor in hepatocellular carcinoma. Oncotarget. 2016; 7:16248-16261.

Gao Y, Wang B, Gao S. *BRD7* Acts as a Tumor Suppressor Gene in Lung Adenocarcinoma.
PLoS One. 2016; 11:e0156701

Li D, Yang Y, Zhu G, Liu X, Zhao M, Li X, Yang Q. MicroRNA-410 promotes cell proliferation by targeting *BRD7* in non-small cell lung cancer. FEBS Lett. 2015; 589:2218-2223

Niu W, Luo Y, Wang X, Zhou Y, Li H, Wang H, Fu Y, Liu S, Yin S, Li J, Zhao R, Liu Y, Fan S et al. BRD7 inhibits the Warburg effect and tumor progression through inactivation of HIF1α/LDHA axis in breast cancer. Cell Death Dis. 2018; 9:519.

Yu X, Li Z, Shen J. *BRD7*: a novel tumor suppressor gene in different cancers. Am J Transl Res. 2016; 8:742-748.

#### Inhibitor of growth protein 1 encoded by the *ING1* gene

*ING1* encodes a nuclear, cell cycle-regulated protein, overexpression of which efficiently blocks cell growth and is capable of inducing apoptosis in different experimental systems (Toyama *et al*., 1999). ING1 is known to cooperate with p53/TP53 in the negative regulatory pathway of cell growth by modulating p53-dependent transcriptional activation.

The tumor suppressor status of *ING1* has been fully established since several studies have described the loss of ING1 protein expression in human tumors and *ING1* knockout mice were reported to have spontaneously developed tumors, B cell lymphomas, and soft tissue sarcomas (Guérillon, Larrieu and Pedeux, 2013).

ING1 levels were found to be lower in breast tumors compared to adjacent normal breast tissue (Thakur *et al*. 2014). Decreasing levels of ING1 increased, and increasing levels decreased migration and invasion of cancer cells *in vitro*. ING1 overexpression also blocked cancer cell metastasis *in vivo* and eliminated tumor-induced mortality in mouse models.

ING1 can inhibit the growth of lung cancer cell lines through the induction of cell cycle arrest and apoptosis by forming a complex with p53 (Luo *et al*., 2011, Bose *et al*., 2014)

Genetic alterations that abrogate the normal function of *ING1* may contribute to esophageal squamous cell carcinogenesis (Chen *et al*., 2001). Mutations of the *ING1* tumor suppressor gene detected in human melanoma abrogate nucleotide excision repair activity of the protein (Campos *et al*., 2004). Nonsense mutations cluster in the region of residues 339-378. These mutations eliminate the Zn finger domain and polybasic region, which are involved in interaction with histone H3 trimethylated at Lys4 (H3K4me3). It is noteworthy thar histone H3K4me3 binding is required for the DNA repair and apoptotic activities of the ING1 tumor suppressor (Pena *et al*., 2008).

Bose P, Thakur SS, Brockton NT, Klimowicz AC, Kornaga E, Nakoneshny SC, Riabowol KT, Dort JC Tumor cell apoptosis mediated by cytoplasmic *ING1* is associated with improved survival in oral squamous cell carcinoma patients. Oncotarget. 2014; 5:3210-3219.

Campos EI, Martinka M, Mitchell DL, Dai DL, Li G. Mutations of the *ING1* tumor suppressor gene detected in human melanoma abrogate nucleotide excision repair. Int J Oncol. 2004; 25:73-80.

Chen L, Matsubara N, Yoshino T, Nagasaka T, Hoshizima N, Shirakawa Y, Naomoto Y, Isozaki H, Riabowol K, Tanaka N. Genetic alterations of candidate tumor suppressor *ING1* in human esophageal squamous cell cancer. Cancer Res. 2001; 61:4345-4349.

Guérillon C, Larrieu D, Pedeux R. *ING1* and *ING2*: multifaceted tumor suppressor genes. Cell Mol Life Sci. 2013; 70:3753-3772.

Luo ZG, Tang H, Li B, Zhu Z, Ni CR, Zhu MH. Genetic alterations of tumor suppressor *ING1* in human non-small cell lung cancer. Oncol Rep. 2011; 25:1073-1081

Peña PV1, Hom RA, Hung T, Lin H, Kuo AJ, Wong RP, Subach OM, Champagne KS, Zhao R, Verkhusha VV, Li G, Gozani O, Kutateladze TG. Histone H3K4me3 binding is required for the DNA repair and apoptotic activities of ING1 tumor suppressor.
J Mol Biol. 2008 Jul 4;380(2):303-12.

Thakur S, Singla AK, Chen J, Tran U, Yang Y, Salazar C, Magliocco A, Klimowicz A, Jirik F, Riabowol K. Reduced ING1 levels in breast cancer promotes metastasis. Oncotarget. 2014; 5:4244-4256.

Toyama T, Iwase H, Watson P, Muzik H, Saettler E, Magliocco A, DiFrancesco L, Forsyth P, Garkavtsev I, Kobayashi S, Riabowol K. Suppression of *ING1* expression in sporadic breast cancer. Oncogene. 1999; 18:5187-5193.

#### MAX gene-associated protein, encoded by the *MGA* gene

MGA functions as a dual-specificity transcription factor, regulating the expression of both MAX-network and T-box family target genes. Suppresses transcriptional activation by MYC and inhibits MYC-dependent cell transformation. Recurrent inactivation of *MGA*, a suppressor of MYC, has been shown to occur in lymphocytic leukemia, and in both non-small cell lung cancer and small cell lung cancer, colorectal cancer (De Paoli *et al*., 2013, Romero *et al*., 2014, Jo *et al*., 2016).

De Paoli L, Cerri M, Monti S, Rasi S, Spina V, Bruscaggin A, Greco M, Ciardullo C, Famà R, Cresta S, Maffei R, Ladetto M, Martini M, et al. MGA, a suppressor of MYC, is recurrently inactivated in high risk chronic lymphocytic leukemia. Leuk Lymphoma. 2013; 54:1087-1090.

Jo YS, Kim MS, Yoo NJ, Lee SH. Somatic mutation of a candidate tumour suppressor *MGA* gene and its mutational heterogeneity in colorectal cancers. Pathology. 2016; 48:525-527.

Romero OA, Torres-Diz M, Pros E, Savola S, Gomez A, Moran S, Saez C, Iwakawa R, Villanueva A, Montuenga LM, Kohno T, Yokota J, Sanchez-Cespedes M. MAX inactivation in small cell lung cancer disrupts MYC-SWI/SNF programs and is synthetic lethal with BRG1. Cancer Discov. 2014; 4:292-303.

#### Proline-rich transmembrane protein 2, encoded by the *PRRT2* gene

PPRT2, as a component of the outer core of AMPAR complex, is involved in ion channel functions. *PRRT2* has been shown to be significantly downregulated in glioblastoma tissues compared with normal brain tissue (Bi *et al*., 2017, Li *et al*., 2018). Overexpression of PRRT2 strongly impaired the cell viability and promoted cell apoptosis. These anti-tumor effects indicate that PRRT2 acts as a tumor suppressor in glioma. PRRT2 has been shown to have an inhibitory effect on proliferation, consistent with the low expression level of PRRT2 in cancer versus normal samples (Alves *et al*., 2017).

Alves IT, Cano D, Böttcher R, van der Korput H, Dinjens W, Jenster G, Trapman J. A mononucleotide repeat in PRRT2 is an important, frequent target of mismatch repair deficiency in cancer. Oncotarget. 2017; 8:6043-6056

Bi G, Yan J, Sun S, Qu X. PRRT2 inhibits the proliferation of glioma cells by modulating unfolded protein response pathway. Biochem Biophys Res Commun. 2017; 485:454-460.

Li Z, Guo J, Ma Y, Zhang L, Lin Z. Oncogenic Role of MicroRNA-30b-5p in Glioblastoma Through Targeting Proline-Rich Transmembrane Protein 2. Oncol Res. 2018; 26:219-230

#### Ras GTPase-activating protein 1, encoded by the *RASA1* gene

RASA1 is an inhibitory regulator of the Ras-cyclic AMP pathway. Consistent with the tumor suppressor role of RASA1, the circular RNA circ-ITCH was shown to suppress ovarian carcinoma progression through targeting miR-145/*RASA1* signaling, by increasing the level of RASA1 (Hu *et al*., 2018).

There is evidence that *RASA1* is a potent tumor suppressor gene that is frequently downregulated or inactivated in several human cancer types. RASA1 expression is frequently reduced in breast cancer tissues, and the reduced RASA1 expression is associated with breast cancer progression and poor survival and disease-free survival of patients (Liu *et al*., 2015).

In hepatocellular carcinoma patients low level of RASA1 expression correlated with a significantly poorer survival compared to those with high level of RASA1 expression, suggesting that RASA1 could serve as an independent prognostic marker for hepatocellular carcinoma patients (Chen *et al*., 2017).

Analyses of melanoma whole genome sequencing data have led to the identification of two novel, clustered somatic missense mutations (Y472H and L481F) in RASA1 (Sung *et al*., 2016). Unlike wild type RASA1, these mutants, do not suppresses soft agar colony formation and tumor growth of melanoma cell lines. In addition to mutations, loss of RASA1 expression was frequently observed in metastatic melanoma samples and a low level of RASA1 mRNA expression was associated with decreased overall survival in melanoma patients. Thus, these data support that RASA1 is inactivated by mutations or by suppressed expression in melanoma and that RASA1 plays a tumor suppressive role.

The tumor suppressor role of RASA1 is also supported by the fact that knockdown or miR targeting of *RASA1* significantly enhanced invasion and migration of multiple pancreatic cancer cells (Sun *et al*., 2013, Kent, Mendell and Rottapel, 2016).

Chen YL, Huang WC, Yao HL, Chen PM, Lin PY, Feng FY, Chu PY. Down-regulation of RASA1 Is Associated with Poor Prognosis in Human Hepatocellular Carcinoma. Anticancer Res. 2017; 37:781-785.

Hu J, Wang L, Chen J, Gao H, Zhao W, Huang Y, Jiang T, Zhou J, Chen Y. The circular RNA circ-ITCH suppresses ovarian carcinoma progression through targeting miR-145/RASA1 signaling. Biochem Biophys Res Commun. 2018; 505:222-228.

Kent OA, Mendell JT, Rottapel R. Transcriptional Regulation of miR-31 by Oncogenic KRAS Mediates Metastatic Phenotypes by Repressing RASA1. Mol Cancer Res. 2016; 14:267-277.

Liu Y, Liu T, Sun Q, Niu M, Jiang Y, Pang D. Downregulation of Ras GTPase‑ activating protein 1 is associated with poor survival of breast invasive ductal carcinoma patients. Oncol Rep. 2015; 33:119-124.

Sun D, Wang C, Long S, Ma Y, Guo Y, Huang Z, Chen X, Zhang C, Chen J, Zhang J C/EB P-β-activated microRNA-223 promotes tumour growth through targeting RASA1 in human colorectal cancer. Br J Cancer. 2015; 112:1491-500.

Sung H, Kanchi KL, Wang X, Hill KS, Messina JL, Lee JH, Kim Y, Dees ND, Ding L, Teer JK, Yang S, Sarnaik AA, Sondak VK, et al. Inactivation of RASA1 promotes melanoma tumorigenesis via R-Ras activation. Oncotarget. 2016; 7:23885-23896.

#### E3 ubiquitin-protein ligase RNF128, encoded by the *RNF128* gene

E3 ubiquitin-protein ligase RNF128 catalyzes 'Lys-48'- and 'Lys-63'-linked polyubiquitin chains formation. Consistent with its suggested tumor suppressor role, downregulation of *RNF128* was found to predict poor prognosis in patients with urothelial carcinoma and urinary bladder. Downregulation of *RNF128* was correlated with cancer invasiveness and metastasis as well as reduced survival in patients (Lee *et al*., 2016). *RNF128* downregulation was also shown to correlate with the malignant phenotype of melanoma (Wei *et al*., 2019).

Lee YY, Wang CT, Huang SK, Wu WJ, Huang CN, Li CC, Chan TC, Liang PI, Hsing CH, Li CF. Downregulation of *RNF128* Predicts Progression and Poor Prognosis in Patients with Urothelial Carcinoma of the Upper Tract and Urinary Bladder. J Cancer. 2016; 7:2187-2196.

Wei CY, Zhu MX, Yang YW, Zhang PF, Yang X, Peng R, Gao C, Lu JC, Wang L, Deng XY, Lu NH1, Qi FZ, Gu JY. Downregulation of *RNF128* activates Wnt/β-catenin signaling to induce cellular EMT and stemness via CD44 and CTTN ubiquitination in melanoma. J Hematol Oncol. 2019; 12:21.

#### Monocarboxylate transporter 1, MCT1 encoded by the *SLC16A1* gene

SLC16A1 is a multipass plasma membrane protein that functions as a proton-coupled monocarboxylate transporter. It catalyzes the rapid transport across the plasma membrane of many monocarboxylates such as lactate. Depending on the tissue and on cicumstances, mediates the import or export of lactic acid. Deficiency of this lactate transporter may result in an acidic intracellular environment created by muscle activity with consequent degeneration of muscle.

Although the high values of rNSM would suggest a tumor suppressor role for *SLC16A1*, several studies suggest that the protein may serve a pro-oncogenic role. For example, depletion of *SLC16A1* was found to decrease cellular proliferation and invasion in both neuroblastoma and malignant cutaneous melanoma cell lines, suggesting its role as an oncogene (Avitabile *et al*., 2019). The pro-oncogenic role of MCT1 is also supported by the results of studies on esophageal squamous cell carcinoma ESCC. Kaplan-Meier survival analysis of ESCC patients in a high-MCT1 group had a lower overall survival and lower progression-free survival, whereas downregulation of MCT1 suppressed proliferation and survival of ESCC cells *in vitro* (Chen *et al*., 2019). Disrupting MCT1 function leads to an accumulation of intracellular lactate that rapidly disables tumor cell growth (Doherty *et al*., 2014).

MCT1 expression is elevated in glycolytic breast tumors, and high MCT1 expression predicts poor prognosis in breast and lung cancer patients. Similarly, the observations that MCT1 inhibition impairs proliferation of glycolytic breast cancer cells co-expressing MCT1 and MCT4 and that MCT1 loss-of-function decreases breast cancer cell proliferation and blocks growth of mammary fat pad xenograft tumors suggest a pro-oncogenic or tumor essential role for MCT1 (Hong *et al*., 2016).

A recent study, however, has led to the conclusion that MCT1 and MCT4 have opposing roles in carcinogenesis (Sukeda *et al*., 2019). In a retrospective survey conducted on patients who underwent surgical resection for pancreatic ductal adenocarcinoma the expression of MCT1, MCT4, and GLUT1 was assessed in tumor cells and cancer-associated fibroblasts (CAFs) and the impact of their expression on patient outcome was also analyzed. In tumor cells, MCT1 expression was associated with extended overall and progression-free survival and decreased nodal metastasis. Conversely, MCT4 expression in CAFs was associated with shortened survival. In other words, in tumor cells, MCT1 expression is associated with better prognosis and reduced nodal metastasis in pancreatic cancer, contrary to findings of previous studies.

It is noteworthy in this respcect that based on the pattern of mutations SLC16A1/MCT1 appers to be a tumor suppressor rather than a tumor essential gene in as much as it has a high proportion of truncating mutations.. It seems possible that glycolytic tumor cells that must get rid of lactate are selected for increased efflux and decreased influx of lactate and this might be achieved by increased expression of MCT4 and decreased activity of MCT1.

Avitabile M, Succoio M, Testori A, Cardinale A, Vaksman Z, Lasorsa VA, Cantalupo S, Esposito M, Cimmino F, Montella A, Formicola D, Koster J, Andreotti V, et al. Neural crest-derived tumor neuroblastoma and melanoma share 1p13.2 as susceptibility locus that shows a long-range interaction with the SLC16A1 gene. Carcinogenesis. 2019. pii: bgz153.

Chen X, Chen X, Liu F, Yuan Q, Zhang K, Zhou W, Guan S, Wang Y, Mi S, Cheng Y. Monocarboxylate transporter 1 is an independent prognostic factor in esophageal squamous cell carcinoma. Oncol Rep. 2019; 41:2529-2539.

Doherty JR, Yang C, Scott KE, Cameron MD, Fallahi M, Li W, Hall MA, Amelio AL, Mishra JK, Li F, Tortosa M, Genau HM, Rounbehler RJ, et al. Blocking lactate export by inhibiting the Myc target MCT1 Disables glycolysis and glutathione synthesis. Cancer Res. 2014;74:908-920.

Hong CS, Graham NA, Gu W, Espindola Camacho C, Mah V, Maresh EL, Alavi M, Bagryanova L, Krotee PAL, Gardner BK, Behbahan IS, Horvath S, Chia D, et al. MCT1 Modulates Cancer Cell Pyruvate Export and Growth of Tumors that Co-express MCT1 and MCT4. Cell Rep. 2016; 14:1590-1601.

Sukeda A, Nakamura Y, Nishida Y, Kojima M, Gotohda N, Akimoto T, Ochiai A. Expression of Monocarboxylate Transporter 1 Is Associated With Better Prognosis and Reduced Nodal Metastasis in Pancreatic Ductal Adenocarcinoma. Pancreas. 2019; 48:1102-1110.

#### Sprouty-related, EVH1 domain-containing protein 1, encoded by the *SPRED1* gene

The *SPRED1* gene, which encodes a negative regulator of mitogen-activated protein kinase (MAPK) signaling, has been shown to function as a tumor suppressor gene in several types of cancer (Pasmant *et al*., 2015, Ablain *et al*., 2018, Sun *et al*., 2019).

Ablain J, Xu M, Rothschild H, Jordan RC, Mito JK, Daniels BH, Bell CF, Joseph NM, Wu H, Bastian BC, Zon LI, Yeh I. Human tumor genomics and zebrafish modeling identify *SPRED1* loss as a driver of mucosal melanoma. Science. 2018;362:1055-1060

Sun J, Zhang J, Wang Y, Li Y, Zhang R. A Pilot Study of Aberrant CpG Island Hypermethylation of *SPRED1* in Acute Myeloloid Leukemia. Int J Med Sci. 2019; 16:324-330

Pasmant E, Gilbert-Dussardier B, Petit A, de Laval B, Luscan A, Gruber A, Lapillonne H, Deswarte C, Goussard P, Laurendeau I, Uzan B, Pflumio F, Brizard F, et al. SPRED1, a RAS MAPK pathway inhibitor that causes Legius syndrome, is a tumour suppressor downregulated in paediatric acute myeloblastic leukaemia. Oncogene. 2015; 34:631-638.

#### Homeobox protein TGIF1, encoded by the *TGIF1* gene

TGIF binds to a retinoid X receptor (RXR) responsive element from the cellular retinol-binding protein II promoter (CRBPII-RXRE). Inhibits the 9-cis-retinoic acid-dependent RXR alpha transcription activation of the retinoic acid responsive element. Active transcriptional corepressor of SMAD2.

There is evidence that TGIF1 may function as a tumor suppressor. In pancreatic ductal adenocarcinoma genetic inactivation of *TGIF1* in the context of oncogenic KRASG12D, culminated in the development of highly aggressive and metastatic pancreatic ductal adenocarcinoma (Parajuli *et al*., 2019, Weng *et al*., 2019). These authors have found that TGIF1 associates with TWIST1 and inhibits TWIST1 expression and activity, and this function is suppressed in the vast majority of human pancreatic ductal adenocarcinoma by KRASG12D /MAPK-mediated TGIF1 phosphorylation. Ablation of TWIST1 in KRASG12D;TGIF1KO mice blocked pancreatic ductal adenocarcinoma formation, providing evidence that TGIF1 restrains KRASG12D -driven pancreatic ductal adenocarcinoma through its ability to antagonize TWIST1.

The majority of available evidence, however, suggests that the protein plays a cancer promoting role. TGIF1 has been shown to promote the growth and migration of cancer cells in nonsmall cell lung cancer (Xiang *et al*., 2015). The authors have shown that expression of TGIF1 is elevated in NSCLC tissues, that TGIF1 promoted the growth and migration of cancer cells and that knocking down the expression of *TGIF1* inhibited the growth and migration of NSCLC cells. These studies have also shown that TGIF1 exerted its oncogenic role through beta-catenin/TCF signaling.

Studies on triple negative breast cancer have revealed that high levels of TGIF expression correlate with poor prognosis since TGIF promotes Wnt-driven mammary tumorigenesis. As to the molecular mechanism of the oncogenic role of TGIF: it has been shown that TGIF interacts with and sequesters Axin1 and Axin2 into the nucleus, disassembles the β-catenin-destruction complex leading to the accumulation of β-catenin that activates expression of Wnt target genes (Zhang *et al*., 2015, Razzaque and Atfi, 2016).

In harmony with an oncogenic role of TGIF in breast cancer, silencing of *TGIF* was found to suppress the migration, invasion and metastasis of the human breast cancer cells in both *in vitro* and *in vivo* experiments (Wang *et al*., 2018).

*TGIF1* has also been found to be significantly upregulated in some colorectal cancers and to promote adenoma growth in the context of mutant Apc (Shah *et al*., 2019). Overexpression of TGIF1 markedly promoted the proliferation of colorectal cancer cells through the activation of Wnt/β-catenin signaling (Wang *et al*., 2017).

In summary, the majority of data suggest that *TGIF1* may act as an oncogene, despite the fact that the high proportion of truncating indel mutations would indicate a tumor suppressor function. Since the transcription regulator TGIF1 may play both pro-oncogenic and tumor suppressor functions (in different cellular processes) our observation that during tumor evolution selection for truncating mutations appears to dominate for TGIF1 suggests that the selection pressure to eliminate the tumor suppressor activity may override the pressure to preserve it oncogenic activities.

Parajuli P, Singh P, Wang Z, Li L, Eragamreddi S, Ozkan S, Ferrigno O, Prunier C, Razzaque MS, Xu K, Atfi A. TGIF1 functions as a tumor suppressor in pancreatic ductal adenocarcinoma. EMBO J. 2019; 38:e101067.

Razzaque MS, Atfi A. TGIF function in oncogenic Wnt signaling. Biochim Biophys Acta. 2016; 1865:101-104.

Shah A, Melhuish TA, Fox TE, Frierson HF Jr, Wotton D. TGIF transcription factors repress acetyl CoA metabolic gene expression and promote intestinal tumor growth. Genes Dev. 2019; 33:388-402.

Wang JL, Qi Z, Li YH, Zhao HM, Chen YG, Fu W. TGFβ induced factor homeobox 1 promotes colorectal cancer development through activating Wnt/β-catenin signaling. Oncotarget. 2017; 8:70214-70225.

Wang Y, Li L, Wang H, Li J, Yang H. Silencing *TGIF* suppresses migration, invasion and metastasis of MDA‑ MB‑ 231 human breast cancer cells. Oncol Rep. 2018; 39:802-808

Weng CC, Hsieh MJ, Wu CC, Lin YC, Shan YS, Hung WC, Chen LT, Cheng KH. Loss of the transcriptional repressor TGIF1 results in enhanced Kras-driven development of pancreatic cancer. Mol Cancer. 2019; 18:96.

Xiang G, Yi Y, Weiwei H, Weiming W. TGIF1 promoted the growth and migration of cancer cells in nonsmall cell lung cancer. Tumour Biol. 2015; 36:9303-9310

Zhang MZ, Ferrigno O, Wang Z, Ohnishi M, Prunier C, Levy L, Razzaque M, Horne WC, Romero D, Tzivion G, Colland F, Baron R, Atfi A. TGIF governs a feed-forward network that empowers Wnt signaling to drive mammary tumorigenesis. Cancer Cell. 2015; 27:547-560

#### Trinucleotide repeat-containing gene 6B protein, encoded by the *TNRC6B* gene

*TNRC6B* is a key miRNA-processing gene that plays a role in RNA-mediated gene silencing by both micro-RNAs (miRNAs) and short interfering RNAs (siRNAs). TNRC6B is required for miRNA-dependent translational repression and siRNA-dependent endonucleolytic cleavage of complementary mRNAs by argonaute family proteins.

Genomic analysis of liver cancer have identified *TNRC6B* as a significantly mutated gene, suggesting that it may be an important driver gene (Li *et al*., 2018). Consistent with its putative tumor suppressor role, DNA methylation of *TNRC6B* has been suggested to play a role in early carcinogenesis (Joyce *et al.*, 2018).

Joyce BT, Zheng Y, Zhang Z, Liu L, Kocherginsky M, Murphy R, Achenbach CJ, Musa J, Wehbe F, Just A, Shen J, Vokonas P, Schwartz J, et al. miRNA-Processing Gene Methylation and Cancer Risk. Cancer Epidemiol Biomarkers Prev. 2018; 27:550-557.

Li X, Xu W, Kang W, Wong SH, Wang M, Zhou Y, Fang X, Zhang X, Yang H, Wong CH, To KF, Chan SL, Chan MTV, et al. Genomic analysis of liver cancer unveils novel driver genes and distinct prognostic features. Theranostics. 2018; 8:1740-1751.

#### Dual specificity protein kinase TTK, encoded by the *TTK* gene

TTK, capable of phosphorylating serine, threonine, and tyrosine residues of proteins, plays a role in cell proliferation. Although, intuitively the high rate of truncating mutations would suggest a tumor suppressor role for TTK, all the available evidence indicates that it acts as an oncogene.

It has been shown that dual specificity kinase TTK is strongly overexpressed in human pancreatic ductal adenocarcinoma, suggesting a cancer promoting role. In harmony with such a role, following *TTK* knockdown cell proliferation was significantly attenuated whereas apoptosis and necrosis rates were significantly increased. Apoptosis was associated with increased formation of micronuclei, suggesting that loss of TTK results in chromosomal instability and mitotic catastrophe (Kaistha *et al*., 2014).

Levels of TTK protein were also found to be significantly elevated in neoplastic tissues of liver cancer patients, when compared with adjacent hepatic tissues. In an experimental animal model it was shown that *in vitro* knockdown of *TTK* effectively blocks intrahepatic growth of human hepatic carcinoma cell xenografts, suggesting that targeted TTK inhibition might have clinical utility in the therapy of liver cancer (Miao *et al*., 2016).

In a recent study dual specificity protein kinase TTK has been identified as the most up-regulated and differentially expressed kinase in glioma stem-like cells that are responsible for tumorigenesis and subsequent tumor recurrence in glioblastoma. TTK expression was highly enriched in glioblastoma and was inversely correlated with a poor prognosis (Wang *et al.*, 2017).

The deubiquitinase, USP9X has been implicated in multiple cancers and its oncogenic effects were shown to be exerted at least in part through dual specificity protein kinase TTK (Chen *et al*., 2018). USP9X was found to stabilize TTK by efficient deubiquitination of the kinase; levels of USP9X and TTK were significantly elevated and positively correlated in tumor tissues, suggesting that the USP9X-TTK axis plays a critical role in carcinogenesis. In harmony with the synergism of these oncogenes, knockdown of *USP9X* or *TTK* inhibited cell proliferation, migration and tumorigenesis.

The explanation for the apparent contradiction of the oncogenic role of TTK and the abundance of truncating mutations in the protein probably lies in the fact that – unlike in the case of typical tumor suppressor genes – mutations are not randomly distributed along the protein sequence. The truncating mutations are practically restricted to the very C-terminal end of the protein (EKKRGKK, residues 851-857), downstream of the catalytic domain and missense mutations also cluster in this C-terminal end. It seems likely that this region is involved in some negative control of the activity of TTK and missense and truncating mutations liberate TTK from this negative control. It is unclear at present whether the mutations affecting this C-terminal motif activate the TTK proto-oncogene by interfering with its ubiquitination or by affecting its subcellular localization.

Chen X, Yu C, Gao J, Zhu H, Cui B, Zhang T, Zhou Y, Liu Q, He H, Xiao R, Huang R, Xie H, Gao D, Zhou H. A novel USP9X substrate TTK contributes to tumorigenesis in non-small-cell lung cancer. Theranostics. 2018; 8:2348-2360.

Kaistha BP, Honstein T, Müller V, Bielak S, Sauer M, Kreider R, Fassan M, Scarpa A, Schmees C, Volkmer H, Gress TM, Buchholz M. Key role of dual specificity kinase TTK in proliferation and survival of pancreatic cancer cells. Br J Cancer. 2014; 111:1780-1787.

Miao R, Wu Y, Zhang H, Zhou H, Sun X, Csizmadia E, He L, Zhao Y, Jiang C, Miksad RA, Ghaziani T, Robson SC, Zhao H. Utility of the dual-specificity protein kinase TTK as a therapeutic target for intrahepatic spread of liver cancer. Sci Rep. 2016; 6:33121.

Wang J, Xie Y, Bai X, Wang N, Yu H, Deng Z, Lian M, Yu S, Liu H, Xie W, Wang M. Targeting dual specificity protein kinase TTK attenuates tumorigenesis of glioblastoma. Oncotarget. 2017; 9:3081-3088.

#### Zinc finger CCCH domain-containing protein 13, encoded by the *ZC3H13* gene

ZC3H13 is associated with a complex that mediates N6-methyladenosine (m6A) methylation of RNAs, a modification that plays a role in the efficiency of mRNA splicing and RNA processing. It acts s as a key regulator of m6A methylation by promoting m6A methylation of mRNAs at the 3'-UTR. ZC3H13 has been shown to serve as a tumor suppressor in colorectal cancer (Zhu *et al*., 2019).

Zhu D, Zhou J, Zhao J, Jiang G, Zhang X, Zhang Y, Dong M. ZC3H13 suppresses colorectal cancer proliferation and invasion via inactivating Ras-ERK signaling. J Cell Physiol. 2019; 234:8899-8907

#### mRNA decay activator protein ZFP36L2, encoded by the *ZFP36L2* gene

*ZFP36L2* has been selected as a gene characterized by very high values of indel_rNSM, suggesting positive selection for truncating mutations. Although the closely related *ZFP36L1* gene is not present in the lists defined by the CG_SO and CG_SSI lists defined by the 2SD cut-off values, it is also characterized by very high values of rNSM (**Supplementary file 3**).

ZFP36L1 and ZFP36L2 zinc-finger RNA-binding proteins destabilize several cytoplasmic AU-rich element (ARE)-containing mRNA transcripts by promoting their poly(A) tail removal or deadenylation, and hence provide a mechanism for attenuating protein synthesis. The proteins are necessary for thymocyte development and prevention of T-cell acute lymphoblastic leukemia transformation by promoting ARE-mediated mRNA decay of the mRNA of oncogenic factors.

Deletion of the genes *ZFP36L1* and *ZFP36L2* leads to perturbed thymic development and T lymphoblastic leukemia (Hodson *et al*., 2010).

ZFP36L1 and ZFP36L2 play a negative role in cell proliferation. Forced expression of ZFP36L1 or ZFP36L2 inhibited cell proliferation in colorectal cancer cell lines, whereas knockdown of these genes increased cell proliferation (Suk *et al*., 2018). ZFP36L2 has been validated as an important tumour-suppressor specific to oesophageal squamous cell carcinomas (Lin *et al*., 2018).

Hodson DJ, Janas ML, Galloway A, Bell SE, Andrews S, Li CM, Pannell R, Siebel CW, MacDonald HR, De Keersmaecker K, Ferrando AA, Grutz G, Turner M. Deletion of the RNA-binding proteins ZFP36L1 and ZFP36L2 leads to perturbed thymic development and T lymphoblastic leukemia. Nat Immunol. 2010; 11:717-724.

Lin DC, Dinh HQ, Xie JJ, Mayakonda A, Silva TC, Jiang YY, Ding LW, He JZ, Xu XE, Hao JJ, Wang MR, Li C, Xu LY et al. Identification of distinct mutational patterns and new driver genes in oesophageal squamous cell carcinomas and adenocarcinomas. Gut. 2018; 67:1769-1779.

Suk FM, Chang CC, Lin RJ, Lin SY, Liu SC, Jau CF, Liang YC. ZFP36L1 and ZFP36L2 inhibit cell proliferation in a cyclin D-dependent and p53-independent manner. Sci Rep. 2018; 8:2742

#### Zinc finger protein 276, encoded by the *ZNF276* gene

Zinc finger protein is involved in transcriptional regulation.

It has been suggested that ZNF276 may be a tumor suppressor in breast cancer progression in colorectal cancers (Wong *et al*., 2016).

Although such a role would be consistent with positive selection for inactivating mutations, analysis of the distribution of nonsense mutations along the protein sequence suggests that the high rNSM value is an artefact, rather than a signature of positive selection for inactivating mutations. The high rate of nonsense substitutions vs. sense substitutions is due to the fact that the majority of sequences contain nonsense substitution at a single site (p.Q217*). Since there is no reason why selection would favor nonsense mutation at a single site it seems more likely that it reflects some sort of data deposition error. It is noteworthy in this respect that all the samples containing the p.Q217* mutations originate from different regions of pancreatic tumor tissue samples from a single study (Yachida *et al*; 2016).


Wong JC, Gokgoz N, Alon N, Andrulis IL, Buchwald M. Cloning and mutation analysis of ZFP276 as a candidate tumor suppressor in breast cancer. J Hum Genet. 2003; 48:668-671.

Yachida S, Wood LD, Suzuki M, Takai E, Totoki Y, Kato M, Luchini C, Arai Y, Nakamura H, Hama N, Elzawahry A, Hosoda F, Shirota T et al. Genomic Sequencing Identifies ELF3 as a Driver of Ampullary Carcinoma. Cancer Cell. 2016; 29:229-240.


#### Zinc finger protein 750, encoded by the *ZNF750* Gene

Zinc finger protein 750 is a transcription factor required for terminal epidermal differentiation, it acts downstream of p63/TP63. Its mutations have been shown to abolish the ability to induce epidermal terminal differentiation. In harmony with its mutation pattern, numerous studies suggest a tumor suppressor role for ZNF750.

Analysis of cancer genes across 21 tumor types identified *ZNF750* as a gene harboring many early frameshift and nonsense mutations in head and neck cancer and as the only known gene residing in a small current focal deletion in head and neck and lung squamous cancers (Lawrence *et al*., 2014). *ZNF750* has also been identified as a tumor suppressor in oral and esophageal squamous cell carcinoma (Yang *et al*., 2017, Nambara *et al*., 2017, Hazawa *et al*., 2017, Otsuka *et al*., 2018). Studies on the clonal evolution in esophageal squamous cell carcinoma revealed that the majority of driver mutations in this cancer occurred in the tumor-suppressor genes, including *TP53, KMT2D* and *ZNF750* (Hao *et al*., 2016).


Hao JJ, Lin DC, Dinh HQ, Mayakonda A, Jiang YY, Chang C, Jiang Y, Lu CC, Shi ZZ, Xu X, Zhang Y, Cai Y, Wang JW, et al. Spatial intratumoral heterogeneity and temporal clonal evolution in esophageal squamous cell carcinoma. Nat Genet. 2016; 48:1500-1507

Hazawa M, Lin DC, Handral H, Xu L, Chen Y, Jiang YY, Mayakonda A, Ding LW, Meng X, Sharma A, Samuel S, Movahednia MM, Wong RW et al. ZNF750 is a lineage-specific tumour suppressor in squamous cell carcinoma. Oncogene. 2017; 36:2243-2254.

Lawrence MS, Stojanov P, Mermel CH, Garraway LA,. Golub TR, Meyerson M, Gabriel SB, Lander ES, Getz G. Discovery and saturation analysis of cancer genes across 21 tumor types
Nature. 2014; 505: 495–501.

Nambara S, Masuda T, Tobo T, Kidogami S, Komatsu H, Sugimachi K, Saeki H, Oki E, Maehara Y, Mimori K. Clinical significance of *ZNF750* gene expression, a novel tumor suppressor gene, in esophageal squamous cell carcinoma. Oncol Lett. 2017; 14:1795-1801.

Otsuka R, Akutsu Y, Sakata H, Hanari N, Murakami K, Kano M, Toyozumi T, Takahashi M, Matsumoto Y, Sekino N, Yokoyama M, Okada K, Shiraishi T, et al. ZNF750 Expression Is a Potential Prognostic Biomarker in Esophageal Squamous Cell Carcinoma. Oncology. 2018; 94:142-148

Yang H, Pan L, Xu C, Zhang Y, Li K, Chen S, Zhang B, Liu Z, Wang LX, Chen H. Overexpression of tumor suppressor gene *ZNF750* inhibits oral squamous cell carcinoma metastasis. Oncol Lett. 2017;14:5591-5596.

### Novel cancer genes positively selected for missense mutations

##### Aurora kinase A encoded by the *AURKA* gene

AURKA, also known as a Breast tumor-amplified kinase, is a mitotic serine/threonine kinase that contributes to the regulation of cell cycle progression. It associates with the centrosome and the spindle microtubules during mitosis and plays a critical role in various mitotic events.

In harmony with the notion that AURKA's mutation pattern reflects a pro-oncogenic role for the protein, elevated expression of AURKA has been shown to induce oncogenic phenotypes (Takahahi *et al*., 2015, Treekitkarnmongkol *et al*., 2016).

Similarly, the observation that downregulation, inhibition or depletion of AURKA reduced viability and invasiveness of cancer cells (Sillars-Hardebol *et al*., 2012, Li *et al*. 2018, van Gijn *et al*., 2019) also argues for an oncogenic role of the protein.

Significantly, specific knockdown of *AURKA* in cultured pancreatic cancer cells strongly suppressed *in vitro* cell growth and *in vivo* tumorigenicity (Hata *et al*, 2005). Recently a novel AURKA mutation (V352I) was identified from clinical specimens and it was shown that AURKA (V352I)-induced carcinogenesis was earlier and much more severe than wild-type AURKA, implying that the V352I mutation may accelerate cancer progression (Su *et al*., 2019).

Although many *AURKA* mutations were identified in cancer patients, it is noteworthy that there is no evidence for the clustering or 'recurrence' of mutations. The most likely explanation for the lack of clustering of mutations is that since AURKA interacts with numerous proteins (e.g. PIFO, GADD45A, AUNIP, NIN, FRY, SIRT2, MYCN, HNRNPU, AAAS, KLHL18, CUL3, FOXP1) there may be multiple sites where missense mutations affecting these interactions may result in disregulation of the activity of AURKA.

In summary, although all the available experimental information argues for an oncogenic role of *AURKA*, there was no evidence for the clustering of its missense mutations. In our view this observation illustrates that recurrence of missense mutations is not a *sine qua non* criterion of oncogenes.

Recent studies have also revealed that AURKA and TWIST1 are linked in a feedback loop controlling tumorigenesis and metastasis. AURKA phosphorylates TWIST1, inhibits its ubiquitylation, increases its transcriptional activity and favors its homodimerization. TWIST1 prevents AURKA degradation, thereby triggering a feedback loop. Ablation of either AURKA or TWIST1 completely inhibits epithelial-to-mesenchymal transition, suggesting that inhibition of AURKA and TWIST1 are synergistic in inhibiting tumorigenesis and metastasis (Wang *et al*., 2017).

Although the *TWIST1* gene is not present in the datasets (**Supplementary files 3 and 6)** that contain the metadata for transcripts containing at least 100 confirmed somatic, non polymorphic mutations identified in tumor tisses, inspection of the primary dataset (**Supplementary file 2)** indicates that it is characterized by very high value of rSMN (**Supplementary file 3**), indicating strong signature of purifying selection (see section on **Negatively selected genes**) consistent with the view that – in synergism with AURKA – it plays an important role in promoting tumorigenesis.

Hata T, Furukawa T, Sunamura M, Egawa S, Motoi F, Ohmura N, Marumoto T, Saya H, Horii A. RNA interference targeting aurora kinase a suppresses tumor growth and enhances the taxane chemosensitivity in human pancreatic cancer cells. Cancer Res. 2005; 65:2899-2905

Li X1,2, Xu W1, Kang W3, Wong SH1, Wang M4, Zhou Y4, Fang X4, Zhang X4, Yang H4,5, Wong CH6, To KF3, Chan SL6, Chan MTV7, et al. Genomic analysis of liver cancer unveils novel driver genes and distinct prognostic features. Theranostics. 2018; 8:1740-1751.

Sillars-Hardebol AH, Carvalho B, Tijssen M, Beliën JA, de Wit M, Delis-van Diemen PM, Pontén F, van de Wiel MA, Fijneman RJ, Meijer GA. TPX2 and AURKA promote 20q amplicon-driven colorectal adenoma to carcinoma progression. Gut. 2012; 61:1568-1575.

Su ZL, Su CW, Huang YL, Yang WY, Sampurna BP, Ouchi T, Lee KL, Wu CS, Wang HD, Yuh CH. A Novel AURKA Mutant-Induced Early-Onset Severe Hepatocarcinogenesis Greater than Wild-Type via Activating Different Pathways in Zebrafish. Cancers (Basel). 2019; 11. pii: E927.

Takahashi Y, Sheridan P, Niida A, Sawada G, Uchi R, Mizuno H, Kurashige J, Sugimachi K, Sasaki S, Shimada Y, Hase K, Kusunoki M, Kudo S, et al. The AURKA/TPX2 axis drives colon tumorigenesis cooperatively with MYC. Ann Oncol. 2015; 26:935-942.

Treekitkarnmongkol W, Katayama H, Kai K, Sasai K, Jones JC, Wang J, Shen L, Sahin AA, Gagea M, Ueno NT, Creighton CJ, Sen S. Aurora kinase-A overexpression in mouse mammary epithelium induces mammary adenocarcinomas harboring genetic alterations shared with human breast cancer. Carcinogenesis. 2016; 37:1180-1189.

van Gijn SE, Wierenga E, van den Tempel N, Kok YP, Heijink AM, Spierings DCJ, Foijer F, van Vugt MATM, Fehrmann RSN. TPX2/Aurora kinase A signaling as a potential therapeutic target in genomically unstable cancer cells. Oncogene. 2019; 38:852-867.

Wang J, Nikhil K, Viccaro K, Chang L, Jacobsen M, Sandusky G, Shah K. The Aurora-A-Twist1 axis promotes highly aggressive phenotypes in pancreatic carcinoma. J Cell Sci. 2017; 130:1078-1093.

#### Cyclin-dependent kinase 8, encoded by the *CDK8* gene

The *CDK8* gene is a coactivator involved in regulated gene transcription of nearly all RNA polymerase II-dependent genes.

*CDK8* is a colorectal cancer oncogene that regulates beta-catenin activity. Suppression of CDK8 expression inhibits proliferation in colon cancer cells characterized by high levels of CDK8 and beta-catenin hyperactivity (Firestein *et al.*, 2008). CDK8 has been shown to promote SMAD1-driven epithelial-to-mesenchymal transition through YAP1 recruitment (Serrao *et al.*, 2018). There is a large body of evidence that CDK8 is a key oncogenic driver in many cancers (Philip *et al.*, 2018). *CDK8* was found to be amplified or overexpressed in many colon cancers and CDK8 expression correlated with shorter patient survival (Liang *et al.*, 2018)**.**

Firestein R, Bass AJ, Kim SY, Dunn IF, Silver SJ, Guney I, Freed E, Ligon AH, Vena N, Ogino S, Chheda MG, Tamayo P, Finn S et al. CDK8 is a colorectal cancer oncogene that regulates beta-catenin activity. Nature. 2008; 455:547-551.

Liang J, Chen M, Hughes D, Chumanevich AA, Altilia S, Kaza V, Lim CU, Kiaris H, Mythreye K, Pena MM, Broude EV, Roninson IB. CDK8 Selectively Promotes the Growth of Colon Cancer Metastases in the Liver by Regulating Gene Expression of TIMP3 and Matrix Metalloproteinases. Cancer Res. 2018; 78:6594-6606

Philip S, Kumarasiri M, Teo T, Yu M, Wang S. Cyclin-Dependent Kinase 8: A New Hope in Targeted Cancer Therapy? J Med Chem. 2018; 61:5073-5092

Serrao A, Jenkins LM, Chumanevich AA, Horst B, Liang J, Gatza ML, Lee NY, Roninson IB, Broude EV, Mythreye K. Mediator kinase CDK8/CDK19 drives YAP1-dependent BMP4-induced EMT in cancer. Oncogene. 2018; 37:4792-4808.

#### Isocitrate dehydrogenase [NAD] subunit beta, mitochondrial, encoded by the *IDH3B* gene

IDH3B plays an essential role in the activity of isocitrate dehydrogenase. The heterodimer composed of the alpha (IDH3A) and beta (IDH3B) subunits and the heterodimer composed of the alpha (IDH3A) and gamma (IDH3G) subunits, have significant activity but the full activity of the heterotetramer (containing two subunits of IDH3A, one of IDH3B and one of IDH3G) requires the assembly of both heterodimers.

Our Pubmed search failed to identify publications with major relevance for the role of *IDH3B* in carcinogenesis. It is noteworthy, however, that the *IDH3B* gene contains recurrent somatic missense mutations at residue R131 that is equevalent with R132 and R140 of the paralogous enzymes, IDH1 and IDH2, respectively, that are affected by recurrent oncogenic missense mutations. These mutations of IDH1 and IDH2 result in loss of normal enzymatic function and the abnormal production of 2-hydroxyglutarate. 2-hydroxyglutarate has been found to inhibit enzymatic function of many alpha-ketoglutarate dependent enzymes, including histone and DNA demethylases, causing widespread epigenetic changes in the genome thereby promoting tumorigenesis. It seems likely that the R131 mutations of IDH3B may conntribute to carcinogenesis by a similar mechanism.

#### E3 ubiquitin-protein ligase MARCH7, encoded by the *MARCH7* gene.

March7 is an E3 ubiquitin-protein ligase, an enzyme that accepts ubiquitin from an E2 ubiquitin-conjugating enzyme and then directly transfer the ubiquitin to targeted substrates.

Several studies support an oncogenic role for the ubiquitin E3 ligase MARCH7. Studies on ovarian tissues have revealed that expression of MARCH7 was higher in ovarian cancer tissues than normal ovarian tissues. Silencing *MARCH7* decreased, whereas ectopic expression of MARCH7 increased cell proliferation, migration and invasion, suggesting that MARCH7 is oncogenic and a potential target for ovarian cancer therapy (Hu *et al*., 2015). The expression of MARCH7 was significantly higher in cervical cancer tissues than normal cervical tissues, suggesting that this oncogene may also serves as a potential target for cervical cancer therapy (Hu *et al*., 2018).

The expression level of MARCH7 in endometrial cancer tissues was also found to be significantly higher than that in normal endometrium tissues, suggesting that it may be an oncogenic factor in endometrial cancer (Liu *et al*., 2019). The oncogenic role of MARCH7 is supported by the fact its knockdown inhibited the invasion and metastasis of endometrial cancer cells *in vitro* and *in vivo*, whereas the opposite effect was observed after overexpressing MARCH7.

Hu J, Meng Y, Zeng J, Zeng B, Jiang X. Ubiquitin E3 Ligase MARCH7 promotes proliferation and invasion of cervical cancer cells through VAV2-RAC1-CDC42 pathway. Oncol Lett. 2018; 16:2312-2318.

Hu J, Meng Y, Yu T, Hu L, Mao M. Ubiquitin E3 ligase MARCH7 promotes ovarian tumor growth. Oncotarget. 2015; 6:12174-12187.

Liu L, Hu J, Yu T, You S, Zhang Y, Hu L. miR-27b-3p/MARCH7 regulates invasion and metastasis of endometrial cancer cells through Snail-mediated pathway. Acta Biochim Biophys Sin (Shanghai). 2019; 51:492-500.

#### GTP-binding protein RIT1, encoded by the *RIT1* gene

The high value of rMSN reflects primarily the recurrence of substitutions (Met90Ile, Met90Val) of Met90 of RIT1 protein.

RIT1 plays a crucial role in the activation of MAPK signaling cascades that mediate a wide variety of cellular functions, including cell proliferation, survival, and differentiation.

Since the Met90Ile substitution has been shown to result in an increased MAPK-ERK signaling (Aoki *et al*., 2013, Koenighofer *et al*., 2016), it is plausible to assume that the high rate of missense mutations reflects positive selection of oncogenic driver mutations.

In harmony with this conclusion, studies on endometrial cancer have revealed that RIT1 mRNA and protein were significantly overexpressed in endometrial cancer cell lines and in endometrial cancer tissues compared to non-cancerous endometrial tissue samples (Xu *et al*., 2015). Elevated expression of RIT1 was significantly correlated with pathological type, clinical stage. Kaplan-Meier survival analysis indicated that RIT1 expression was associated with poor overall survival of endometrial cancer patients, suggesting that elevated expression of RIT1 may contribute to the progression of endometrial cancer.

In a study of lung adenocarcinoma cases, several somatic mutations (including Met90Ile) were identified in the *RIT1* gene that were found to cluster in a hotspot near the switch II domain of the GTPase protein (Berger *et al*., 2014). Ectopic expression of these mutated *RIT1* genes was found to induce cellular transformation *in vitro* and *in vivo*, confirming that these substitutions are driver mutations and that *RIT1* is an oncogene in lung adenocarcinoma.


Aoki Y, Niihori T, Banjo T, Okamoto N, Mizuno S, Kurosawa K, Ogata T, Takada F, Yano M, Ando T, Hoshika T, Barnett C, Ohashi H, et al. Gain-of-function mutations in RIT1 cause Noonan syndrome, a RAS/MAPK pathway syndrome. Am. J. Hum. Genet.2013; 93:173-180

Berger AH, Imielinski M, Duke F, Wala J, Kaplan N, Shi GX, Andres DA, Meyerson M. Oncogenic RIT1 mutations in lung adenocarcinoma. Oncogene. 2014; 33:4418-4423.

Koenighofer M, Hung CY, McCauley JL, Dallman J, Back EJ, Mihalek I, Gripp KW, Sol-Church K, Rusconi P, Zhang Z, Shi GX, Andres DA, Bodamer OA. Mutations in RIT1 cause Noonan syndrome - additional functional evidence and expanding the clinical phenotype. Clin. Genet. 2016; 89:359-366

Xu F, Sun S, Yan S, Guo H, Dai M, Teng Y. Elevated expression of RIT1 correlates with poor prognosis in endometrial cancer. Int J Clin Exp Pathol. 2015; 8:10315-10324.


#### Yes-associated protein 1, encoded by the *YAP1* gene

Yes-associated protein 1 is known to be the critical downstream regulatory target in the Hippo signaling pathway that plays a pivotal role in tumor suppression by restricting proliferation and promoting apoptosis. This pathway is composed of a kinase cascade that eventually inactivates YAP1 since phosphorylation of YAP1 by the tumor suppressors LATS1/2 inhibits its translocation into the nucleus.

Several lines of evidence indicate that YAP1 is an oncogene. *YAP1* was found to act as oncogenic target of 11q22 amplification in multiple cancer subtypes, whereas *YAP1* silencing significantly decreases cell proliferation (Lorenzetto *et al*., 2014, Hamanaka *et al*., 2019). *YAP1* was shown to promote growth of prostate cancer, whereas knock down of its expression or inhibition of YAP1 function significantly suppressed tumor recurrence (Jiang *et al*., 2017). The key role of YAP1 in carcinogenesis is also supported by the fact that the tumor suppressor LATS2 inhibits the malignant behaviors of glioma cells by inactivating of YAP1 (Shi *et al*., 2019).

Although several *YAP1* mutations were identified in cancer patients, there is no evidence for the clustering or 'recurrence' of mutations. Similarly to the case of AURKA (see above), the most plausible explanation for the lack of clustering of mutations of this oncogene is that since YAP1 interacts with several proteins (e.g. YES kinase, LATS1, LATS2, TP73, RUNX1, WBP1, WBP2, TEAD1, TEAD2, TEAD3, TEAD4, HCK, MAPK8, MAPK9, CK1, ABL1) mutations at several different sites may affect these interactions and may result in disregulation of the activity of YAP1. In our view the cases of AURKA, YAP1 and YES1 illustrate that recurrence of missense mutations is not a *sine qua non* criterion of oncogenes.

Hamanaka N, Nakanishi Y, Mizuno T, Horiguchi-Takei K, Akiyama N, Tanimura H, Hasegawa M, Satoh Y, Tachibana Y, Fujii T, Sakata K, Ogasawara K, Ebiike H, et al. YES1 is a targetable oncogene in cancers harboring *YES1* gene amplification. Cancer Res. 2019; 79:5734-5745.
Jiang N, Ke B, Hjort-Jensen K, Iglesias-Gato D, Wang Z, Chang P, Zhao Y, Niu X, Wu T, Peng B, Jiang M, Li X, Shang Z, et al. YAP1 regulates prostate cancer stem cell-like characteristics to promote castration resistant growth. Oncotarget. 2017; 8:115054-115067.

Lorenzetto E, Brenca M, Boeri M, Verri C, Piccinin E, Gasparini P, Facchinetti F, Rossi S, Salvatore G, Massimino M, Sozzi G, Maestro R, Modena P. YAP1 acts as oncogenic target of 11q22 amplification in multiple cancer subtypes. Oncotarget. 2014; 5:2608-2621.

Shi Y, Geng D, Zhang Y, Zhao M, Wang Y, Jiang Y, Yu R, Zhou X. LATS2 Inhibits Malignant Behaviors of Glioma Cells via Inactivating YAP. J Mol Neurosci. 2019; 68:38-48.

#### Tyrosine-protein kinase Yes, encoded by the *YES1* gene

Tyrosine-protein kinase Yes (also known as proto-oncogene c-Yes) is a multidomain non-receptor protein tyrosine kinase containing an SH3 domain, an SH2 domain and a protein kinase domain. YES1 is involved in the regulation of cell growth and survival, apoptosis, cell-cell adhesion, cytoskeleton remodeling, and differentiation. It plays a role in cell cycle progression by phosphorylating the cyclin-dependent kinase 4/CDK4 thus regulating the G1 phase. YES1 has been shown to phosphorylate YAP1, leading to the localization of a YAP1-TBX-β-catenin complex to the promoters of antiapoptotic genes, thereby promoting carcinogenes (Rosenbluh *et al*., 2012). A small-molecule inhibitor of YES1 impeded the proliferation of β-catenin-dependent cancers in both cell lines and animal models.

Several lines of evidence have established an oncogenic role for *YES1*.

It has been demonstrated recently that YES1 is essential for lung cancer growth and progression in non-small cell lung cancer, suggesting that it is a promising therapeutic target in lung cancer. YES1 overexpression induced metastatic spread in preclinical *in vivo* models. whereas *YES1* genetic depletion by CRISPR/Cas9 technology significantly reduced tumor growth and metastasis (Garmendia *et al*., 2019).

In harmony with an oncogenic role of *YES1*, several microRNAs have been shown to inhibit the proliferation of tumor cells by targeting *YES1* (Tan, Lim and Tan, 2015, Shen *et al*., 2019, Zhao *et al*., 2020).

The oncogenic role of YES1 in cancer is also supported by the observation that it is amplified in several types of cancer, suggesting that it could be an attractive target for a cancer drug (Fan *et al*., 2018, Hamanaka *et al*., 2019). Hamanaka *et al*., (2019) have generated a YES1 kinase inhibitor, and have shown that YES1 kinase inhibition by this drug led to antitumor activity against *YES1*-amplified cancers *in vitro* and *in vivo*. The authors have also shown that Yes-associated protein 1 (YAP1) played a role downstream of YES1 and contributed to the growth of *YES1*-amplified cancers. indicating that the regulation of YAP1 by YES1 plays an important role in *YES1*-amplified cancers. These findings identify YES1 as a targetable oncogene of significant potential for clinical utility (Rai, 2019).

Although *YES1* contains an increased proportion of nonsynonymous mutations there is no evidence for the clustering or 'recurrence' of mutations. Similarly to the cases of *AURKA* and *YAP1* (see above), the most plausible explanation for the lack of clustering of mutations of this oncogene is that since YES1 is a multidomain protein that interacts with several proteins, mutations at several different sites may affect these interactions and may result in disregulation of the activity of YAP1. In our view the cases of AURKA, YAP1 and YES1 illustrate that recurrence of missense mutations is not a *sine qua non* criterion of oncogenes.

Fan PD, Narzisi G, Jayaprakash AD, Venturini E, Robine N, Smibert P, Germer S, Yu HA, Jordan EJ, Paik PK, Janjigian YY, Chaft JE, Wang L et al. YES1 amplification is a mechanism of acquired resistance to EGFR inhibitors identified by transposon mutagenesis and clinical genomics. Proc Natl Acad Sci U S A. 2018; 115:E6030-E6038.

Garmendia I, Pajares MJ, Hermida-Prado F, Ajona D, Bértolo C, Sainz C, Lavín A, Remírez AB, Valencia K, Moreno H, Ferrer I, Behrens C, Cuadrado M et al. YES1 Drives Lung Cancer Growth and Progression and Predicts Sensitivity to Dasatinib. Am J Respir Crit Care Med. 2019; 200:888-899.

Hamanaka N, Nakanishi Y, Mizuno T, Horiguchi-Takei K, Akiyama N, Tanimura H, Hasegawa M, Satoh Y, Tachibana Y, Fujii T, Sakata K, Ogasawara K, Ebiike H etal. YES1 Is a Targetable Oncogene in Cancers Harboring YES1 Gene Amplification. Cancer Res. 2019; 79:5734-5745.

Rai K. Personalized Cancer Therapy: YES1 Is the New Kid on the Block.
Cancer Res. 2019; 79:5702-5703.

Rosenbluh J, Nijhawan D, Cox AG, Li X, Neal JT, Schafer EJ, Zack TI, Wang X, Tsherniak A, Schinzel AC, Shao DD, Schumacher SE, Weir BA et al. β-Catenin-driven cancers require a YAP1 transcriptional complex for survival and tumorigenesis. Cell. 2012; 151:1457-1473.

Shen Y, Chen F, Liang Y. MicroRNA-133a inhibits the proliferation of non-small cell lung cancer by targeting *YES1*.Oncol Lett. 2019; 18:6759-6765.

Tan W, Lim SG, Tan TM. Up-regulation of microRNA-210 inhibits proliferation of hepatocellular carcinoma cells by targeting YES1. World J Gastroenterol. 2015; 21:13030-13041.

Zhao S, Jie C, Xu P, Diao Y. MicroRNA-140 inhibit prostate cancer cell invasion and migration by targeting YES proto-oncogene 1. J Cell Biochem. 2020;121:482-488.

### Negatively selected tumor essential genes

##### Atypical chemokine receptor 3, , encoded by the *ACKR3* (*CXCR7*) gene

ACKR3 is a member of the group of chemokine receptors that acts as a receptor for chemokines CXCL11 and CXCL12/SDF1. It is activated by CXCL11 in malignant hemapoietic cells, leading to phosphorylation of ERK1/2 (MAPK3/MAPK1) and enhanced cell adhesion and migration.

*ACKR3* is a known cancer gene, from Tier 1 of the Cancer Gene Census; it has a cancer hallmark annotation. Its importance in carcinogenesis is underlined by the fact that high expression of ACKR3 is associated with poor survival in several types of cancer.

As to the role of ACKR3 in hallmarks of cancer: it has been suggested that ACKR3 promotes proliferative signaling, angiogenesis, evasion of programmed cell death and invasion and metastasis.

Several studies support the key role of ACKR3 in tumor invasion and metastasis (Li *et al*., 2014, Stacer *et al*., 2016, Zhao *et al*., 2017, Puddinu *et al*., 2017, Melo et al., 2018, Qian *et al*., 2018). Since knock-down or pharmacological inhibition of *ACKR3* has been shown to reduce tumor invasion and metastasis, ACKR3 is a promising therapeutic target for the control of tumor dissemination.

Li XX, Zheng HT1, Huang LY, Shi DB, Peng JJ, Liang L, Cai SJ. Silencing of *CXCR7* gene represses growth and invasion and induces apoptosis in colorectal cancer through ERK and β-arrestin pathways. Int J Oncol. 2014; 45:1649-5167

Melo RCC, Ferro KPV, Duarte ADSS, Olalla Saad ST. CXCR7 participates in CXCL12-mediated migration and homing of leukemic and normal hematopoietic cells. Stem Cell Res Ther. 2018; 9:34.

Puddinu V, Casella S, Radice E, Thelen S, Dirnhofer S, Bertoni F, Thelen M. ACKR3 expression on diffuse large B cell lymphoma is required for tumor spreading and tissue infiltration. Oncotarget. 2017; 8:85068-85084.

Qian T, Liu Y, Dong Y, Zhang L, Dong Y, Sun Y, Sun D. CXCR7 regulates breast tumor metastasis and angiogenesis in vivo and in vitro. Mol Med Rep. 2018; 17:3633-3639.

Stacer AC, Fenner J, Cavnar SP, Xiao A, Zhao S, Chang SL, Salomonnson A, Luker KE, Luker GD. Endothelial CXCR7 regulates breast cancer metastasis. Oncogene. 2016; 35:1716-1724.

Zhao ZW, Fan XX, Song JJ, Xu M, Chen MJ, Tu JF, Wu FZ, Zhang DK, Liu L, Chen L, Ying XH, Ji JS. ShRNA knock-down of CXCR7 inhibits tumour invasion and metastasis in hepatocellular carcinoma after transcatheter arterial chemoembolization. J Cell Mol Med. 2017; 21:1989-1999.

#### CX3C chemokine receptor 1, encoded by the *CX3CR1* gene

CX3CR1 is a member of the group of chemokine receptors that play a major role in tumor metastasis. The interactions of chemokines, also known as chemotactic cytokines, with their receptors regulate immune and inflammatory responses. However, recent studies have demonstrated that cancer cells subvert the normal chemokine role, transforming them into fundamental constituents of the tumor microenvironment with tumor-promoting effects. CX3CR1 is the receptor for the CX3C chemokine fractalkine (CX3CL1) that mediates both its adhesive and migratory functions.

CX3CR1 expression has been shown to be associated with the process of cellular migration *in vitro* and tumor metastasis of clear cell renal cell carcinoma *in vivo* (Yao et et., 2014).

Recent studies indicate that tumor-associated macrophages MΦ can influence cancer progression and metastasis and that CCR2 and CX3CR1 play important roles in metastasis. Schmall *et al*. (2015) have shown that coculturing of tumor-associated macrophages with mouse Lewis lung carcinoma caused up-regulation of CCR2/CCL2 and CX3CR1/CX3CL1 in both the cancer cells and the macrophages. In vivo, MΦ depletion and genetic ablation of *CCR2* and *CX3CR1* all inhibited LLC1 tumor growth and metastasis, and enhanced survival. Furthermore, mice treated with CCR2 antagonist mimicked genetic ablation of CCR2, showing reduced tumor growth and metastasis. These findings indicate that tumor-associated MΦ play a central role in lung cancer growth and metastasis, with bidirectional cross-talk between MΦ and cancer cells via CCR2 and CX3CR1 signaling. These studies suggest that the therapeutic strategy of blocking CCR2 and CX3CR1 may prove beneficial for halting metastasis.

CX3CR1 is highly expressed in gastric cancer tissues and is related to lymph node metastasis and larger tumor size. CX3CR1 overexpression promoted gastric cancer cell migration, invasion, proliferation and survival (Wei *et al*., 2015).

CX3CR1 is overexpressed in human breast tumors and cancer cells utilize the chemokine receptor CX3CR1 to exit the blood circulation and metastize to the skeleton. To assess the clinical potential of targeting CX3CR1 in breast cancer Shen *et al*., (2016) have used neutralizing antibody for this receptor, transcriptional suppression by CRISPR interference as well as a potent and selective small-molecule antagonist of CX3CR1 in preclinical animal models of metastasis. The authors have found that inactivation of CX3CR1 impairs the lodging of circulating tumor cells to the skeleton and impairs further growth of established metastases. These data suggest that CX3CR1 has a important role in promoting metastasis_activity and that CX3CR1 antagonists may valuable as drugs of tumor therapy.

Schmall A, Al-Tamari HM, Herold S, Kampschulte M, Weigert A, Wietelmann A, Vipotnik N, Grimminger F, Seeger W, Pullamsetti SS, Savai R. Macrophage and cancer cell cross-talk via CCR2 and CX3CR1 is a fundamental mechanism driving lung cancer. Am J Respir Crit Care Med. 2015; 191:437-447.

Shen F, Zhang Y, Jernigan DL, Feng X, Yan J, Garcia FU, Meucci O, Salvino JM, Fatatis A. Novel Small-Molecule CX3CR1 Antagonist Impairs Metastatic Seeding and Colonization of Breast Cancer Cells. Mol Cancer Res. 2016; 14:518-527.

Wei LM, Cao S, Yu WD, Liu YL, Wang JT. Overexpression of CX3CR1 is associated with cellular metastasis, proliferation and survival in gastric cancer. Oncol Rep. 2015; 33:615-624.

Yao X, Qi L, Chen X, Du J, Zhang Z, Liu S. Expression of CX3CR1 associates with cellular migration, metastasis, and prognosis in human clear cell renal cell carcinoma. Urol Oncol. 2014; 32:162-170.

**C-C chemokine receptor type 2, encoded by the *CCR2* gene**
**C-C chemokine receptor type 5, encoded by the *CCR5* gene**

Although the *CCR2* gene of C-C chemokine receptor type 2 and the *CCR5* gene of C-C chemokine receptor type 5 are not present in the CG_SO and CG_SSI lists defined by the 2SD cut-off values they are also characterized by very high values of rSNM (**Supplementary file 3**), suggesting that they may also play importanct roles in tumor metastasis.

CCR2 is the key functional receptor for the chemokine ligand CCL2. Its binding with CCL2 on monocytes and macrophages mediates chemotaxis and migration induction. Recent studies indicate that CCR2 and CX3CR1 play important roles in metastasis (Schmall *et al*. 2015). The CCL2-CCR2 signaling axis has generated increasing interest in recent years due to its association with the progression of cancer.. The CCL2-CCR2, signaling pair has been shown to have multiple pro-tumorigenic roles, mediating tumor growth and angiogenesis (Lim *et al*., 2016).

CCR5 serves as a receptor for a number of inflammatory CC-chemokines including CCL3/MIP-1-alpha, CCL4/MIP-1-beta. Recent studies have revealed that C-C chemokine receptor type 5 plays a key role in progression of tumorigenesis, Expression of CCR5 augments regulatory T cell differentiation and migration to sites of inflammation. The misexpression of CCR5 in epithelial cells, induced upon oncogenic transformation, hijacks this migratory phenotype (Aldinucci and Casagrande, 2018, Jiao *et al*., 2019).

Aldinucci D, Casagrande N. Inhibition of the CCL5/CCR5 Axis against the Progression of Gastric Cancer. Int J Mol Sci. 2018; 19. pii: E1477.

Jiao X, Nawab O, Patel T, Kossenkov AV, Halama N, Jaeger D, Pestell RG. Recent Advances Targeting CCR5 for Cancer and Its Role in Immuno-Oncology. Cancer Res. 2019; 79:4801-4807.

Lim SY, Yuzhalin AE, Gordon-Weeks AN, Muschel RJ. Targeting the CCL2-CCR2 signaling axis in cancer metastasis. Oncotarget. 2016; 7:28697-28710.

Schmall A, Al-Tamari HM, Herold S, Kampschulte M, Weigert A, Wietelmann A, Vipotnik N, Grimminger F, Seeger W, Pullamsetti SS, Savai R. Macrophage and cancer cell cross-talk via CCR2 and CX3CR1 is a fundamental mechanism driving lung cancer. Am J Respir Crit Care Med. 2015; 191:437-447

#### Dentin sialophosphoprotein, encoded by the *DSPP* gene

The *DSPP* gene has been selected as a gene showing very high values of rSMN, suggesting negative selection of missense and nonsense mutations (**Supplementary file 3**). It must be pointed out that based on the high silent/missense ratio *DSPP* has also been identified by others as a gene showing signs of strong negative selection (Zhou *et al*., 2017).

Dentin sialophosphoprotein is a secreted protein that has been shown to play an important role in dentinogenesis. It binds high amount of calcium and facilitates initial mineralization of dentin matrix collagen as well as regulate the size and shape of the crystals, therefore it seemed surprising that its gene would qualify as a negatively selected tumor essential gene.

There is evidence in the scientific literature that the protein may have a tumorigenic role in oral cancer (Chaplet *et al*., 2006, Johi *et al*., 2010, Saxena *et al*., 2015, Gkouveris *et al*., 2018), Nikitakis *et al*., 2018). Nevertheless, the high silent to missense rate is not a reflection of the importance of *DSPP* for carcinogenesis. The *DSPP* gene contains a 2-kb repeat domain containing over 200 tandem copies of a nominal 9-basepair (AGC AGC GAC) repeat encoding a series of tandem Ser Ser Asp repeats and the unusully high rate of silent mutations is restricted to this region of the gene.

A study of 188 normal human chromosomes revealed that the repeat domain of *DSPP* is hypervariable with extraordinary rates of change including slip-replication indel events and predominantly C-to-T transition SNPs (McKnight *et al*., 2008). In harmony with the increased rate and predominance of C-to-T transition in the AGC AGC GAC (Ser-Ser-Asp) repeats, the vast majority of substitutions in this repeat region of the *DSSP* gene are silent. The unusually high silent to missense mutation ratio of the *DSPP* gene is thus not due to purifying selection of a tumor essential gene.

Chaplet M, Waltregny D, Detry C, Fisher LW, Castronovo V, Bellahcène A. Expression of dentin sialophosphoprotein in human prostate cancer and its correlation with tumor aggressiveness. Int J Cancer. 2006; 118:850-856.

Gkouveris I, Nikitakis NG, Aseervatham J, Ogbureke KUE. The tumorigenic role of DSPP and its potential regulation of the unfolded protein response and ER stress in oral cancer cells. Int J Oncol. 2018; 53:1743-1751.

Joshi R, Tawfik A, Edeh N, McCloud V, Looney S, Lewis J, Hsu S, Ogbureke KU. Dentin sialophosphoprotein (DSPP) gene-silencing inhibits key tumorigenic activities in human oral cancer cell line, OSC2. PLoS One. 2010; 5:e13974.

McKnight DA, Suzanne Hart P, Hart TC, Hartsfield JK, Wilson A, Wright JT, Fisher LW. A comprehensive analysis of normal variation and disease-causing mutations in the human DSPP gene. Hum Mutat. 2008; 29:1392-404.

Nikitakis NG, Gkouveris I, Aseervatham J, Barahona K, Ogbureke KUE. DSPP-MMP20 gene silencing downregulates cancer stem cell markers in human oral cancer cells. Cell Mol Biol Lett. 2018; 23:30.

Saxena G, Koli K, de la Garza J, Ogbureke KU. Matrix metalloproteinase 20-dentin sialophosphoprotein interaction in oral cancer. J Dent Res. 2015; 94:584-593.

Zhou Z, Zou Y, Liu G, Zhou J, Wu J, Zhao S, Su Z, Gu X. Mutation-profile-based methods for understanding selection forces in cancer somatic mutations: a comparative analysis. Oncotarget. 2017;8:58835-58846.

#### Forkhead box protein G1, encoded by the *FOXG1*

FOXG1 is a member of the FOX (Forkhead box) protein family of transcription factors that play important roles in regulating the expression of genes involved in cell growth, proliferation, differentiation and longevity. FOXG1 localizes to mitochondria and coordinates cell differentiation and bioenergetics (Pancrazi *et al*., 2015).

The tumor promoting role of FOXG1 is supported by the observation that childhood medulloblastomas are characterized by 2-7-fold copy gain for *FOXG1*. *FOXG1* copy gain (>2 to 21 folds) was seen in 93% of a validating set of tumors and showed a positive correlation with protein expression (Adesina *et al*., 2007).

The oncogenic role of FOXG1 is also supported by the observation that a decrease of FOXG1 in medulloblastoma cells offers a survival advantage in mice (Adesina *et al*., 2015)., whereas high expression of FOXG1 was associated with poor survival of glioblastoma patients (Robertson *et al*., 2015).

The carcinogenesis promoting activity of FOXG1 is supported by the observation that endogenous FOXG1 expression levels were positively correlated to the glioblastoma multiforme disease progression (Wang *et al*., 2018). Overexpression of FOXG1 protein resulted in increased cell viability, and it was suggested that FOXG1 functions as an onco-factor by promoting proliferation and inhibiting differentiation..

Recent studies on glioblastoma have shown that transcription factors FOXG1 and TLE1 promote glioblastoma propagation by supporting maintenance of brain tumour-initiating cells (Dali *et al*., 2018). Since the expressions of caspase family members were significantly altered in response to change of FOXG1 expression, it has been suggested that FOXG1 also contributes to carcinogenesis as a negative regulator of glioma cell apoptosis (Chen *et al*., 2018).

Adesina AM, Nguyen Y, Mehta V, Takei H, Stangeby P, Crabtree S, Chintagumpala M, Gumerlock MK. FOXG1 dysregulation is a frequent event in medulloblastoma. J Neurooncol. 2007; 85:111-122.

Adesina AM, Veo BL, Courteau G, Mehta V, Wu X, Pang K, Liu Z, Li XN, Peters L. FOXG1 expression shows correlation with neuronal differentiation in cerebellar development, aggressive phenotype in medulloblastomas, and survival in a xenograft model of medulloblastoma. Hum Pathol. 2015; 46:1859-1871.

Chen J, Wu X, Xing Z, Ma C, Xiong W, Zhu X, He X. FOXG1 Expression Is Elevated in Glioma and Inhibits Glioma Cell Apoptosis. J Cancer. 2018; 9:778-783.

Dali R, Verginelli F, Pramatarova A, Sladek R, Stifani S. Characterization of a FOXG1:TLE1 transcriptional network in glioblastoma-initiating cells. Mol Oncol. 2018; 12:775-787

Pancrazi L, Di Benedetto G, Colombaioni L, Della Sala G, Testa G, Olimpico F, Reyes A, Zeviani M, Pozzan T, Costa M. Foxg1 localizes to mitochondria and coordinates cell differentiation and bioenergetics. Proc Natl Acad Sci U S A. 2015; 112:13910-13915.

Robertson E, Perry C, Doherty R, Madhusudan S. Transcriptomic profiling of Forkhead box transcription factors in adult glioblastoma multiforme. Cancer Genomics Proteomics. 2015; 12:103-112.

Wang L, Wang J, Jin T, Zhou Y, Chen Q. FoxG1 facilitates proliferation and inhibits differentiation by downregulating FoxO/Smad signaling in glioblastoma. Biochem Biophys Res Commun. 2018; 504:46-53.

#### Forkhead box protein P2, encoded by *FOXP2* gene

Forkhead box protein P2 (FOXP2) is a transcriptional repressor.

The role of FOXP2 in cancer is somewhat controversial, it appears to have oncogenic or tumor suppressor roles, depending on the cellular and histological features of tumors. While FOXP2 has been found to be down-regulated in breast cancer, hepatocellular carcinoma and gastric cancer biopsies, overexpressed FOXP2 has been reported in multiple myelomas,, several subtypes of lymphomas, as well as in neuroblastomas and some prostate cancers (Herrero *et al*., 2018).

Numerous recent studies indicate a tumor suppressor like role for FOXP2 (Campbell *et al*., 2010, Cuiffo *et al*., 2014, 2015, Yan *et al*., 2015, Diao *et al*., Song *et al*., 2017, Chen *et al*. 2018, Li *et al*., 2019), others present evidence for an oncogene-like role of the protein (Campbell *et al*., 2010, Zhong *et al*., 2017, Wu *et al*., 2018, Wang *et al*., 2019 ).

The high silent to missense ratio of substitution mutations observed in the case of the *FOXP2* gene does not seem to be a reflection of purifying selection, that might be in harmony of an oncogene-like role, but definitely not with a tumor suppressor role.

The translated region of the *FOXP2* gene contains a long stretch of CAG repeats (residiues 177-216), corresponding to the polyQ segment of the protein. Silent mutations are clustered in the polyQ tract of the protein encoded by the imperfect polymorphic region, suggesting that the increased silent to missense rate of substitutions in this gene has much less to do with purifying selection than with microsatellite instability.


Campbell AJ, Lyne L, Brown PJ, Launchbury RJ, Bignone P, Chi J, Roncador G, Lawrie CH, Gatter KC, Kusec R, Banham AH. Aberrant expression of the neuronal transcription factor FOXP2 in neoplastic plasma cells. Br J Haematol. 2010; 149:221–230.

Chen MT, Sun HF, Li LD, Zhao Y, Yang LP, Gao SP, Jin W. Downregulation of FOXP2 promotes breast cancer migration and invasion through TGFβ/SMAD signaling pathway. Oncol Lett. 2018; 15:8582-8588.

Cuiffo BG, Campagne A, Bell GW, Lembo A, Orso F, Lien EC, Bhasin MK, Raimo M, Hanson SE, Marusyk A, El-Ashry D, Hematti P, Polyak K, et al. MSC-regulated microRNAs converge on the transcription factor FOXP2 and promote breast cancer metastasis. Cell Stem Cell. 2014; 15:762–774.

Diao H, Ye Z, Qin R. miR-23a acts as an oncogene in pancreatic carcinoma by targeting FOXP2.
J Investig Med. 2018;66: 676-683.

Herrero MJ, Gitton Y. The untold stories of the speech gene, the FOXP2 cancer gene. Genes Cancer. 2018; 9:11-38

Li ZY, Zhang ZZ, Bi H, Zhang QD, Zhang SJ, Zhou L, Zhu XQ, Zhou J. Upregulated microRNA‑ 671‑ 3p promotes tumor progression by suppressing forkhead box P2 expression in non‑ small‑ cell lung cancer. Mol Med Rep. 2019; 20:3149-3159.

Song XL, Tang Y, Lei XH, Zhao SC, Wu ZQ. miR-618 Inhibits Prostate Cancer Migration and Invasion by Targeting FOXP2. J Cancer. 2017; 8:2501-2510.

Wang WX, Yu HL, Liu X. MiR-9-5p suppresses cell metastasis and epithelial-mesenchymal transition through targeting FOXP2 and predicts prognosis of colorectal carcinoma. Eur Rev Med Pharmacol Sci. 2019; 23:6467-6477.

Wu J, Liu P, Tang H, Shuang Z, Qiu Q, Zhang L, Song C, Liu L, Xie X, Xiao X. FOXP2 Promotes Tumor Proliferation and Metastasis by Targeting GRP78 in Triple-negative Breast Cancer. Curr Cancer Drug Targets. 2018; 18:382-389.

Yan X, Zhou H, Zhang T, Xu P, Zhang S, Huang W, Yang L, Gu X, Ni R, Zhang T. Downregulation of FOXP2 promoter human hepatocellular carcinoma cell invasion. Tumor Biol. 2015; 36:9611–9619.

Zhong C, Liu J, Zhang Y, Luo J, Zheng J. MicroRNA-139 inhibits the proliferation and migration of osteosarcoma cells via targeting forkhead-box P2. Life Sci. 2017; 191:68-73.


#### Glucose-6-phosphate 1-dehydrogenase, encoded by the *G6PD* gene

Glucose-6-phosphate 1-dehydrogenase catalyzes the rate-limiting step of the oxidative pentose-phosphate pathway, its main function is to provide reducing power (NADPH) and pentose phosphates for fatty acid and nucleic acid synthesis. There is strong support for the importance of G6PD for tumor growth. Progression of tumor cells to more aggressive phenotypes requires not only the upregulation of glygolysis but also the pentose phosphate pathway as a provider of reducing power and ribose phosphate to the cell for maintenance of redox balance and biosynthesis of nucleotides and lipids, making G6PD a promising target in cancer therapy (Zhang *et al*., 2014).

The key importance of G6PD for tumor growth is supported by the fact that elevated G6PD levels promote cancer progression in numerous tumor types, that high G6PD expression is a poor prognostic factor and that knockdown of G6PD suppresses cell viability and growth (Wang *et al*., 2012, Pu *et al*., 2015, Wang *et al*., 2015, Poulain *et al*., 2017, Chen *et al*., 2018, Yang *et al*., 2018, Barajas *et al*., 2018 Yang *et al*., 2019).

Barajas JM, Reyes R, Guerrero MJ, Jacob ST, Motiwala T, Ghoshal K.The role of miR-122 in the dysregulation of glucose-6-phosphate dehydrogenase (G6PD) expression in hepatocellular cancer. Sci Rep. 2018; 8:9105.

Chen X, Xu Z, Zhu Z, Chen A, Fu G, Wang Y, Pan H, Jin B. Modulation of G6PD affects bladder cancer via ROS accumulation and the AKT pathway in vitro. Int J Oncol. 2018; 53:1703-1712.

Poulain L, Sujobert P, Zylbersztejn F, Barreau S, Stuani L, Lambert M, Palama TL, Chesnais V, Birsen R, Vergez F, Farge T, Chenevier-Gobeaux C, Fraisse M, et al. High mTORC1 activity drives glycolysis addiction and sensitivity to G6PD inhibition in acute myeloid leukemia cells. Leukemia. 2017; 31:2326-2335.

Pu H, Zhang Q, Zhao C, Shi L, Wang Y, Wang J, Zhang M. Overexpression of G6PD is associated with high risks of recurrent metastasis and poor progression-free survival in primary breast carcinoma. World J Surg Oncol. 2015; 13:323.

Wang J, Yuan W, Chen Z, Wu S, Chen J, Ge J, Hou F, Chen Z. Overexpression of G6PD is associated with poor clinical outcome in gastric cancer. Tumour Biol. 2012; 33:95-101.

Wang X, Li X, Zhang X, Fan R, Gu H, Shi Y, Liu H. Glucose-6-phosphate dehydrogenase expression is correlated with poor clinical prognosis in esophageal squamous cell carcinoma. Eur J Surg Oncol. 2015; 41:1293-1299.

Yang CA, Huang HY, Lin CL, Chang JG. G6PD as a predictive marker for glioma risk, prognosis and chemosensitivity. J Neurooncol. 2018; 139:661-670.

Yang HC, Wu YH, Yen WC, Liu HY, Hwang TL, Stern A, Chiu DT. The Redox Role of G6PD in Cell Growth, Cell Death, and Cancer. Cells. 2019;8. pii: E1055.

Zhang C, Zhang Z, Zhu Y, Qin S. Glucose-6-phosphate dehydrogenase: a biomarker and potential therapeutic target for cancer. Anticancer Agents Med Chem. 2014; 14:280-289.

#### Mitogen-activated protein kinase 13, encoded by the *MAPK13* gene

MAPK13 (p38δ mitogen-activated protein kinase) is a serine/threonine kinase which acts as an essential component of the MAP kinase signal transduction pathway. MAPK13 plays an important role in the cascades of cellular responses evoked by extracellular stimuli such as proinflammatory cytokines. The protein is involved in the regulation of epidermal keratinocyte differentiation, apoptosis and skin tumor development.

Although MAPK13 shows signatures of negative selection that would suggest a pro-oncogenioc role for the protein, expermintal data are controversial as to its role in carcinogenesis: there is evidence for both a pro-oncogenic and tumor suppressor roles of MAPK13.

The observation that p38delta promotes cell proliferation and tumor development in epidermis suggests that it has a pro-oncogenic role (Schindler *et al.*, 2009). Analyzes of the gene expression profiles have shown that MAPK13 is expressed in uterine, ovary, stomach, colon, liver and kidney cancer tissues at higher levels compared with adjacent normal tissues. *MAPK13* gene knockdown has been shown to abrogate the tumor-initiating ability of cancer stem-like cells, indicating that the gene has a cancer-promoting role (Yasuda *et al*., 2016). The protein p38δ is highly expressed in all types of human breast cancers, whereas lack of p38δ resulted in reduced primary tumor size and blocked the metastatic potential to the lungs (Wada *et al*., 2017). The fact that mice with germline deletion of the p38δ gene are significantly protected from chemical skin carcinogenesis also suggests a cancer promoting role for the protein (Kiss *et al*., 2016). Interestingly, cell-selective targeted ablation of p38δ in keratinocytes and in immune (myeloid) cells on skin tumor development had different effects. Conditional keratinocyte-specific p38δ ablation reduced malignant progression in males and females relative to their wild-type counterparts. In contrast, conditional myeloid cell-specific p38δ deletion inhibited skin tumorigenesis in male but not female mice. These results reveal that cell-specific p38δ targeting modifies susceptibility to skin carcinogenesis in a context-, stage-, and sex-specific manner (Kiss *et al*., 2019).

The closely related MAPK14, MAPK12 and MAPK13 proteins are known to modulate the immune response, and since chronic inflammation is a known risk factor for tumorigenesis it seems possible that the role of MAPK13 in carcinogenesis may be associated with inflammation. Del Reino *et al*., (2014) have analyzed the role of MAPK12 and MAPK13 in colon cancer associated to colitis and have shown that the deficiency of MAPK12 and MAPK13 significantly decreased tumor formation, in parallel with a decrease in proinflammatory cytokine and chemokine production.

In contrast with the observations arguing for a pro-oncogenic role of the protein, loss of p38δ mitogen-activated protein kinase expression has been shown to promote oesophageal squamous cell carcinoma proliferation, migration and anchorage-independent growth, suggesting that it has a tumor suppressor role (O'Callaghan *et al*., 2013). Similarly, inactivation of the gene in lung cancer cells has been shown to lead to upregulation of the stemness proteins, thus promoting the cancer stem cell properties of these cells (Fang *et al*., 2017). Promoter methylation of *MAPK13* was found to be present in the majority of primary and metastatic melanomas. Restoration of MAPK13 expression in melanoma cells exhibiting epigenetic silencing of this gene reduced proliferation, indicative of tumor suppressive functions for the protein (Gao *et al*., 2013).

In summary, although MAPK13 plays both pro-oncogenic and tumor suppressor functions in different cellular processes our observation that during tumor evolution negative selection dominates for MAPK13 suggests that the selection pressure to preserve the tumor promoting activities of MAPK13 activity overrides the pressure to eliminate its tumor suppressor activities.

Del Reino P, Alsina-Beauchamp D, Escós A, Cerezo-Guisado MI, Risco A, Aparicio N, Zur R, Fernandez-Estévez M, Collantes E, Montans J, Cuenda A Pro-oncogenic role of alternative p38 mitogen-activated protein kinases p38γ and p38δ, linking inflammation and cancer in colitis-associated colon cancer.Cancer Res. 2014; 74:6150-6160.

Fang Y, Wang J, Wang G, Zhou C, Wang P, Zhao S, Zhao S, Huang S, Su W, Jiang P, Chang A, Xiang R, Sun P Inactivation of p38 MAPK contributes to stem cell-like properties of non-small cell lung cancer. Oncotarget. 2017; 8:26702-26717.

Gao L, Smit MA, van den Oord JJ, Goeman JJ, Verdegaal EM, van der Burg SH, Stas M, Beck S, Gruis NA, Tensen CP, Willemze R, Peeper DS, van Doorn R. Genome-wide promoter methylation analysis identifies epigenetic silencing of *MAPK13* in primary cutaneous melanoma. Pigment Cell Melanoma Res. 2013; 26:542-554.

Kiss A, Koppel AC, Anders J, Cataisson C, Yuspa SH, Blumenberg M, Efimova T. Keratinocyte p38δ loss inhibits Ras-induced tumor formation, while systemic p38δ loss enhances skin inflammation in the early phase of chemical carcinogenesis in mouse skin. Mol Carcinog. 2016; 55:563-574

Kiss A, Koppel AC, Murphy E, Sall M, Barlas M, Kissling G, Efimova T. Cell Type-Specific p38δ Targeting Reveals a Context-, Stage-, and Sex-Dependent Regulation of Skin Carcinogenesis. Int J Mol Sci. 2019; 20(7). pii: E1532.

O'Callaghan C, Fanning LJ, Houston A, Barry OP. Loss of p38δ mitogen-activated protein kinase expression promotes oesophageal squamous cell carcinoma proliferation, migration and anchorage-independent growth. Int J Oncol. 2013; 43:405-415.

Schindler EM, Hindes A, Gribben EL, Burns CJ, Yin Y, Lin MH, Owen RJ, Longmore GD, Kissling GE, Arthur JS, Efimova T. p38delta Mitogen-activated protein kinase is essential for skin tumor development in mice. Cancer Res. 2009; 69:4648-4655.

Wada M, Canals D, Adada M, Coant N, Salama MF, Helke KL, Arthur JS, Shroyer KR, Kitatani K, Obeid LM, Hannun YA. P38 delta MAPK promotes breast cancer progression and lung metastasis by enhancing cell proliferation and cell detachment. Oncogene. 2017; 36:6649-6657.

Yasuda K, Hirohashi Y, Kuroda T, Takaya A, Kubo T, Kanaseki T, Tsukahara T, Hasegawa T, Saito T, Sato N, Torigoe T MAPK13 is preferentially expressed in gynecological cancer stem cells and has a role in the tumor-initiation. Biochem Biophys Res Commun. 2016; 472:643-647.

#### Protein AF-9, encoded by the *MLLT3* gene

The *MLLT3* gene (present in CGC list of cancer genes) has been selected as a gene showing very high values of rSMN, suggesting negative selection of missense and nonsense mutations (**Supplementary file 8**).

It must be pointed out that based on the high silent/missense ratio *MLLT3* (as well as *TBP* and *DSPP*) has also been identified by others as a gene subject to negative selection (Zhou *et al*., 2017).

Protein AF-9 is a component of a complex required to increase the catalytic rate of RNA polymerase II transcription by suppressing transient pausing by the polymerase at multiple sites along the DNA.

Several studies indicate that *MLLT3* is a proto-oncogene, its inactivation or downregulation suppresses lymphoma cell proliferation, invasion and inhibits metastasis and proliferation of prostate cancer (Zhang *et al*., 2012, Meng *et al*., 2017).

Despite the tumor promoting role of *MLLT3*, the high silent to missense ratio of substitution mutations does not seem to be a reflection of strong negative selection. The translated region of the *MLLT3* gene contains a long stretch of AGC repeats (encoding the polyS segment of the protein, residues 149-194). The 'excess' of silent mutations are clustered in the polyS tract of the protein encoded by the imperfect polymorphic AGC microsatellite region of the *MLLT3* gene, that is known to be highly unstable (Walker *et al*., 1994).

Meng FJ, Meng FM, Wu HX, Cao XF. miR-564 inhibited metastasis and proliferation of prostate cancer by targeting MLLT3. Eur Rev Med Pharmacol Sci. 2017; 21:4828-4834.

Walker GJ, Walters MK, Palmer JM, Hayward NK. The *MLLT3* gene maps between D9S156 and D9S171 and contains an unstable polymorphic trinucleotide repeat. Genomics. 1994; 20:490-491.

Zhang T, Luo Y, Wang T, Yang JY. MicroRNA-297b-5p/3p target Mllt3/Af9 to suppress lymphoma cell proliferation, migration and invasion in vitro and tumor growth in nude mice. Leuk Lymphoma. 2012; 53:2033-2040.

Zhou Z, Zou Y, Liu G, Zhou J, Wu J, Zhao S, Su Z, Gu X. Mutation-profile-based methods for understanding selection forces in cancer somatic mutations: a comparative analysis. Oncotarget. 2017;8:58835-58846.

**Neuro-oncological ventral antigen 1 (Nova-1), encoded by the *NOVA1* gene**

Nova-1 is an RNA-binding protein involved in the regulation of RNA splicing.

The importance of Nova1 for tumor growth is supported by the observation that overexpressed intratumoral NOVA1 was associated with poor survival rate and increased recurrence rate of hepatocellular carcinoma (HCC) and was an independent prognostic factor for overall survival rate and tumor recurrence. HCC cell lines over-expressing NOVA1 exhibited greater potentials in cell proliferation, invasion and migration, while knockdown of NOVA1 had the opposite effects. All these findings indicate that NOVA1 may act as a prognostic marker for poor outcome and high recurrence in HCC (Zhang *et al*., 2014).

Similarly, NOVA1 expression was found to be up-regulated in melanoma samples and cell lines. and knockdown of NOVA1 suppressed melanoma cell proliferation, migration and invasion in both A375 and A875 cell lines These results suggested that NOVA1 acted as an oncogene in the development of melanoma (Yu *et al*., 2018).

Recent studies have shown that the tumor suppressor microRNA-592 suppresses the malignant phenotypes of thyroid cancer by downregulating NOVA1. Whereas overexpression of miR‑592 resulted in decreased cell proliferation, migration, and invasion in thyroid cancer, ectopic NOVA1 expression effectively abolished the tumor-suppressing effects of miR‑ 592 overexpression in thyroid cancer cells vitro and in vivo (Luo *et al*., 2019).

Recent studies have provided an explanation for the role of NOVA1 in carcinogenesis. Sayed *et al*., (2019) have shown that NOVA1 as well as the polypyrimidine-tract binding protein PTBP1 acts as enhancers of full-length TERT splicing, increasing telomerase activity, promoting telomere maintenance in cancer cells, thereby favoring theit replicative immortality.

Luo Y, Hao T, Zhang J, Zhang M, Sun P, Wu L. MicroRNA-592 suppresses the malignant phenotypes of thyroid cancer by regulating lncRNA NEAT1 and downregulating NOVA1. Int J Mol Med. 2019; 44:1172-1182.

Sayed ME, Yuan L, Robin JD, Tedone E, Batten K, Dahlson N, Wright WE, Shay JW, Ludlow AT. NOVA1 directs PTBP1 to hTERT pre-mRNA and promotes telomerase activity in cancer cells. Oncogene. 2019; 38:2937-2952

Yu X, Zheng H, Chan MTV, Wu WKK. NOVA1 acts as an oncogene in melanoma via regulating FOXO3a expression. J Cell Mol Med. 2018; 22:2622-2630.

Zhang YA, Zhu JM, Yin J, Tang WQ, Guo YM, Shen XZ, Liu TT. High expression of neuro-oncological ventral antigen 1 correlates with poor prognosis in hepatocellular carcinoma. PLoS One. 2014; 9:e90955.

**Calcium/calmodulin-dependent protein kinase type 1B, encoded by the *PNCK* gene**

Pregnancy up-regulated non-ubiquitous calmodulin kinase PNCK is a calcium/calmodulin-dependent protein kinase belonging to a calcium-triggered signaling cascade. It phosphorylates and activates CAMK1 that, upon calcium influx, regulates transcription activators activity, cell cycle, hormone production and cell differentiation.

Several lines of evidence suggest that PNCK promotes carcinogenesis.

PNCK has been found to be highly overexpressed in human primary human breast cancers compared with benign mammary tissue (Gardner *et al*., 2000). Increased expression of PNCK is associated with poor prognosis in clear cell renal cell carcinoma. The mRNA level of PNCK was significantly higher in tumorous tissues than in the adjacent non-tumorous tissues.

Multivariate analysis indicated that PNCK expression was an independent predictor for poor survival of clear cell renal cell carcinoma patients (Wu *et al*., 2013). Overexpression of PNCK in breast cancer cells was shown to result in increased proliferation, clonal growth and cell-cycle progression (Deb *et al*., 2015).

Recent studies have shown that *PNCK* depletion inhibits proliferation and induces apoptosis of human nasopharyngeal carcinoma cells *in vitro* and *in vivo*, suggesting it might be a novel therapeutic target for treatment of nasopharyngeal carcinoma (Xu *et al*., 2019).

Deb TB, Zuo AH, Barndt RJ, Sengupta S, Jankovic R, Johnson MD. Pnck overexpression in HER-2 gene-amplified breast cancer causes Trastuzumab resistance through a paradoxical PTEN-mediated process. Breast Cancer Res Treat. 2015; 150:347-361.

Gardner HP, Ha SI, Reynolds C, Chodosh LA. The caM kinase, Pnck, is spatially and temporally regulated during murine mammary gland development and may identify an epithelial cell subtype involved in breast cancer. Cancer Res. 2000; 60:5571-5577.

Wu S, Lv Z, Wang Y, Sun L, Jiang Z, Xu C, Zhao J, Sun X, Li X, Hu L, Tang A, Gui Y, Zhou F, et al. Increased expression of pregnancy up-regulated non-ubiquitous calmodulin kinase is associated with poor prognosis in clear cell renal cell carcinoma. PLoS One. 2013; 8:e59936.

Xu Y, Wang J, Cai S, Chen G, Xiao N, Fu Y, Chen Q, Qiu S. PNCK depletion inhibits proliferation and induces apoptosis of human nasopharyngeal carcinoma cells in vitro and in vivo. J Cancer. 2019;10:6925-6932..

#### Runt-related transcription factor 2, encoded by the *RUNX2* gene

The protein is a member of the RUNX family of transcription factors and has a Runt DNA-binding domain. RUNX2 is a transcription factor involved in osteoblastic differentiation and skeletal morphogenesis. RUNX2 plays a cell proliferation regulatory role in cell cycle entry and exit in osteoblasts. These functions are especially important when discussing bone cancer, particularly osteosarcoma development that can be attributed to aberrant cell proliferation control.

Several studies indicate that RUNX2 plays a key role in carcinogenseis. RUNX2 overexpression was found to promote aggressiveness and metastatic spreading, whereas *RUNX2* knockdown inhibits tumor growth and metastasis suggesting an oncogenic role for the protein (Tandon *et al*., 2014, Tandon *et al*., 2016, Shin *et al*., 2016, Li *et al*., 2016, Wang *et al*., 2016, Sancisi *et al*., 2017, Lu *et al*., 2018, Ji *et al*., 2019, Herreño *et al*., 2019).

Although strong purifying selection would not contradict the tumor promoting role of RUNX2, the high silent to missense ratio of substitution mutations is not a reflection of the strength of negative selection of missense and nonsense substitutions.

A noteworthy feature of the *RUNX2* gene is that its translated region contains a long stretch of CAG repeats (encoding the polyQ segment of the protein, residues 49-71). Interestingly, substitutions are not randomly distributed along the sequence of *RUNX2*: they are clustered in the polyQ tract of the protein encoded by the imperfect polymorphic CAG microsatellite region of the *RUNX2* gene. Since in cancer cells defective in mismatch-repair, microsatellites are known to become unstable due to increased frequency of replication error (Benachenhou, Labuda and Sinnett,1998), it seems likely that this increases and distorts mutation pattern in the polyQ region of *RUNX2*, and this mutation hotspot may give the false impression of strong purifying selection.

Benachenhou N, Labuda D, Sinnett D. Allelic instability of TBP gene in replication error positive tumors. Int J Cancer. 1998; 78:525-526.

Herreño AM, Ramírez AC, Chaparro VP, Fernandez MJ, Cañas A, Morantes CF, Moreno OM, Brugés RE, Mejía JA, Bustos FJ, Montecino M, Rojas AP. Role of RUNX2 transcription factor in epithelial mesenchymal transition in non-small cell lung cancer lung cancer: Epigenetic control of the RUNX2 P1 promoter. Tumour Biol. 2019; 41:1010428319851014.

Ji Q, Cai G, Liu X, Zhang Y, Wang Y, Zhou L, Sui H, Li Q. MALAT1 regulates the transcriptional and translational levels of proto-oncogene RUNX2 in colorectal cancer metastasis. Cell Death Dis. 2019;10:378.

Li XQ, Lu JT, Tan CC, Wang QS, Feng YM.RUNX2 promotes breast cancer bone metastasis by increasing integrin α5-mediated colonization. Cancer Lett. 2016; 380:78-86.

Lu H, Jiang T, Ren K, Li ZL, Ren J, Wu G, Han X. RUNX2 Plays An Oncogenic Role in Esophageal Carcinoma by Activating the PI3K/AKT and ERK Signaling Pathways. Cell Physiol Biochem. 2018; 49:217-225.

Sancisi V, Manzotti G, Gugnoni M, Rossi T, Gandolfi G, Gobbi G, Torricelli F, Catellani F, Faria do Valle I, Remondini D, Castellani G, Ragazzi M, Piana S, Ciarrocchi A. RUNX2 expression in thyroid and breast cancer requires the cooperation of three non-redundant enhancers under the control of BRD4 and c-JUN. Nucleic Acids Res. 2017; 45:11249-11267.

Shin MH, He Y, Marrogi E, Piperdi S, Ren L, Khanna C, Gorlick R, Liu C, Huang J. A RUNX2-Mediated Epigenetic Regulation of the Survival of p53 Defective Cancer Cells. PLoS Genet. 2016; 12:e1005884.

Tandon M, Chen Z, Othman AH, Pratap J. Role of Runx2 in IGF-1Rβ/Akt- and AMPK/Erk-dependent growth, survival and sensitivity towards metformin in breast cancer bone metastasis. Oncogene. 2016; 35:4730-4740.

Tandon M, Chen Z, Pratap J. Runx2 activates PI3K/Akt signaling via mTORC2 regulation in invasive breast cancer cells. Breast Cancer Res. 2014; 16:R16.

Wang X, Li L, Wu Y, Zhang R, Zhang M, Liao D, Wang G, Qin G, Xu RH, Kang T.CBX4 Suppresses Metastasis via Recruitment of HDAC3 to the Runx2 Promoter in Colorectal Carcinoma. Cancer Res. 2016; 76:7277-7289.

#### Monocarboxylate transporter 4 (MCT 4), encoded by the *SLC16A3* gene

Monocarboxylate transporter 4 (MCT4) or Solute carrier family 16 member 3 (SLC16A3) is a member of the proton-linked monocarboxylate transporter. Protein family. It catalyzes the rapid transport across the plasma membrane of many monocarboxylates such as lactate.

Since due to abnormal conversion of pyruvic acid to lactic acid by tumor cells even under normoxia, the altered metabolism of glucose consuming tumors must rapidly efflux lactic acid to the microenvironment to maintain a robust glycolytic flux and to prevent poisoning themselves (Mathupala et al., 2007). Survival and maintenance of the glycolytic phenotype of tumor cells is ensured by monocarboxylate transporter 4 (MCT4, encoded by the *SLC16A3* gene) that efficiently transports L-lactate out of the cell (Ganapathy, Thangaraju and Prasad, 2009).

As high metabolic and proliferative rates in cancer cells lead to production of large amounts of lactate, extruding transporters are essential for the survival of cancer cells. This point may be illustrated by the fact that knockdown of MCT4 increased tumor-free survival and decreased in vitro proliferation rate of tumor cells (Andersen *et al*., 2018).

Using a functional screen Baenke *et al*., (2015) have also demonstrated that monocarboxylate transporter 4 is an important regulator of breast cancer cell survival: MCT4 depletion reduced the ability of breast cancer cells to grow, suggesting that it might be a valuable therapeutic target.

In harmony with the essentiality of MCT4 for tumor growth, several studies indicate that expression of the hypoxia-inducible monocarboxylate transporter MCT4 is increased in tumors and its expression correlates with clinical outcome, thus it may serve as a valuable prognostic factor (Witkiewicz *et al*., 2012, Doyen *et al*., 2014, Baek *et al*., 2014)

Consistent with the key importance of MCT4 for the survival of tumor cells, its selective inhibition to block lactic acid efflux appears to be a promising therapeutic strategy against highly glycolytic malignant tumors (Todenhöfer *et al.*, 2018, Choi *et al*., 2016, 2018, Zhao *et al*., 2019)


Andersen AP, Samsøe-Petersen J, Oernbo EK, Boedtkjer E, Moreira JMA, Kveiborg M, Pedersen SF. The net acid extruders NHE1, NBCn1 and MCT4 promote mammary tumor growth through distinct but overlapping mechanisms. Int J Cancer. 2018; 142:2529-2542.

Baek G, Tse YF, Hu Z, Cox D, Buboltz N, McCue P, Yeo CJ, White MA, DeBerardinis RJ, Knudsen ES, Witkiewicz AK. MCT4 defines a glycolytic subtype of pancreatic cancer with poor prognosis and unique metabolic dependencies. Cell Rep. 2014; 9:2233-2249.

Baenke F, Dubuis S, Brault C, Weigelt B, Dankworth B, Griffiths B, Jiang M, Mackay A, Saunders B, Spencer-Dene B, Ros S, Stamp G, Reis-Filho JS, et al. Functional screening identifies MCT4 as a key regulator of breast cancer cell metabolism and survival. J Pathol. 2015; 237:152-165.

Choi SY, Xue H, Wu R, Fazli L, Lin D, Collins CC, Gleave ME, Gout PW, Wang Y. The *MCT4* Gene: A Novel, Potential Target for Therapy of Advanced Prostate Cancer. Clin Cancer Res. 2016;22:2721-2733.

Choi SYC, Ettinger SL, Lin D, Xue H, Ci X, Nabavi N, Bell RH, Mo F, Gout PW, Fleshner NE, Gleave ME, Collins CC, Wang Y. Targeting MCT4 to reduce lactic acid secretion and glycolysis for treatment of neuroendocrine prostate cancer. Cancer Med. 2018. 7:3385–3392.

Doyen J, Trastour C, Ettore F, Peyrottes I, Toussant N, Gal J, Ilc K, Roux D, Parks SK, Ferrero JM, Pouysségur J. Expression of the hypoxia-inducible monocarboxylate transporter MCT4 is increased in triple negative breast cancer and correlates independently with clinical outcome. Biochem Biophys Res Commun. 2014; 451:54-61.

Ganapathy V, Thangaraju M, Prasad PD. Nutrient transporters in cancer: relevance to Warburg hypothesis and beyond. Pharmacol Ther. 2009;121:29-40.

Mathupala SP, Colen CB, Parajuli P, Sloan AE. Lactate and malignant tumors: a therapeutic target at the end stage of glycolysis. J. Bioenerg. Biomembr. 2007; 39:73–77.

Todenhöfer T, Seiler R, Stewart C, Moskalev I, Gao J, Ladhar S, Kamjabi A, Al Nakouzi N, Hayashi T, Choi S, Wang Y, Frees S, Daugaard M et al. Selective Inhibition of the Lactate Transporter MCT4 Reduces Growth of Invasive Bladder Cancer Mol Cancer Ther. 2018; 17:2746-2755.

Witkiewicz AK, Whitaker-Menezes D, Dasgupta A, Philp NJ, Lin Z, Gandara R, Sneddon S, Martinez-Outschoorn UE, Sotgia F, Lisanti MP. Using the "reverse Warburg effect" to identify high-risk breast cancer patients: stromal MCT4 predicts poor clinical outcome in triple-negative breast cancers. Cell Cycle. 2012; 11:1108-1117.

Zhao Y, Li W, Li M, Hu Y, Zhang H, Song G, Yang L, Cai K, Luo Z. Targeted inhibition of MCT4 disrupts intracellular pH homeostasis and confers self-regulated apoptosis on hepatocellular carcinoma. Exp Cell Res. 2019; 31:111591.


#### Solute carrier family 2, facilitated glucose transporter member 1, encoded by the *SLC2A*1 gene

SLC2A1 functions as a facilitative glucose transporter, which is responsible for glucose uptake.

Significantly, several nutrient transporter protein genes were found among the genes showing the strongest signs of purifying selection. The most likely explanation for the selective pressure to preserve their integrity is that tumor cells have an increased demand for nutrients and this demand is met by enhanced cellular entry of nutrients through upregulation of specific transporters (Ganapathy, Thangaraju and Prasad, 2009).

The uncontrolled cell proliferation of tumor cells involves not only deregulated control of cell proliferation but also major adjustments of energy metabolism in order to fuel cell growth and division in the hypoxic microenvironments in which they reside. Otto Warburg was the first to observe an anomalous characteristic of cancer cell energy metabolism: even in the presence of oxygen, cancer cells limit their energy metabolism largely to glycolysis, leading to a state that has been termed "aerobic glycolysis (Warburg, 1956). Cancer cells are known to compensate for the lower efficiency of ATP production through glycolysis than oxidative phosphorylation by upregulating glucose transporters, such as GLUT1, thus increasing glucose import into the cytoplasm (Jones and Thompson, 2009, DeBerardinis *et al*., 2008, Hsu and Sabatini, 2008).

The markedly increased uptake of glucose has been documented in many human tumor types, by noninvasively visualizing glucose uptake through positron emission tomography using a radiolabeled analog of glucose as a reporter**.** This reliance of tumor cells on glycolysis is also supported by the hypoxia response system: under hypoxic conditions not only glucose transporters but also multiple enzymes of the glycolytic pathway are upregulated (Jones and Thompson, 2009, DeBerardinis *et al*., 2008. Semenza, 2010a,b, Kroemer and Pouyssegur, 2008)

In our view, the central role of GLUT1 in cancer metabolism is reflected by the fact that the gene (*SLC2A1* gene of solute carrier family member 2 protein) encoding this glucose transporter is among the genes that show the strongest signatures of purifying selection (see **Supplementary file 6**).

The key importance of GLUT1 in cancer may be illustrated by the fact that high levels of GLUT1 expression correlates with a poor overall survival and is associated with increased malignant potential, invasiveness and poor prognosis (Wang *et al*., 2017, Deng *et al*., 2018, de Castro *et al*., 2018).

The strict requirement for GLUT1 in the early stages of mammary tumorigenesis highlights the potential for glucose restriction as a breast cancer preventive strategy (Wellberg *et al*., 2016). The tumor essentiality of GLUT1 may also be illustrated by the fact that knockdown of GLUT1 inhibits cell glycolysis and proliferation and inhibits the growth of tumors (Xiao *et al*., 2018). In view of its essentiality for tumor growth, GLUT1 is a promising target for cancer therapy (Shibuya *et al*., 2015, Noguchi *et al*., 2016, Chen *et al*., 2017)

Recent studies suggest that the YAP1-TEAD1-GLUT1 axis plays a major role in reprogramming of cancer energy metabolism by modulating glycolysis (Lin and Xu, 2017). These authors have shown that YAP1 and TEAD1 are involved in transcriptional control of the the glucose transporter GLUT1: whereas. knockdown of YAP1 inhibited glucose consumption, and lactate production of breast cancer cells. overexpression of GLUT1 restored glucose consumption and lactate production.


Chen Q, Meng YQ, Xu XF, Gu J. Blockade of GLUT1 by WZB117 resensitizes breast cancer cells to adriamycin. Anticancer Drugs. 2017; 28:880-887.

DeBerardinis RJ, Lum JJ, Hatzivassiliou G. Thompson C.B. The biology of cancer: Metabolic reprogramming fuels cell growth and proliferation. Cell Metab. 2008; 7:11-20

de Castro TB, Mota AL, Bordin-Junior NA, Neto DS, Zuccari DAPC. Immunohistochemical Expression of Melatonin Receptor MT1 and Glucose Transporter GLUT1 in Human Breast Cancer. Anticancer Agents Med Chem. 2018; 18:2110-2116.

Deng Y, Zou J, Deng T, Liu J. Clinicopathological and prognostic significance of GLUT1 in breast cancer: A meta-analysis. Medicine (Baltimore). 2018; 97:e12961.

Ganapathy V, Thangaraju M, Prasad PD. Nutrient transporters in cancer: relevance to Warburg hypothesis and beyond. Pharmacol Ther. 2009; 121:29-40.

Hsu PP, Sabatini DM. Cancer cell metabolism: Warburg and beyond. Cell. 2008; 134:703-707

Jones RG. Thompson C.B. Tumor suppressors and cell metabolism: a recipe for cancer growth. Genes Dev. 2009; 23:537-548

Kroemer G. Pouyssegur J. Tumor cell metabolism: Cancer's Achilles' heel. Cancer Cell. 2008; 13:472-482

Lin C, Xu X. YAP1-TEAD1-Glut1 axis dictates the oncogenic phenotypes of breast cancer cells by modulating glycolysis. Biomed Pharmacother. 2017; 95:789-794.

Noguchi C, Kamitori K, Hossain A, Hoshikawa H, Katagi A, Dong Y, Sui L, Tokuda M, Yamaguchi F. D-Allose Inhibits Cancer Cell Growth by Reducing GLUT1 Expression. Tohoku J Exp Med. 2016; 238:131-141.

Semenza GL. Defining the role of hypoxia-inducible factor 1 in cancer biology and therapeutics. Oncogene. 2010; 29:625-634

Semenza GL. HIF-1: upstream and downstream of cancer metabolism. Curr. Opin. Genet. Dev. 2010; 20:51-56

Shibuya K, Okada M, Suzuki S, Seino M, Seino S, Takeda H, Kitanaka C. Targeting the facilitative glucose transporter GLUT1 inhibits the self-renewal and tumor-initiating capacity of cancer stem cells. Oncotarget. 2015; 6:651-661.

Wang J, Ye C, Chen C, Xiong H, Xie B, Zhou J, Chen Y, Zheng S, Wang L. Glucose transporter GLUT1 expression and clinical outcome in solid tumors: a systematic review and meta-analysis.
Oncotarget. 2017; 8:16875-16886.

Warburg O. On the origin of cancer cells. Science. 1956;123:309-314.

Wellberg EA, Johnson S, Finlay-Schultz J, Lewis AS, Terrell KL, Sartorius CA, Abel ED, Muller WJ, Anderson SM. The glucose transporter GLUT1 is required for ErbB2-induced mammary tumorigenesis. Breast Cancer Res. 2016; 18:131.

Xiao H, Wang J, Yan W, Cui Y, Chen Z, Gao X, Wen X, Chen J. GLUT1 regulates cell glycolysis and proliferation in prostate cancer. Prostate. 2018; 78:86-94.

#### Solute carrier family 2, facilitated glucose transporter member 8, encoded by the *SLC2A8* gene

The SLC2A8/GLUT8 is a member of the glucose transporter superfamily that mediates the transport of glucose and fructose.

In harmony with the strong signatures of negative selection there is evidence that GLUT8 plays an important role in carcinogenesis: it is overexpressed in and is required for proliferation and viability of tumors (Goldman *et al*., 2006, McBrayer *et al*., 2012).

Goldman NA, Katz EB, Glenn AS, Weldon RH, Jones JG, Lynch U, Fezzari MJ, Runowicz CD, Goldberg GL, Charron MJ. GLUT1 and GLUT8 in endometrium and endometrial adenocarcinoma. Mod Pathol. 2006; 19:1429-1436.

McBrayer SK, Cheng JC, Singhal S, Krett NL, Rosen ST, Shanmugam M. Multiple myeloma exhibits novel dependence on GLUT4, GLUT8, and GLUT11: implications for glucose transporter-directed therapy Blood. 2012; 119:4686-4697.

#### TATA-box-binding protein, encoded by the *TBP* gene

The *TBP* gene has been selected as a gene showing very high values of rSMN, suggesting negative selection of missense and nonsense mutations (**Supplementary file 8**). It must be pointed out that based on the high silent/missense ratio *TBP* (as well as *DSPP* and *MLLT3*) has also been identified by others as a gene subject to negative selection (Zhou *et al*., 2017).

The protein is a general transcription factor that functions at the core of the DNA-binding multiprotein factor TFIID. Binding of TFIID to the TATA box is the initial transcriptional step of the pre-initiation complex, playing a role in the activation of eukaryotic genes transcribed by RNA polymerase II. In view of such a basic cell essential function, it seemed justified to assume that it is the indispensability of the gene for the survival of tumor cells (just like any other cell) that subjects it to strong purifying selection and the high silent/missense ratio is a reflection of this negative selection. TBP has been thought to be an invariant housekeeping protein, however, several studies have shown that TBP expression is significantly increased in both colon adenocarcinomas as well as adenomas relative to normal tissue, supporting the idea that increases in TBP expression actually drive tumorigenesis (Johnson *et al*., 2003a,b, Johnson *et al*., 2017).

Inspection of the spectrum of somatic mutations of the *TBP* gene suggests that the high silent/missense ratio is unlikely to be simply due to negative selection that may hold for both oncogenes and tumor essential genes. A noteworthy feature of the *TBP* gene is that its translated region contains a long stretch of CAG repeats (encoding the polyQ segment of the protein, residues 57-95). The distribution of silent mutations is markedly non-random: they are clustered in the polyQ tract of the protein encoded by the imperfect polymorphic CAG microsatellite region of the *TBP* gene. Since in cancer cells defective in mismatch-repair, microsatellites are known to become unstable due to increased frequency of replication error (Benachenhou, Labuda and Sinnett, 1998), it seems likely that this is why the rate of mutation in the polyQ region of TBP is much higher than in other regions of the gene. The high silent to missense rate is thus not due to negative selection acting on missense and nonsense substitutions. Rather, it may reflect the fact that the imperfect polymorphic CAG microsatellite region of the *TBP* gene serves as a mutation hotspot, with a biased substitution pattern.

Benachenhou N, Labuda D, Sinnett D. Allelic instability of TBP gene in replication error positive tumors. Int J Cancer. 1998; 78:525-526.

Johnson SA, Dubeau L, Kawalek M, Dervan A, Schönthal AH, Dang CV, Johnson DL. Increased expression of TATA-binding protein, the central transcription factor, can contribute to oncogenesis. Mol Cell Biol. 2003; 23:3043-3051.

Johnson SA, Dubeau L, White RJ, Johnson DL. The TATA-binding protein as a regulator of cellular transformation. Cell Cycle. 2003; 2:442-444.

Johnson SAS, Lin JJ, Walkey CJ, Leathers MP, Coarfa C, Johnson DL. Elevated TATA-binding protein expression drives vascular endothelial growth factor expression in colon cancer. Oncotarget. 2017; 8:48832-48845.

Zhou Z, Zou Y, Liu G, Zhou J, Wu J, Zhao S, Su Z, Gu X. Mutation-profile-based methods for understanding selection forces in cancer somatic mutations: a comparative analysis. Oncotarget. 2017;8:58835-58846

#### Thromboxane A2 receptor, encoded by the *TBXA2R* gene

TBXA2R is a plasma membrane protein that serves as a receptor for thromboxane A2, a potent stimulator of platelet aggregation. The activity of this receptor is mediated by a G-protein that activates a phosphatidylinositol-calcium second messenger system.

Studies on the expression of thromboxane A2 receptor, TBXA2R in a cohort of human breast cancer patients revealed that breast tumour tissues expressed higher levels of TBXA2R compared with normal mammary tissues and that TBXA2R expression was most significantly increased in grade 3 tumours. Kaplan-Meier survival analysis has also shown that patients with high levels of TBXA2R had significantly shorter disease-free survival. The observation that TBXA2R is highly expressed in aggressive tumours and linked with poor prognosis indicates that TBXA2R has a significant prognostic value in clinical breast cancer (Watkins *et al*., 2005).

The role of TBXA2R in carcinogenesis is also supported by the observation that Thromboxane A2 was shown to enhance tumor metastasis and that the tumor promoting activity required intact TBXA2 receptor (Matsui *et al*., 2012). These studies revealed that TBXA2-TBXA2R signaling plays a critical role in tumor colonization through P-selectin-mediated interactions between platelets-tumor cells and tumor cells-endothelial cells, suggesting that blockade of this signaling might be useful in the treatment of tumor metastasis.

Although the involvement of TBXA2-TBXA2R signaling in cancer invasion and metastasis apperars to be clearly established**,** there may be other mechanisms by which TBXA2 promotes these processes. Li *et al.* (2013) have shown that a TBXA2 mimetic induced the expression of the monocyte chemoattractant chemokine ligand protein CCL2, suggesting that TBXA2 may also stimulate invasion of cancer cells through CCL2-CCR2 mediated macrophage recruitment.

Recent studies on Triple Negative Breast Cancer (TNBC) cell lines revealed that TBXA2R expression was higher in these cell lines and that *TBXA2R* knockdowns consistently showed dramatic cell killing in TNBC cells (Orr *et al*., 2016). It has also been shown that TBXA2R enhanced TNBC cell migration, invasion, indicating that the gene is required for the survival and migratory behaviour of a subset of TNBCs.

A phenome-wide association study has shown that a single nucleotide polymorphism in the gene *TBXA2R* is associated with increased metastasis in multiple primary cancers, suggesting the requirements for thromboxane A2 (TXA2) and TBXA2R in the basic mechanism of metastasis, and the clinical applicability of TBXA2R antagonists as adjuvant therapy in multiple cancers (Pulley *et al*., 2018).

Li X, Tai HH. Activation of thromboxane A2 receptor (TP) increases the expression of monocyte chemoattractant protein -1 (MCP-1)/chemokine (C-C motif) ligand 2 (CCL2) and recruits macrophages to promote invasion of lung cancer cells. PLoS One. 2013; 8:e54073.

Matsui Y, Amano H, Ito Y, Eshima K, Suzuki T, Ogawa F, Iyoda A, Satoh Y, Kato S, Nakamura M, Kitasato H, Narumiya S, Majima M. Thromboxane $A_2$ receptor signaling facilitates tumor colonization through P-selectin-mediated interaction of tumor cells with platelets and endothelial cells. Cancer Sci. 2012; 103:700-707.

Orr K, Buckley NE, Haddock P, James C, Parent JL, McQuaid S, Mullan PB. Thromboxane A2 receptor (TBXA2R) is a potent survival factor for triple negative breast cancers (TNBCs). Oncotarget. 2016; 7:55458-55472.

Pulley JM, Jerome RN, Ogletree ML, Bernard GR, Lavieri RR, Zaleski NM, Hong CC, Shirey-Rice JK, Arteaga CL, Mayer IA, Holroyd KJ, Cook RS. Motivation for Launching a Cancer Metastasis Inhibition (CMI) Program. Target Oncol. 2018; 13:61-68.

Watkins G, Douglas-Jones A, Mansel RE, Jiang WG. Expression of thromboxane synthase, TBXAS1 and the thromboxane A2 receptor, TBXA2R, in human breast cancer. Int Semin Surg Oncol. 2005; 2:23.

#### Tumor protein p73, encoded by the *TP73* gene

The protein is known to participate in the apoptotic response to DNA damage: isoforms containing the N-terminal transactivation domain are pro-apoptotic, isoforms lacking the transactivation domain are anti-apoptotic.

Although p73 shows substantial homology with p53, despite the established role of p53 as a tumor suppressor, p73 does not have a similar tumor suppressor role in malignancy: unlike p53-/- mice, p73 knockout mice do not develop tumors. In fact, N-terminally truncated p73 isoforms, lacking the transactivation domain were shown to possess oncogenic potential (Stiewe and Pützer, 2002, Stiewe *et al*., 2002).

Numerous studies have shown that ΔNp73, the oncogenic isoform of p73 lacking the transactivation domain, is frequently up-regulated in many carcinomas and is indicative of poor prognosis (Zaika *et al*., 2002, Petrenko, Zaika and Moll, 2003, Domínguez *et al*., 2006, Hassan *et al*., 2014, Hassan, Dave and Singh, 2014, Lucena-Araujo *et al*., 2015).

Our observation that p73, an oncogenic protein, shows only strong signatures of purifying selection provides one of the clearest examples illustrating the point that in the case of oncogenes purifying selection is not necessarily associated with positive selection for driver mutations. It must be pointed out here that it has been noted earlier by others that, despite its clear role in carcinogenesis, the *TP73* gene is almost never mutated (Bisso, Collavin and Del Sal, 2011, Maas *et al*., 2013). One may argue that in this case the molecular change that drives carcinogenesis is the change of splicing that favors the formation of the oncogenic isoform of p73.

Bisso A, Collavin L, Del Sal G. p73 as a pharmaceutical target for cancer therapy. Curr Pharm Des. 2011; 17:578-590.

Domínguez G, García JM, Peña C, Silva J, García V, Martínez L, Maximiano C, Gómez ME, Rivera JA, García-Andrade C, Bonilla F. DeltaTAp73 upregulation correlates with poor prognosis in human tumors: putative in vivo network involving p73 isoforms, p53, and E2F-1. J Clin Oncol. 2006; 24:805-815.

Hassan HM, Dave BJ, Singh RK1. TP73, an under-appreciated player in non-Hodgkin lymphoma pathogenesis and management. Curr Mol Med. 2014 May;14(4):432-9.

Hassan HM, Varney ML, Jain S, Weisenburger DD, Singh RK, Dave BJ. Disruption of chromosomal locus 1p36 differentially modulates TAp73 and ΔNp73 expression in follicular lymphoma. Leuk Lymphoma. 2014; 55:2924-2931.

Lucena-Araujo AR, Kim HT, Thomé C, Jacomo RH, Melo RA, Bittencourt R, Pasquini R, Pagnano K, Glória AB, Chauffaille Mde L, Athayde M, Chiattone CS, Mito I, et al. High ΔNp73/TAp73 ratio is associated with poor prognosis in acute promyelocytic leukemia. Blood. 2015; 126:2302-2306.

Maas AM, Bretz AC, Mack E, Stiewe T. Targeting p73 in cancer. Cancer Lett. 2013; 332:229-236.

Petrenko O, Zaika A, Moll UM. deltaNp73 facilitates cell immortalization and cooperates with oncogenic Ras in cellular transformation in vivo. Mol Cell Biol. 2003; 23:5540-5555.

Stiewe T, Pützer BM. Role of p73 in malignancy: tumor suppressor or oncogene? Cell Death Differ. 2002; 9:237-245.

Stiewe T, Zimmermann S, Frilling A, Esche H, Pützer BM. Transactivation-deficient DeltaTA-p73 acts as an oncogene. Cancer Res. 2002; 62:3598-3602.

Zaika AI1, Slade N, Erster SH, Sansome C, Joseph TW, Pearl M, Chalas E, Moll UM. DeltaNp73, a dominant-negative inhibitor of wild-type p53 and TAp73, is up-regulated in human tumors. J Exp Med. 2002; 196:765-780.

#### Tribbles homolog 2, encoded by the *TRIB2* gene

TRIB2 is a pseudokinase member of the pseudoenzyme class of signaling/scaffold proteins. It interacts with MAPK kinases and regulates activation of MAP kinases.

TRIB2 has been shown to be important in the maintenance of the oncogenic properties of melanoma cells, as its silencing reduces cell proliferation, colony formation. Tumor growth was also substantially reduced upon RNAi-mediated TRIB2 knockdown in an *in vivo* melanoma xenograft model, suggesting that TRIB2 provides the melanoma cells with growth and survival advantages (Zanella *et al*., 2010).

TRIB2 expression is elevated in primary human lung tumors and in non-small cell lung cancer cells, resulting from gene amplification. *TRIB2* knockdown was found to inhibit cell proliferation and *in vivo* tumor growth, indicating that TRIB2 is a potential driver of lung tumorigenesis (Grandinetti *et al*., 2011).

High TRIB2 expression is observed in T cell acute lymphoblastic leukaemias (Hannon *et al*., 2012). TRIB2 has been shown to be critical for both solid and non-solid malignancies and is functionally important for liver cancer cell survival and transformation. TRIB2 was found to be up-regulated in liver cancer cells compared with other cells (Wang *et al*., 2013 a,b).

TRIB2 is emerging as a pivotal target of transcription factors in acute leukemias as evidenced by the fact *TRIB2* knockdown resulted in a block in acute myeloid leukemia cell proliferation (Rishi *et al*., 2014)

In the case of lung adenocarcinoma, patients with higher TRIB2 levels had poorer survival (Zhang *et al*., 2016). The tumor promoting role of this protein is supported by the observation that TRIB2 expression is significantly increased in tumour tissues from patients with extremely poor clinical outcome (Hill *et al*., 2017, Wang *et al*., 2019).

TRIB2 has been shown to be important for the survival of leukemia cells during MLL-TET1-related leukemogenesis and for maintaining differentiation blockade of leukemic cells: *TRIB2* knockdown relieved the inhibition of myeloid cell differentiation induced by the MLL-TET1 fusion protein (Kim *et al*., 2018).

TRIB2 expression has been shown to be elevated in colorectal cancer tissues compared to normal adjacent tissues and high TRIB2 expression indicated poor prognosis of colorectal cancer patients (Hou *et al*., 2018). Depletion of TRIB2 inhibited cancer cell proliferation, induced cell cycle arrest and promoted cellular senescence, whereas overexpression of TRIB2 accelerated cell growth, cell cycle progression and blocked cellular senescence.

Grandinetti KB, Stevens TA, Ha S, Salamone RJ, Walker JR, Zhang J, Agarwalla S, Tenen DG, Peters EC, Reddy VA. Overexpression of TRIB2 in human lung cancers contributes to tumorigenesis through downregulation of C/EBPα. Oncogene. 2011; 30:3328-3335.

Hannon MM, Lohan F, Erbilgin Y, Sayitoglu M, O'Hagan K, Mills K, Ozbek U, Keeshan K. Elevated TRIB2 with NOTCH1 activation in paediatric/adult T-ALL. Br J Haematol. 2012; 158:626-634.

Hill R, Madureira PA, Ferreira B, Baptista I, Machado S, Colaço L, Dos Santos M, Liu N, Dopazo A, Ugurel S, Adrienn A, Kiss-Toth E, Isbilen M, et al. TRIB2 confers resistance to anti-cancer therapy by activating the serine/threonine protein kinase AKT. Nature Communications. 2017; 8:14687.

Hou Z, Guo K, Sun X, Hu F, Chen Q, Luo X, Wang G, Hu J, Sun L. TRIB2 functions as novel oncogene in colorectal cancer by blocking cellular senescence through AP4/p21 signaling. Mol Cancer. 2018;17:172.

Kim HS, Oh SH, Kim JH, Sohn WJ, Kim JY, Kim DH, Choi SU, Park KM, Ryoo ZY, Park TS, Lee SJ. TRIB2 regulates the differentiation of MLL-TET1 transduced myeloid progenitor cells.
Mol Med (Berl). 2018; 96:1267-1277

Rishi L, Hannon M, Salomè M, Hasemann M, Frank AK, Campos J, Timoney J, O'Connor C, Cahill MR, Porse B, Keeshan K. Regulation of Trib2 by an E2F1-C/EBPα feedback loop in AML cell proliferation. Blood. 2014; 123:2389-2400.

Wang J, Park JS, Wei Y, Rajurkar M, Cotton JL, Fan Q, Lewis BC, Ji H, Mao J. TRIB2 acts downstream of Wnt/TCF in liver cancer cells to regulate YAP and C/EBPα function. Mol Cell. 2013; 51:211-225.

Wang J, Zhang Y, Weng W, Qiao Y, Ma L, Xiao W, Yu Y, Pan Q, Sun F. Impaired phosphorylation and ubiquitination by p70 S6 kinase (p70S6K) and Smad ubiquitination regulatory factor 1 (Smurf1) promote tribbles homolog 2 (TRIB2) stability and carcinogenic property in liver cancer. J Biol Chem. 2013; 288:33667-33681**.**

Wang J, Zuo J, Wahafu A, Wang MD, Li RC, Xie WF. Combined elevation of TRIB2 and MAP3K1 indicates poor prognosis and chemoresistance to temozolomide in glioblastoma. CNS Neurosci Ther. 2019 Jul 18.

Zanella F, Renner O, García B, Callejas S, Dopazo A, Peregrina S, Carnero A, Link W. Human TRIB2 is a repressor of FOXO that contributes to the malignant phenotype of melanoma cells. Oncogene. 2010; 29:2973-2982.

Zhang YX, Yan YF, Liu YM, Li YJ, Zhang HH, Pang M, Hu JX, Zhao W, Xie N, Zhou L, Wang PY, Xie SY. Smad3-related miRNAs regulated oncogenic TRIB2 promoter activity to effectively suppress lung adenocarcinoma growth. Cell Death Dis. 2016; 7:e2528.

#### Twist-related protein 1, encoded by the *TWIST1* gene

The *TWIST1* gene is characterized by very high value of rSMN (**Supplementary file 3**), indicating strong signature of purifying selection, suggesting that it plays an important role in promoting tumorigenesis.

Twist-related protein 1, TWIST1 is a transcription factor and master regulator of the epithelial-to-mesenchymal transition that significantly contributes to tumor growth and metastasis. TWIST1 is overexpressed in a variety of tumors and numerous studies have shown that targeting TWIST1 significantly inhibits tumor growth (Wushou *et al*., 2014, Zhu *et al*., 2016, Xu *et al*., 2017a,b, Mikheev *et al*., 2018).

Recent studies have revealed that AURKA and TWIST1 are linked in as much as ablation of either AURKA or TWIST1 completely inhibits epithelial-to-mesenchymal transition (Wang *et al*., 2017).

Mikheev AM, Mikheeva SA, Severs LJ, Funk CC, Huang L, McFaline-Figueroa JL, Schwensen J, Trapnell C, Price ND, Wong S, Rostomily RC. Targeting TWIST1 through loss of function inhibits tumorigenicity of human glioblastoma. Mol Oncol. 2018; 12:1188-1202.

Wang J, Nikhil K, Viccaro K, Chang L, Jacobsen M, Sandusky G, Shah K. The Aurora-A-Twist1 axis promotes highly aggressive phenotypes in pancreatic carcinoma. J Cell Sci. 2017; 130:1078-1093.

Wushou A, Hou J, Zhao YJ, Shao ZM. Twist-1 up-regulation in carcinoma correlates to poor survival. Int J Mol Sci. 2014; 15:21621-21630.

Xu Y, Lee DK, Feng Z, Xu Y, Bu W, Li Y, Liao L, Xu J. Breast tumor cell-specific knockout of Twist1 inhibits cancer cell plasticity, dissemination, and lung metastasis in mice. Proc Natl Acad Sci U S A. 2017; 114:11494-11499

Xu Y, Qin L, Sun T, Wu H, He T, Yang Z, Mo Q, Liao L, Xu J. Twist1 promotes breast cancer invasion and metastasis by silencing Foxa1 expression. Oncogene. 2017; 36:1157-1166.

Zhu QQ, Ma C, Wang Q, Song Y, Lv T. The role of TWIST1 in epithelial-mesenchymal transition and cancers. Tumour Biol. 2016; 37:185-197.
